# The Bacterial Kinase AnmK Integrates into Tick Genome and Biology

**DOI:** 10.64898/2026.03.13.711553

**Authors:** Alexandra Cerqueira de Araujo, Ondřej Hajdušek, Veronika Urbanová, Tereza Šedivá, Luise Robbertse, Alexander Titz, Olivier Plantard, Yannis Stahl, Christoph Mayer, Jan Perner, Claude Rispe

## Abstract

Horizontal Gene Transfer (HGT) contributes to eukaryotic evolution, potentially bringing phenotypic novelties to the recipient organisms. Ticks pose a severe threat to human health as vectors of various pathogens, including viruses, bacteria and protozoa, while stably hosting endosymbiotic bacteria. As such, these obligate blood-feeding parasites have been and continue to be exposed to HGT from a broad range of donors. To determine whether bacterial-to-tick HGT has contributed to important tick traits, we surveyed *Ixodes* tick genomes for HGT events. We revealed duplications of the known bacteria-derived gene *dae2* and discovered two novel cases of bacterial HGT, the most remarkable of which involves a bacterial peptidoglycan metabolic gene (*anmK*) acquired by the common ancestor of ticks. The acquisition of an intron demonstrates “eukaryotization” of *anmK* within tick genomes. Transcript profiling revealed that *anmK* expression is upregulated during blood feeding, peaking in female ovaries, a niche occupied by horizontally acquired endosymbionts. Biochemical analysis confirmed that, to some extent, recombinant AnmK retains kinase activity on its cognate substrate – the bacterial cell wall component 1,6-anhydro-N-acetylmuramic acid. Immunolocalization showed that the enzyme is predominantly localized towards outer layers of the vitellogenic oocytes. Silencing of *anmK* in different tick species compromised blood-feeding and reproduction, demonstrating that this domesticated bacterial enzyme underpins reproductive fitness across tick species. Our findings exemplify the ability of horizontally acquired genes to integrate into the host biology and thereby shape host life history.

## 1. Introduction

The normal inheritance route for genetic material is vertical, from parents to offspring of the same species. Yet occasional transfer of genetic elements from one species to another can occur in a process known as horizontal gene transfer (HGT). Massive horizontal transfers occurred in the early stages of the evolution of eukaryotes, following their association with bacteria that subsequently evolved into organelles, such as mitochondria and chloroplasts. During this process, hundreds of bacterial genes moved to host genomes. In more recent evolutionary times, spanning the last hundreds of million years, such transfers have been relatively rare in most organisms, with rotifers standing out among metazoans for their high rate of recent HGT ^1^.

Although recent HGT events constitute only a small fraction of the genome of most eukaryotes, they can have significant evolutionary and adaptive consequences. Horizontally acquired genes in eukaryotes can originate from various sources, including viruses, fungi, and bacteria. For example, aphids acquired the ability to synthesize carotenoids from fungi via HGT ^2^, and numerous cases of bacterial-to-eukaryote transfers have been documented (reviewed in ^3^). Notable cases of HGT from viruses to eukaryotes include the protein syncytin, which facilitated the evolution of the placenta in mammals ^4,5^, and genes that enable the production of viral-like particles, which enhanced parasitism success in parasitoid wasps ^6–8^. Mobile genetic elements, or transposable elements (TEs), can also shape eukaryotic genomes by integrating into host genomes and introducing new genetic material. In some cases, horizontal transfer allows TEs to “infect” new species, while providing a source of evolutionary change in the receiving organisms ^9^.

HGT from bacteria has also conferred novel metabolic and ecological capabilities to recipient organisms. For instance, plant-feeding nematodes have acquired bacterial genes that allow them to degrade plant carbohydrates (notably genes of the Glycoside Hydrolase Family 32, or GH32), facilitating infestation of roots ^10^. Additionally, bacterial genes encoding bactericidal toxins active against competitors have been horizontally transferred to eukaryotes, contributing to antimicrobial defense. Interestingly, several cases of HGT concern genes involved in the metabolism of peptidoglycan (PG), a fundamental component of the bacterial cell envelope. The acquired genes participate in either the degradation or the synthesis of PG. The principal role of PG is osmoprotection, along with defining the shape of the bacterial cell and providing bacteria with protection from environmental assaults ^11^. Intracellular bacterial symbionts face specific challenges, including host immune pressure and highly specialized metabolic environments, which profoundly affect their PG biosynthetic pathways. Accordingly, many endosymbiotic bacteria have undergone partial loss of PG biosynthesis genes^12^. A striking example of PG-related HGT is found in mealybugs, which harbor two bacterial endosymbionts, *Moranella* and *Tremblaya*. In this system, PG synthesis is functionally maintained by ten bacterial genes that were horizontally transferred from multiple bacterial clades into the mealybug genome, forming a mosaic biosynthetic pathway^3^. This example highlights how HGT can preserve essential metabolic functions in symbiotic systems while enabling host-level control and regulation.

Ticks, members of the Chelicerata subphylum, are obligate blood-feeding ectoparasites, their group comprising over 900 described species. They play a significant role in the transmission of various pathogens, including bacteria, viruses, and protozoa, posing substantial threats to both human and animal health^13^. Ticks combine prolonged blood feeding on diverse vertebrate hosts with extended off-host development in microbe-rich environments such as soil and leaf litter, while permanently harbouring bacterial symbionts. These ecological and biological interactions place ticks at the interface of multiple potential gene pools, facilitating exposure to HGT from a broad range of donors, including bacteria (pathogenic, symbiotic, and commensal), viruses, and even vertebrate hosts^14^.

Several cases of HGT have already been documented in ticks, with the majority involving viral elements ^15–18^. Some of these endogenous viral elements appear to contribute to immune responses, as in the case of RNA interference (RNAi)-mediated antiviral defense^16,19^. Beyond viruses, HGT from vertebrate donors has also been reported in ticks, including the retrotransposon Bov-B in hard ticks ^20^, the tick adrenomedullin (*TAM*) in soft ticks ^14^, and an interferon-binding motif of mammalian origin integrated in the gene *Dome1*, which triggers the JAK–STAT pathway in *Ixodes scapularis* ^21^. In the cases of *TAM* and *Dome1*, however, the very short domains concerned and their very low conservation between tick and mammalian sequences suggest a possible scenario of convergence rather than proper HGT. To date, the sole reported case of functional HGT from bacteria to ticks involves a horizontally acquired gene in Acari, a group that includes ticks and mites ^22^. This gene, named *domesticated amidase effector 2* (*dae2*), encodes a secreted protein that hydrolyzes PG, thereby exerting antibacterial activity. The protein is present in the tick gut and saliva and RNA interference experiments showed that *dae2* prevents bacterial overgrowth during blood feeding ^22,23^. This suggests that bacterial HGT can contribute to the immune defense of ticks by targeting bacterial cell walls. More recently, a study identified fragments of genomic DNA from bacterial symbionts, (namely *Spiroplasma* and *Midichloria* spp.) integrated into the genome of the tick *Ixodes ricinus*, most likely representing degraded and non-functional genes, based on their short sequences and low RNA-Seq support ^24^.

We hypothesized that ticks’ exposure to bacteria of both types, pathogens and symbionts, could have favored events of horizontal transfer along the evolutionary history of the group. Even rare HGT events could have been evolutionarily meaningful if the transferred genes persisted along tick evolution (co-diversifying with the host genome) and became integrated into host biology. Moreover, they could have enabled ticks to interact with other bacteria they are exposed to during their life-cycle, as has been proposed for *dae2*. To test whether bacterial-to-tick HGT has contributed to important tick traits, we performed a stringent genome-wide screen and then examined the phylogenetic distribution and expression of high-confidence candidates across diverse tick representatives. This approach revealed two previously unrecognized bacterial acquisitions, with *anmK* - a PG-recycling kinase-standing out as a broadly retained, tick-encoded bacterial gene that is consistently associated with ovarian biology. Because tick ovarian symbionts vary in their retention of PG pathway genes, including frequent loss or pseudogenization, the discovery of a conserved, host-encoded *anmK* raises testable hypotheses about domestication of bacterial cell-wall metabolism in the context of symbiont coexistence. We therefore combined comparative genomics with expression and functional analyses to determine how this horizontally acquired gene has been assimilated into tick reproductive biology.

## 2. Results

### 2.1 Tick genomes have integrated several genes of viral and bacterial origin

To identify and quantify the extent of HGT events between ticks and other organisms, we analysed the genome of *I. ricinus* to detect tick genes of alien origin, with possible donors including vertebrates, plants, bacteria, viruses, or other non-arthropod taxa (**Fig. 1A**, **Fig. S1**) using the Alienness vs Predictor (AvP) software. In addition, a specific phylogenetic re-analysis was conducted for the *dae2* gene - which was not identified by AvP but was previously found to be of bacterial origin in the tick *I. scapularis*. The full list of HGT candidate genes identified for *I. ricinus* with both methods is given in **Table S1** and comprises 14 genes of viral origin, three genes of vertebrate origin (see below for criticism), and six genes of bacterial origin.

**Figure 1.**
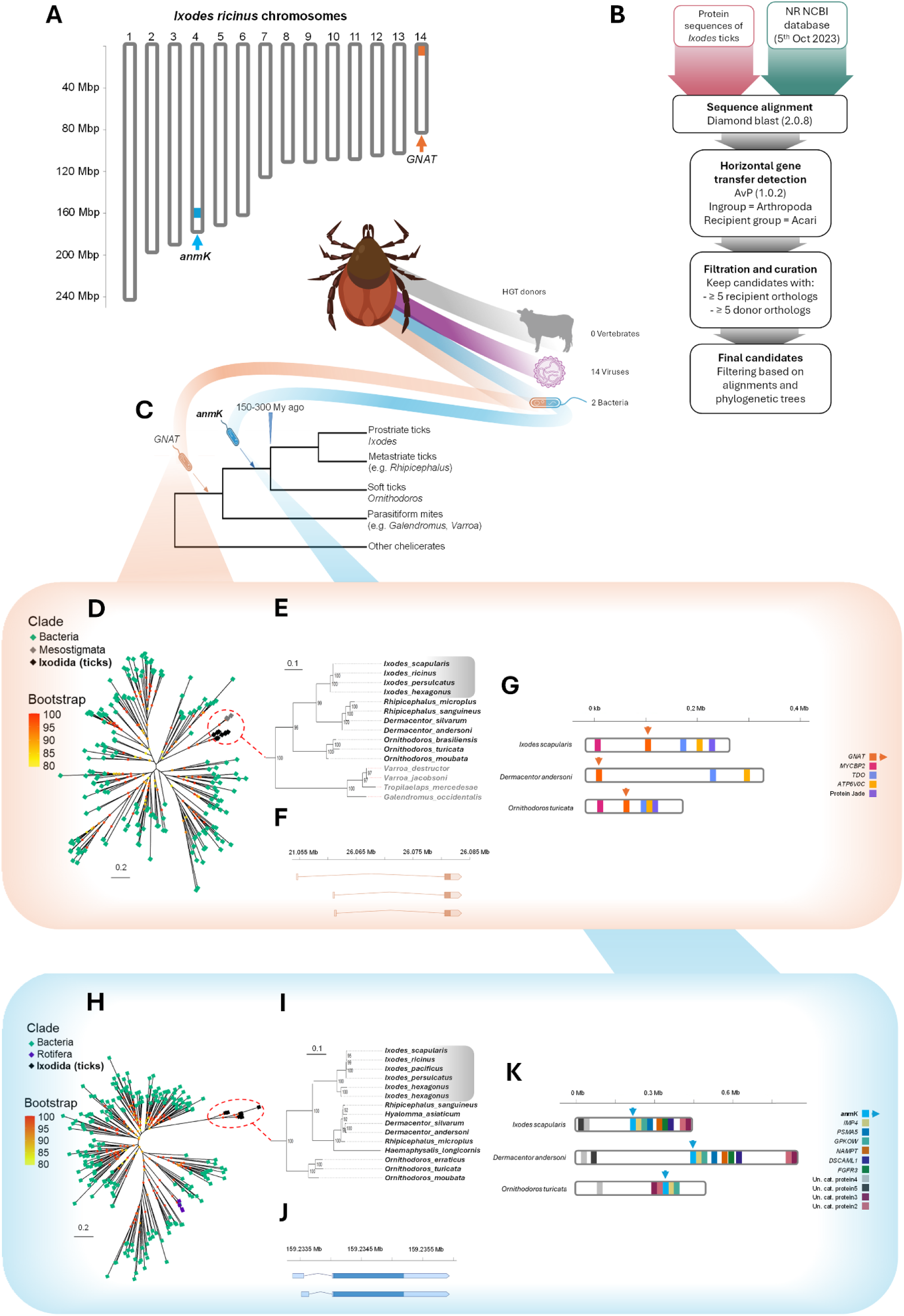
Two newly discovered cases of bacterial horizontal gene transfer in ticks. **A** The fourteen chromosomes of *I. ricinus* and the relative genomic positions of two genes horizontally transferred from bacteria (*anmK* in blue and *GNAT* in orange). **B** Summary of the HGT detection pipeline. Although three sequences from vertebrate donors were detected by the first steps of the pipeline, they were finally discarded after phylogenetic analysis. **C** Evolutionary timeline of the insertion of *anmK* and *GNAT* in tick genomes. The displayed date corresponds to the approximate date of the last common ancestor (LCA) of ticks. The acquisition of *anmK* occurred prior to the tick LCA, while the one of *GNAT* occurred prior to the LCA of ticks + Mesostigmata. **D,H** Maximum-Likelihood phylogenetic tree of *GNAT* (**D**) and *anmK* (**H**). Both genes are present in bacteria (in green), in ticks (in black), GNAT being additionally present in Mesostigmata (grey). Only closely related bacterial sequences are included, representing the potential donor group - *Ignavibacteriota* (**D**) or *Chitinophagaceae* (**H**). The *anmK* tree also shows an independent HGT from the same bacterial group to rotifers (in purple). **E**,**I** Focus on the phylogeny of *GNAT* (**E**) and *anmK* (**I**) in ticks (in black, *Ixodes* species are highlighted with a grey background) and other parasitiforms (Mesostigmata) in grey. **F,J** Both *GNAT* (**F**) and *anmK* (**J**) in *I. ricinus* have acquired an intron and have alternative transcripts. **G,K** Genomic environment of *GNAT* (**G**) and *anmK* (**K**). Microsynteny between tick genomes further supports a unique ancestral HGT event, for both genes.

#### Detection of viral genes integrated in the tick genome (*I. ricinus*)

The 14 genes annotated as *I. ricinus* coding genes and identified as of viral origin could be assigned to five groups (**Fig. S2A to E**): Phlebovirus-like, Nairovirus-like, Totivirus-like, Densovirus-like, and Rhabdovirus-like. All these genes except one exhibited extremely low expression levels based on our RNA-Seq expression atlas (**Fig. S3A**, and data publicly available from the Gene Note Book at https://bipaa.genouest.org/sp/ixodes_ricinus/). However, IricT00012932, a gene with a sequence similar to nairoviruses, was well expressed, particularly in the eggs and in the synganglion. Of note, we found four of these endogenous viral elements (EVEs) form a physical cluster, although they each represent three different groups of viruses: IricT00015847 and IricT00015848 (rhabdoviruses), IricT00015852 (nairovirus) and IricT00015853 (phlebovirus). These sequences were indeed localized within a relatively narrow 900 kb region, with no other coding gene in this interval.

#### No evidence for transfer from vertebrate hosts to the tick genome

Whereas three candidate genes were identified by AvP as potential HGT candidates from vertebrate donors, a more in-depth phylogenetic analysis found no conclusive support for any of these genes being horizontally transferred (**Fig. S4A to C**).

#### Re-analysis of the horizontally transferred toxin dae2 in ticks

We reanalyzed the phylogenetic framework of the transferred bacterial enzyme toxin *tae2* in ticks (renamed *dae2* in *I. scapularis*^22^), discovering multiple copies of this gene. We indeed found that *I. scapularis* possesses a second copy, which we named *dae2-like* (**Fig. S5**). Both *dae2* and *dae2-like* have two closely related co-orthologs in *I. ricinus*, resulting in four genes for this species (*dae2-1* to *4*). RNA-Seq data showed that the most highly expressed copy of *dae2* in *I. ricinus* is *dae2-1*, its expression peaking in hemocytes, while *dae2-3* and dae2-4 showed highest expression in the synganglion (**Fig. S3C**).

#### Phylogenetic evidence for the transfer of an acyltransferase (GNAT) to ticks and Mesotigma

A gene found in tick genomes (named IricT00008184 in *I. ricinus*) was detected as a horizontally transferred bacterial gene by AvP (AI threshold =20). This gene, showing similarity to GCN5-related N-acetyltransferases, was named *GNAT*. Our phylogenetic analysis confirmed that this sequence exists as a single-copy gene in every tick species studied, including hard ticks and soft ticks of the genus *Ornithodoros* (**Fig. 1C**). Additionally, a similar sequence was found in several species of Mesostigmata (which, like ticks, belong to the superorder Parasitiformes), but not in Acariformes, nor in any other Arthropoda group. All blastP hits of these genes were bacterial, which strongly supports that this gene originates from a bacterial donor. Our phylogenetic analysis supports a unique HGT event from a bacterial donor, likely belonging to the Ignavibacteriota phylum to a common ancestor of ticks and Mesostigmata (**Fig. 1D,E**). In Mesostigmata, this gene has several coding exons, while in ticks, it has a single coding exon with one additional 5’UTR exon (the intronic region is supported by mapped RNA-Seq reads for several tick species). The ancestral bacterial gene has thus acquired introns following its introgression, a sign of domestication in a eukaryotic context that may facilitate gene expression and regulation by the eukaryotic machinery ^25^ (**Fig. 1F**). The expression profile based on RNA-Seq data indicates that tick *GNAT* is expressed predominantly in the synganglion of adult ticks during feeding, and to a lesser degree, in the synganglion of unfed ticks and in eggs, whereas the expression is very low in other tissues (**Fig. S3B**).

#### Phylogenetic evidence for the transfer of *anmK* to the common ancestor of ticks

A second case of bacterial HGT concerned a gene named IricT00013998 in the *I. ricinus* genome, which we found to be homologous to *anmK,* a gene encoding a 1,6 anhydro-N-acetylmuramic acid kinase. Among all potential transfer events detected by AvP, this case was the most robustly supported, as it was identified using the most stringent threshold (AI=40, **Table S1**). This gene is widely present in bacteria and normally absent in Eukaryota, yet we found it in every tick genome analysed. AnmK is an enzyme involved in the recycling metabolism of bacterial cell walls, and intervenes in the re-synthesis of peptidoglycan (PG). We found that the gene is present both in hard ticks and soft ticks (*Ornithodoros* spp.) and localizes in a genomic region with conserved synteny in ticks, especially hard ticks (see **Fig. 1K**). The phylogenetic tree of tick *anmK* sequences reflects the evolutionary history of ticks, which is consistent with a single horizontal transfer event that occurred in the common ancestor of ticks, with Chitinophagacea as the bacterial donor group (all bacterial sequences in our phylogeny belonged to this group, as they were the most closely related sequences detected by blast) (**Fig. 1H**). Our phylogeny shows that an independent *anmK* HGT event also occurred in rotifers, from the same bacterial donor group (**Fig. 1H**). Of note, HGT events have shown to be relatively common in rotifers ^1^. RNA-Seq data for *I. ricinus* suggests that *anmK* is well expressed in ticks, with a moderate baseline expression in several tissues and a strong peak of expression in ovaries of adult females during feeding (**Fig. S3B**).

A targeted search for *anmK* in tick-associated bacterial symbionts failed to identify homologs in *Coxiella*-like endosymbionts and in *Rickettsia* tick symbionts, suggesting that the gene is absent from both groups. In contrast, *anmK* was detected in *Midichloria* (associated with *I. ricinus*) and in *Francisella*-like endosymbionts. However, the tick *anmK* sequence was not closely related to those found in tick symbionts. Instead, it robustly formed a clade with sequences from the Chitinophagaceae. Notably, Chitinophagaceae are represented by two distinct clades of *anmK*, which we refer to as clade 1 and clade 2 (**Fig. S6**). The tick *anmK* sequence grouped with clade 2, which is more divergent from other bacterial *anmK* sequences than the less divergent clade 1. Notable substitutions in the tick sequence (for example a substitution of asparagine to cysteine at the site N173) and some bacterial sequences of this clade 2 could be linked with an evolution of function.

Codon usage analysis showed that both *GNAT* and *anmK* have a codon usage typical of the *I. ricinus* genome (**Fig. S7A**) and thus do not display signatures of recently transferred or contaminating DNA. This fits with an ancient transfer and subsequent codon adaptation of both genes. Notably, *anmK* has a codon profile relatively close to ribosomal proteins, a group of genes expected to display an optimal codon usage (**Fig. S7A), B**). This may reflect the fact that this gene is highly expressed at least at one point of the life-cycle, and is thus under a selective pressure for efficient translation. A codon usage analysis comparing the genomes of ticks (*I. ricinus*) and a bacterial species representing the donor group (*Terrimonas* spp.) further confirmed the marked difference in codon usage between tick and bacterial a*nmK* (**Fig. S7C**), which is a signature of its domestication in the tick genome.

Overall, this HGT event is particularly intriguing because the gene encodes a function that is strictly bacterial in origin and because it is expressed at a critical stage and tissue of the tick life cycle, namely in the ovaries of fed ticks. For these reasons, we investigated the potential function of the tick *anmK* homolog in *Ixodes ricinus*, hereafter referred to as *Iranmk*.

### 2.2 Intron acquisition has ‘eukaryotized’ the tick *anmK* gene, with ovary-enriched expression across species

Based on the annotation in the *I. ricinus* genome, *anmK*, like *GNAT*, has a single coding exon, combined with a 5’UTR exon (**Fig. 1J**): RNA-Seq data lend strong support to the presence of an intron, with two possible splice sites after the first exon (one of the alternative transcripts being largely dominant). Therefore, an intron was acquired along *anmK* evolution, an additional mark of eukaryotic domestication ^25^. This was further confirmed by amplifying the region spanning the *anmK* gene and an adjacent *bona fide* eukaryotic gene encoding the U3 small ribonucleoprotein (*IMP4*, **Fig. 2A**). The PCR amplicon (5590 bp) sequence validated the presence of a predicted intron in the 5’-UTR region of the *anmK* gene. To confirm that this intron is spliced out during mRNA maturation, we amplified both genomic DNA and cDNA templates. The cDNA amplicon was 463 bp smaller than the genomic DNA amplicon, corresponding to the intron size (**Fig. 2B**), while both products were sequence-verified, confirming a eukaryotic structure. Last, detection of *anmK* transcripts in oligo(dT)-enriched RNA (RNA-seq) and in oligo(dT)-primed cDNA synthesis indicates that the mRNA is polyadenylated, providing further evidence that this gene has been fully eukaryotized.

**Figure 2.**
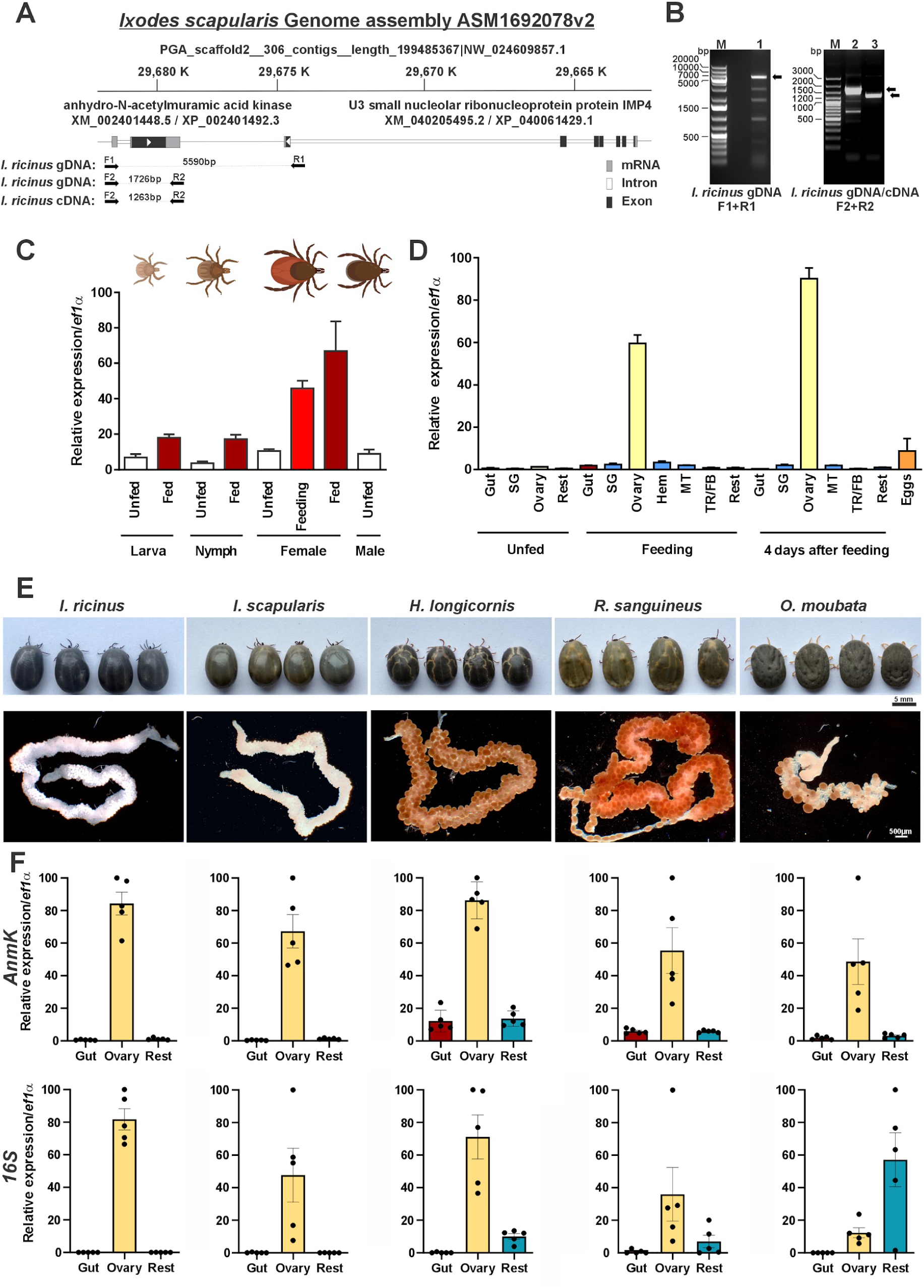
Gene structure and expression of *anmK* in ticks. **A** Localization of *anmK* in the genome of *I. scapularis* (source: NCBI, region around XM_002401448.5). Shown are also the positions of the *I. ricinus*-specific primers (F1 and R1) that were used to amplify a 5590 bp region of genomic DNA containing *anmK*, an intergenic region, and the terminal part of the neighboring *imp4* gene in the amplicon. Primers F2 and R2 were used to amplify the *anmK* gene from gDNA and cDNA of *I. ricinus* with different amplicon lengths (1726 and 1263bp), confirming the presence of a spliceable intron. **B** Agarose gel electrophoresis of PCR amplifications of the corresponding regions from gDNA template (Lane 1 and 2) and cDNA template (Lane 3). The arrows indicate the bands that were excised, cloned and sequenced. **C** RT-qPCR profiles of *IranmK* in *I. ricinus* whole bodies for different stages, sexes and conditions of fed (dark red), feeding (red) and unfed (white) ticks throughout development. Bars indicate mean ± SEM, n = 3. **D** RT-qPCR profiles of *IranmK* in different tissues of *I. ricinus* and under different conditions; SG-salivary glands, MT-Malphigian tubules, Hem-Hemocytes, TR/FB-Trachea, Fat body. Bars indicate mean ± SEM, n = 3. **E** Photographic images of fully engorged females of tick species used in this study, namely *I. ricinus*, *I. scapularis*, *H. longicornis*, *R. sanguineus*, and *O. moubata*. For each species, macroscopic images of ovaries of females dissected out at four days after full engorgement, and are shown below images of engorged tick females. **F** For each of the five tick species in (E), RT-qPCR profiles of *anmK* (top) and prokaryotic 16S mRNA copies (bottom) are shown, demonstrating the positive correlation between *anmK* mRNAs levels and the presence of endosymbiotic bacteria within the tick ovary. Bars indicate mean ± SEM, n = 5.

To examine the tick *anmK* expression pattern throughout the tick life-cycle, we constructed comprehensive cDNA libraries representing different developmental stages (larvae, nymphs and adults) and internal organs of *I. ricinus* ticks (**Fig. 2C-D**). The expression profiles revealed a clear pattern with elevated *IranmK* mRNA levels after blood-feeding at every developmental stage (**Fig. 2C**). Notably, the highest expression levels were observed in the ovaries of adult tick females, peaking at four days after blood-feeding (**Fig. 2D**).

To assess this trait across different tick species, we also examined *anmK* expression in *I. scapularis*, in two non-*Ixodes* (Metastriata) tick species, *Haemaphysalis longicornis* and *Rhipicephalus sanguineus*, and in one species of soft ticks, *Ornithodoros moubata*, thereby covering the major branches of the tick evolutionary tree. In engorged females of all tested species, *anmK* mRNA was specifically present in the ovaries compared to other internal organs (**Fig. 2E-F**), suggesting a localized biological role. Interestingly, tick ovaries are heavily colonized by endosymbiotic bacterial species (**Fig. 2F**), leading to the hypothesis that the AnmK protein may play a role in tick reproduction and possibly also at the tick-endosymbiont interface.

### 2.3. IrAnmK is located in the vicinity of the plasma membrane of female oocytes

To gain further insight into the putative function of anmK in ticks, we developed antibodies against recombinant IrAnmK expressed in *E. coli* (Fig. S9A, B, Fig. S10). The antibodies proved to be highly specific and revealed that the AnmK protein is indeed natively expressed in tick ovaries (Fig. 3A). The raised antibodies were labelled with Alexa 647 using NHS Ester coupling chemistry for immunolocalization experiments (Fig. S9B, C). Using sections of vitellogenic oocytes from females collected at two time-points after full engorgement (4 and 9 days), we observed that IrAnmK is predominantly localized towards the outer layers of the oocytes and later, also surrounding vitellogenic vesicles (Fig. 3B). A minor portion of the signal was also detected in the oocyte cytosol but did not overlap with *Midichloria mitochondrii*, the main endosymbionts of *I. ricinus*, which were visualized by extra-nuclear DAPI staining (Fig. 3B). We thus conclude that IrAnmK encoded in the tick genome is not being exploited by its endosymbiont *M. mitochondrii*, but participates in tick developmental and reproduction processes.

**Figure 3.**
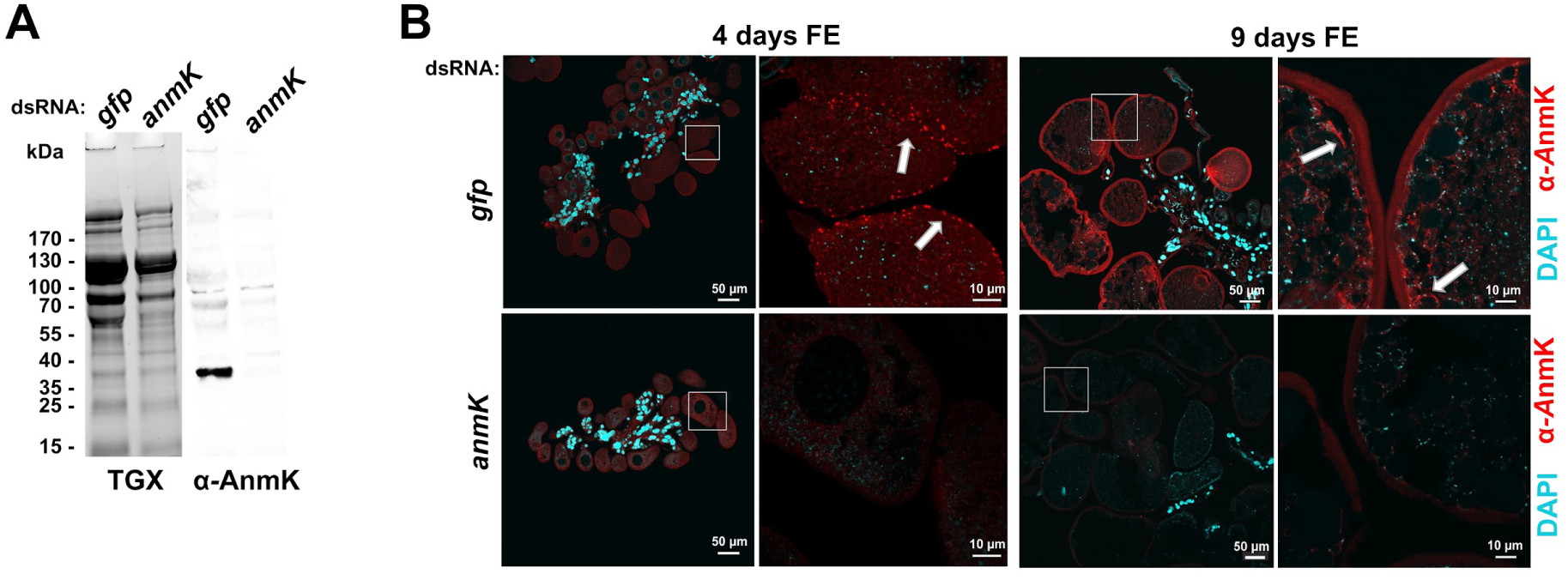
*IranmK* is expressed in tick oocytes and the protein localizes in its plasma membrane vicinity **A** SDS-PAGE (left) of a homogenate of ovaries of *I. ricinus* females four days after full engorgement. Western blotting (right) determines the specificity of raised IgY antibodies with clear signal in gfp control ovaries and absence in the ovaries homogenate from *IranmK*-KD ticks. **B** Immunolocalization of IrAnmK protein in sections of vitellogenic ovaries from *I. ricinus* females four and nine days after full engorgement (FE). Controls with identical antibodies applied to RNA-KD ovary sections confirmed the absence of specific signals. Nucleic acids were stained with DAPI (blue).

### 2.4. Tick AnmK reveals high structural identity with bacterial AnmKs and shows anhMurNAc kinase activity

To assess whether IrAnmK retained the overall structure as well as the active site of its bacterial AnmK counterparts, its 3D structure was predicted using AlphaFold3 ^26^ and compared with the published structure of *Pseudomonas aeruginosa* AnmK (PaAnmK) containing the non-hydrolyzable ATP analog AMPPNP and anhydroMurNAc bound to the active site (PDB: 8CPB ^27^). We additionally included in this analysis the AlphaFold3 structure predictions from EcAnmK, (AlphaFold Database; P77570) and from the putative AnmK of *Terrimonas pollutisoli* (TpAnmK), which belongs to the clade 2 of Chitinophagaceae AnmKs (cf. **Fig. S6**) and is phylogenetically closer to IrAnmK than EcAnmK and PaAnmK. A structure overlay is shown in **Fig. 4A** and shows a very similar overall structure and active site arrangement. Previous studies discovered the apo- and co-structure of bacterial AnmKs and identified the active site amino acid residues critical for substrate binding and catalysis ^27–29^. Accordingly, AnmK contains two subdomains, which are separated by a deep, open cleft, and follows a random-sequential kinetic mechanism. Upon sequential binding of both substrates, two protein loops (L8-14 and L229-241 in PaAnmK) move and generate a closed conformational state that facilitates catalysis. Critical amino acid residues include D14, which coordinates a magnesium ion and the alpha- and beta-phosphates of ATP, T97 and R129, which coordinate the N-acetyl and carboxylate groups of anhMurNAc, respectively, D182, which attacks the anomeric carbon of anhMurNAc, aiding hydrolysis of the anhydro-bond, and E326, which participates in binding of the gamma-phosphate of ATP and anhMurNAc, facilitating the phosphoryl transfer. These amino acids, critical for ATP and anhMurNAc binding (**Fig. 4A**) according to ^27,28^, were all retained in the IrAnmK structure. Nonetheless, subtle differences in residues comprising the active site were identified (**Fig. S6, S8**), most importantly a substitution from asparagine N73 (*E. coli* numbering) to cysteine in tick AnmKs.

**Figure 4.**
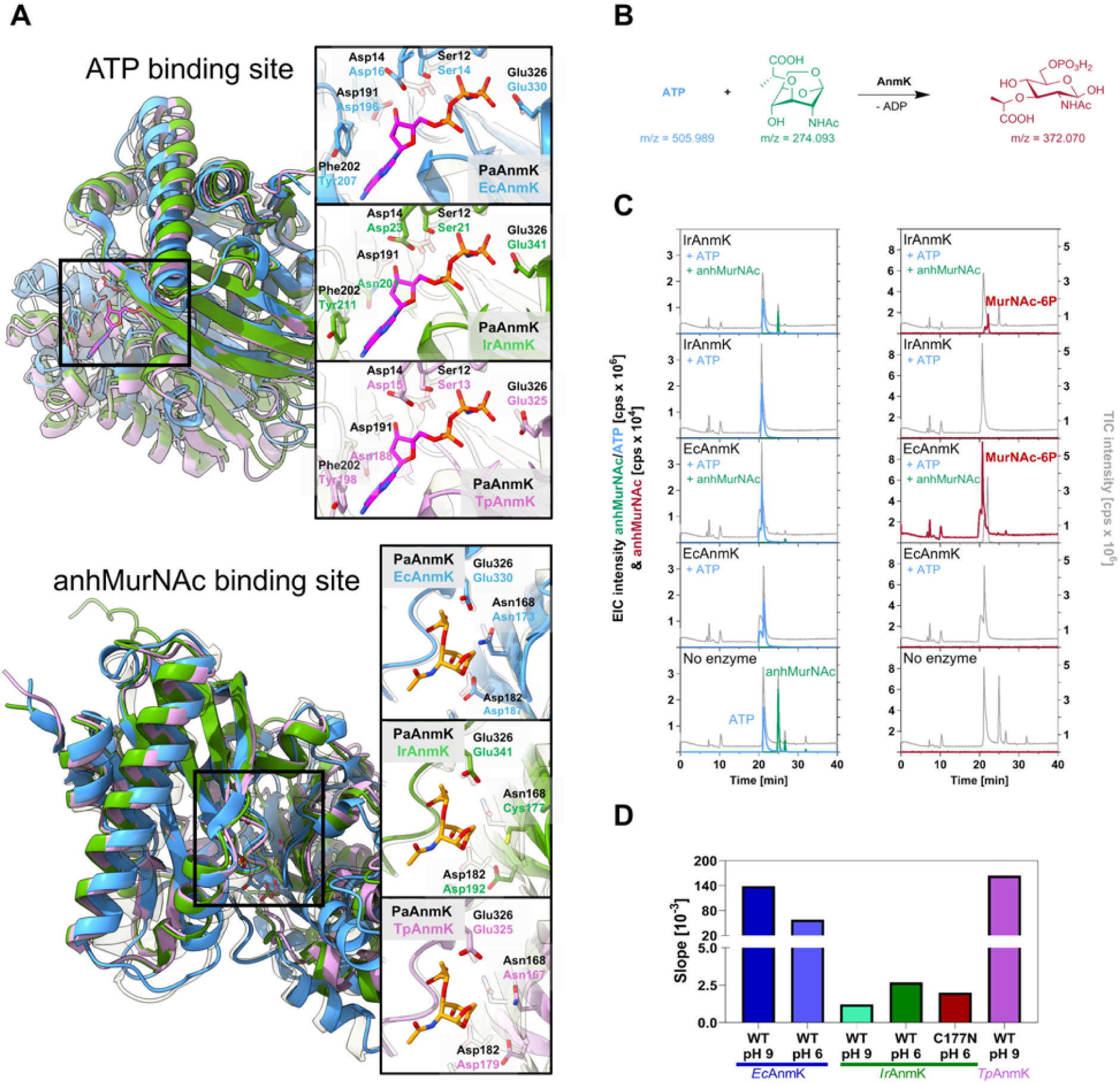
Structure comparison of tick and bacterial AnmK and analysis of their enzymatic activity. **A** Visualization of the 3D structures and structure predictions of AnmK proteins. Structural models of AnmK of *E. coli* (EcAnmK), *I. ricinus* (IrAnmK) and *T. pollutisoli* (TpAnmK) are shown in an overlap comparison with a published co-structure of *Pseudomonas aeruginosa* AnmK (PaAnmk) along with the non-hydrolyzable ATP analog AMPPNP and anhydroMurNAc (PDB: 8CPB; ^27^). For structure predictions of IrAnmK and TpAnmK AlphaFold 3 was used^26^ and the predicted structure of EcAnmK was retrieved from the AlphaFold Database (P77570). Insets on the overlay overall structures with PaAnmK highlight the substrate binding sites: AMPPNP (ATP) is depicted in magenta and anhydroMurNAc (anhMurNAc) in orange. Visualizations were prepared using ChimeraX 1.9. For the ATP binding sites the amino acids Phe202, Asp191, Asp14, Ser12, Glu326 of PaAnmK and their corresponding counterparts from the predicted structures are highlighted; likewise, for the anhMurNAc binding sites, Glu326, Asn168, Asp182 from PaAnmK and their corresponding counterparts are highlighted. All depicted amino acids are reported to be involved in substrate binding in PaAnmK ^27,29^. **B** Schematic of the AnmK reaction. The anhydro-ring of anhMurNAc is hydrolyzed by AnmK and concomitantly phosphorylated by ATP to yield the MurNAc-6-phosphate product. The exact masses of the substrates and the MurNAc-6-phosphate product are shown. **C** Mass spectrometric analysis of the AnmK activity of recombinant putative AnmK from *I. ricinus* (IrAnmK) and AnmK from *E. coli* (EcAnmK) incubated for 20 h in the presence or absence of 1,6-anhydro-N-acetylmuramic acid (anhMurNAc) and ATP. Results from HPLC-MS analysis before and after incubation are shown as the total-ion chromatograms (TIC; grey) and extracted-ion chromatograms (EIC) of anhydroMurNAc (red, [M-H^+^]^-^ = 274.0932 m/z), ATP (blue, [M-H^+^]^-^ = 505.9885 m/z) and MurNAc-6P (green, [M-H^+^]^-^ = 372.0701 m/z). As a control the reaction mixture without enzyme was analyzed. **C** The activity of AnmK homologs from *E. coli* (EcAnmK), *I. ricinus* (IrAnmK) and *T. pollutisoli* (TpAnmK) - a species within the bacterial donor group of *anmK* - was determined at pH 6 and 9 by calculating the reaction rate constants of linear regression equation (slope) (cf Fig. 11). For the IrAnmK variant (C177N), activity was measured only at pH 6. **D** For quantification of AnmK activity a continuous assay was applied that couples ADP production to the conversion of PEP to pyruvate and then lactate by the sequential activity of pyruvate kinase (PK) and lactate dehydrogenase (LDH) –which simultaneously oxidizes NADH to NAD+ resulting in a decrease of absorbance at 340 nm (cf. Suppl. **Fig. S11**).

To test whether recombinant IrAnmK retained anhMurNAc kinase activity, we incubated the protein with 1,6-anhydro-N-acetyl-muramic acid (anhMurNAc) and analyzed product formation via high performance liquid chromatography-mass spectrometry (HPLC-MS). A new compound with an m/z of 372.070, consistent with the mass of MurNAc6P, appeared when IrAnmK was incubated with anhMurNAc and ATP (**Fig. 4B,C**). This product was not detected in the absence of anhMurNAc or enzyme. As a positive control, an analysis of product formation using *E. coli* AnmK (EcAnmK) was conducted in parallel and yielded the same MurNAc6P product. Higher intensity of the product formed in the latter reaction indicated that EcAnmK is more active than IrAnmK under the reaction conditions that we used. To compare the relative rates of reactions of IrAnmK and EcAnmK with anhMurAc, a coupled enzyme assay was conducted that allowed to monitor the time course of the reaction and to calculate relative reaction rates (**Fig. S11**). In this assay, oxidation of NADH to NAD+ is used as a proxy for the formation of the second AnmK reaction product, ADP. The specific enzyme activity was determined from the linear decrease of NADH absorption (**Fig. S11)** and is shown as changes (slope) in differential absorption at 340 nm (Δ340 nm), which increases over time (**Fig. 4D**). Initial time course experiments were conducted at pH 9, since the activity of EcAnmK was reported to be high under alkaline conditions up to pH 10, but low at pH 6 ^30^. Interestingly, when conducting the assay at pH 6, the activity of IrAnmK appeared to be about twice as high as at pH 9, while the activity of the *E. coli* enzyme expectedly dropped (**Fig. 4D and Fig. S11**). Since the enzyme concentration of EcAnmK was 10-times lower than that of IrAnmK in these assays, the latter enzyme’s rate is considerably lower (c. 500-fold). Given that the pH dependence of the AnmK enzymes differs, it can be speculated that the optimal reaction conditions for IrAnmK have yet to be defined. It also cannot be excluded that anhMurNAc is not the favorable substrate for IrAnmK. Using the coupled enzyme assay, we tested the activity of recombinant AnmK from the bacterium *Terrimonas pollutisoli* (TpAnmK), which is phylogenetically closer to IrAnmK than EcAnmK and belongs to the clade 2 of Chitinophagaceae AnmKs (cf. **Fig. S6**). TpAnmK showed rates similar to EcAnmK at pH 9 (**Fig. 4D** and **Fig. S11**), indicating that TpAnmK is a *bona fide* anhMurNAc kinase. Therefore, the reduced activity of IrAnmK is not an ancestral feature shared with the donor group, i.e. bacterial sequences from the clade 2 of Chitinophagaceae AnmKs. As mentioned above, the N173C exchange is the most striking particularity of tick AnmK (**Fig. 4A**). Notably, an asparagin is retained in the fully active TpAnmK at this site (**Fig. S8**). We speculated that the revertant mutation C177N (*I. ricinus* numbering) could render the enzyme more active, but this was not the case (**Fig.4D**). The activity of the IrAnmK_C177N variant was slightly lower than the WT IrAnmK (measured at pH 6), and did not reach the levels of EcAnmK or TpAnmK activities. In conclusion, IrAnmK exhibited distinct anhMurNAc kinase activity, but this activity is significantly lower than that of bacterial proteins and shows a different pH-dependency compared to EcAnmK. These findings suggest a possible change in substrate specificity following horizontal transfer to ticks.

### 2.5. RNA-silencing of *IranmK* compromises the development of the tick ovary

To further investigate the function of *IranmK*, we conducted RNAi-mediated silencing in blood-feeding females (**Fig. 5A**). The efficacy of *IranmK* silencing by RNAi in tick ovaries was 98% (**Fig. 5B**). Importantly, the significant RNA-silencing effect was long-lasting (as long as 6 months), with maintained reduced *Ir-aAnmK* mRNA levels also in eggs and larvae (**Fig. 5A,B**), confirming a robust RNAi model to study the impact of silencing not only on blood-feeding, but also on development, reproduction, and biology of larvae (**Fig. 5B**). The *IranmK*-KD females appeared to remain fully competent in blood-feeding on guinea pigs, reaching comparable engorged weights to controls, indicating no essential function of *IranmK* during blood-feeding of *I. ricinus* adult females (**Fig. 5C**). Upon full engorgement of tick females, haemolipoproteins (vitellogenins and carrier proteins) are transported to the ovary, leading to deposition of maternal haem and facilitating egg development and growth^31^. After *IranmK* silencing, the ovary failed to incorporate haemolymphatic lipoproteins, as apparent by reduced protein content and maternal haem (red color), lagging behind in development of this organ compared to controls (**Fig. 5D-G**). Western blotting confirmed that ovaries of *IranmK*-KD females are specifically depleted of haemolipoproteins vitellins (proteolytic products of vitellogenins - Vg) and carrier protein 3 (CP3) (**Fig. 5H**). However, both weights of laid clutches (**Fig. 5I**) and of hatched larvae (**Fig. 5J, K**) were identical to mock (*gfp*) controls. Larvae that hatched from these females displayed significantly lower success in blood-feeding on their hosts (**Fig. 5L**). The metabolic activity of the *I. ricinus* symbiont *M. mitochondrii* was apparently unaffected in *IranmK*-KD ticks, as shown by *gyrB* mRNA levels measured by RT-qPCR (**Fig. 5M**). Similarly, the expression level of *IranmK* was independent of the presence/absence of the *M. mitochondrii* endosymbiont, as shown by comparison between a wild type and a *Midichloria*-free *I. ricinus* strain (**Fig. S12B**). Finally, injection of model microbes in unfed ticks did not alter the expression levels of *IranmK* (**Fig. S12C**). Overall, dsRNA-mediated silencing revealed an important role for IrAnmK in ovarian development in *Ixodes ricinus*, and thus in tick reproductive fitness and the blood-feeding capacity of their offspring, although this was not reflected in some metrics measured in lab conditions (e.g. clutch weight). In conclusion, while the co-occurrence of IrAnmK and of the *I. ricinus* symbiont *M. mitochondrii* in this organ suggested a potential interaction between them, the existence and exact nature of this interaction remain to be determined.

**Figure 5.**
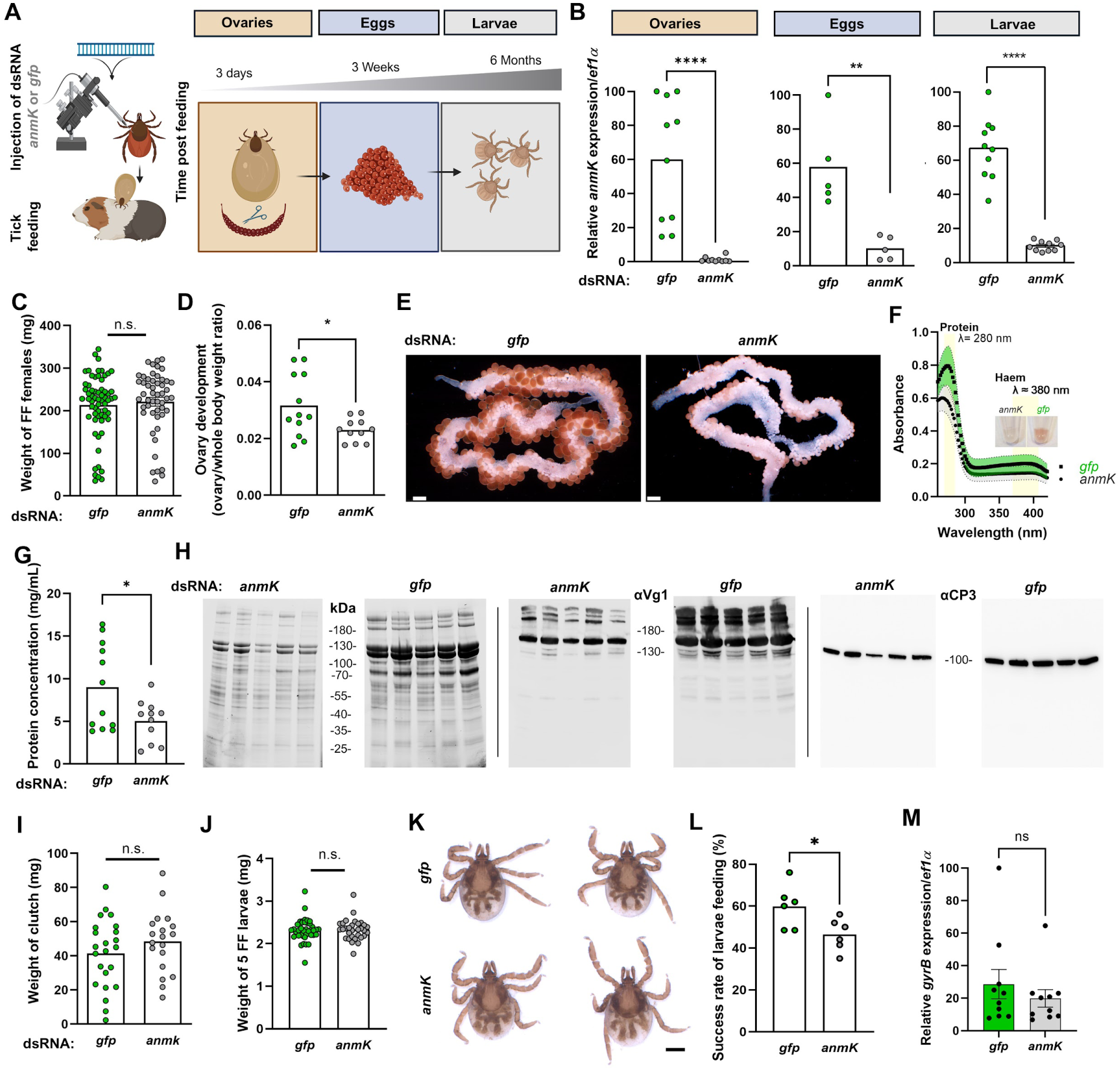
RNA-silencing of *IrAmK* transcript impairs development of tick ovaries and affects the blood-feeding competence of hatched larvae. **A** Schematic overview of the RNAi experiment setting used in this study, with *I. ricinus* females being injected dsRNA before and after blood-feeding. **B** RT-qPCR assessment of RNAi efficiency in ovaries, eggs, and larvae of *I. ricinus* females - each data point represents an independent biological replicate. **C** The bar graph depicts the weights of spontaneously detached fully engorged (FE) *I. ricinus* females, n ≥ 50. **D** Ratio of weights of ovaries to whole body weight of fully fed females six days after detachment, n ≥ 10. **E** Macroscopic images of a representative ovary of *gfp* control and *anmK*-KD females 6 days after detachment, scale bar = 500 µm. **F** The UV-Vis absorption scan of soluble (PBS) extract of ovaries of *gfp* and *anmK*-KD females 6 days after detachment. The inset shows an acetone extract of the ovaries, visualizing the haem deposits in the supernatant of ovaries homogenate. **G** Protein concentration of PBS-solubilised homogenates of tick ovaries. **H** SDS-PAGE (left) and Western blotting using antibodies against vitellogenin 1 and carrier protein 3 (CP3) on the soluble extract of ovaries from females 6 days after full engorgement. **I** Egg clutch weight. **J** Weights of formed larvae. **K** Representative images of formed larvae, scale bar indicates 200µm. **L** Success rate of blood-feeding of tick larvae resulting from fully-engorged females treated with dsRNA (n=6), originating from three independent blood-feeding experiments. **M** RT-qPCR evaluation of *M. mitochondrii* activity in the ovary, six days after detachment - *gyrB* transcriptional level was normalized against tick transcript for elongation factor alpha and used as a proxy of the bacterial cellular activity. Data significance was evaluated by t-test: *, p < 0.05; **, p < 0.01; ****, p < 0.0001; n.s. - not significant.

### 2.6 Cross-interaction between AnmK and endosymbionts of Metastriata ticks

Ticks often harbor one major symbiont in their ovaries (**Fig. 6F**). Genomic mining for genes coding for genes in PG recycling and *de novo* biosynthesis pathways revealed that *M. mitochondrii*, the symbiont of *I. ricinus* ticks, codes for its own AnmK enzyme, and retained essentially all genes of this pathway - although we noted a frameshift at the end of the sequence of *glmS*, which could result in a truncated but still functional protein sequence. The two genomes of *Francisella*-like endosymbionts (FLEs) also have an intact *anmK* gene, but this gene was apparently absent or showing signs of pseudogenization in other tick endosymbionts, including the genome of *Midichloria* associated with *Ixodes holocyclus* (which bore a frameshift in the middle of the sequence). Globally, most tick endosymbionts had several PG-related genes missing or pseudogenized, suggesting an impairment of their capacity to recycle or synthesize PG by themselves - with the exception of *Midichloria* from *I. ricinus* (**Fig. 6G**). To verify the role of AnmK on reproduction success and endosymbiosis interactions in other ticks, we silenced *anmK* in *H. longicornis* and *R. sanguineus*, both species being associated with a *Coxiella*-like symbiont that colonizes their ovary. Of note, *Coxiella*-like symbionts are known to be tightly interlinked with their tick hosts, affecting their blood-feeding biology^32^. As in *I. ricinus*, we observed a profound effect of *anmK* silencing in both *H. longicornis* and *R. sanguineus*, impairing their blood-feeding capacity (**Fig. 6A to E**), in particular for *H. longicornis*, which took more time to oviposit eggs after feeding (**Fig. 6D**). RT-qPCR analysis of symbiont activity showed that expression of 16S in *Coxiella*-like symbiont was significantly reduced upon *anmK*-silencing in its *H. longicornis* host (**Fig. 6H**), in contrast with what we observed in *I. ricinus*.

**Figure 6.**
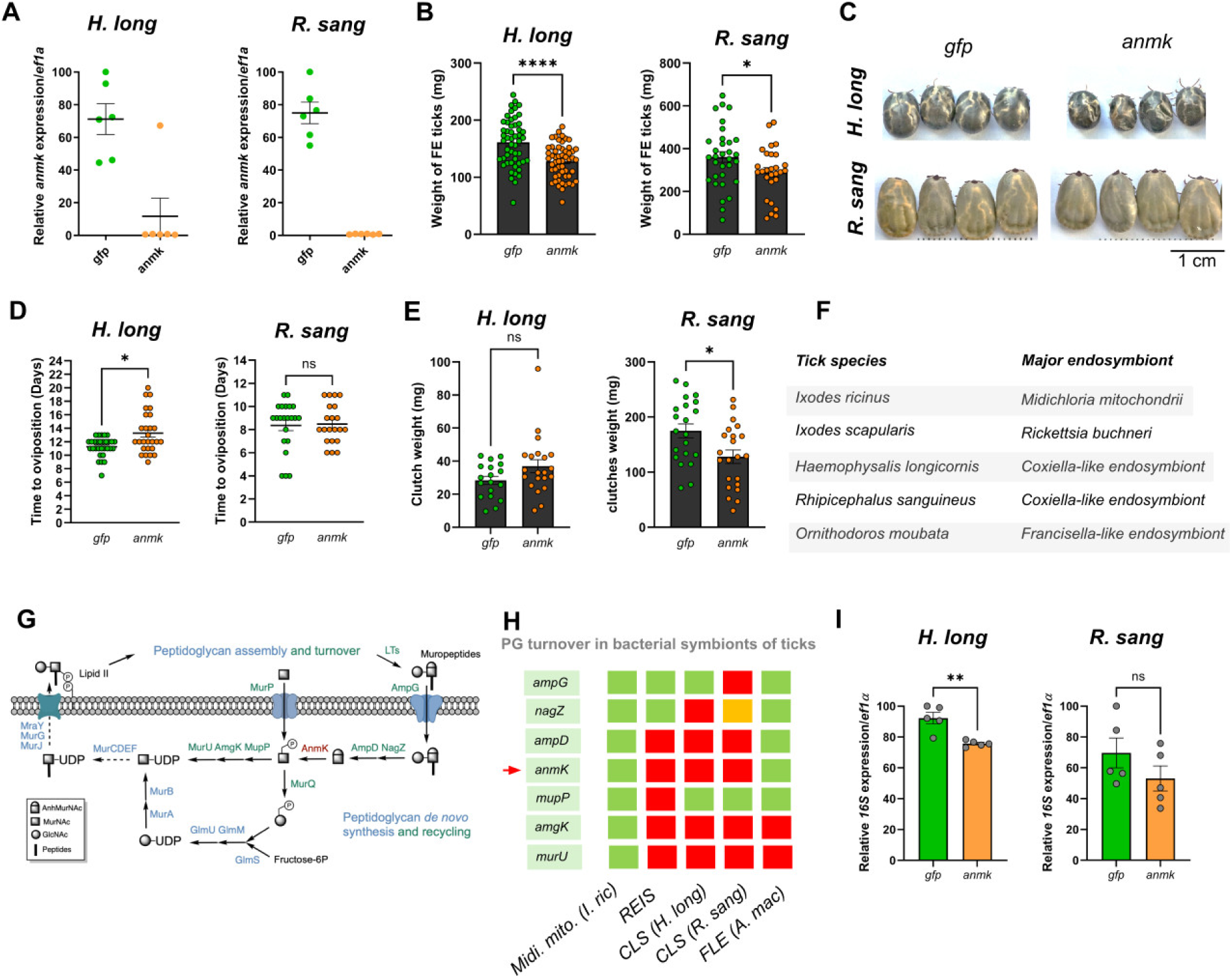
RNAi-silencing of *anmK* transcript in Metastriata representative species **A** RT-qPCR assessment of RNAi efficiency in ovaries of *H. longicornis* and *R. sanguineus* females - each data point represents an independent biological replicate. **B** The bar chart depicts the weights of spontaneously detached fully engorged females, n ≥ 30 for control (*gfp*) and *anmK*-silenced ticks (*anmK*) respectively. **C** Representative images of fully engorged females. **D** Duration of feeding till full engorgement and spontaneous detachment. **E** Clutch weight**. F** Major bacterial endosymbionts for five representative tick species. **G** Genes of the standard PG recycling and *de novo* biosynthesis pathways in bacterial genomes (left). **H** Genes found to be absent (red), pseudogenized (orange) or present and intact (green) in tick endosymbionts. **H** Effect of *anmK* knockdown on the expression of symbiotic genes.

## Discussion

Current knowledge of HGT in ticks is limited, with a few cases of gene transfers from vertebrates described, while only one functional bacterial gene transfer has been documented in the common ancestor of ticks and other Chelicerata ^22^. To broaden this understanding, leveraging the recent availability of several assembled genomes for ticks from the *Ixodes* genus ^31^, we conducted a systematic, genome-wide HGT screen across the genome of *I. ricinus*. Our method compares predicted protein sequences against general databases, and analyses these results in a phylogenetic framework, allowing detection of ancestral HGT events that may otherwise go unnoticed with for example simple BLAST-based searches. Using this approach, we identified a diverse set of HGT candidates originating from vertebrates, viruses, and bacteria.

Methods used to detect HGT, whether in previous studies or in the present analysis, can yield different results and have inherent limitations. First, false positives may arise^33^. Our analysis identified indeed several probable false positives, particularly among sequences initially annotated as vertebrate-derived. Phylogenetic reanalysis revealed that these sequences have undergone significant divergence in ticks, resulting in misleading similarities to vertebrates and complicating HGT detection. These cases are reminiscent of two reported events of vertebrate-to-tick HGT^14,21^, where very low conservation and the involvement of short sequence fragments can be explained by convergence rather than an event of horizontal transfer. Second, our approach is restricted to annotated protein-coding genes. Recent studies using whole-genome BLAST searches in *I. ricinus* ^24,34^ identified multiple bacterial DNA fragments, mainly from *Spiroplasma* and *Midichloria*, integrated into the tick genome. However, the largest laterally transferred block (called “LGT”) contained multiple stop codons, suggesting it contains non-functional remnants rather than domesticated genes, thus explaining why it was not detected in our protein-focused analysis. Third, complex phylogenetic histories can obscure genuine HGT events. This was the case of *dae2*, which our method failed to identify as HGT due to complex patterns of duplication, loss, and possible contamination. As found by Chou and collaborators^22^, the presence of similar genes in other Acari supports the hypothesis that the gene transfer occurred in a common ancestor of Acari. However, sequences from phylogenetically more distant organisms, including a springtail (*Allama fusca*, Collembola), a fungus gnat (*Bradysia coprophila*, Insecta), and a scorpion (*Centruroides sculpturatus*, Chelicerata), unexpectedly also grouped within the same clade. Such a patchy taxonomic distribution does not suggest an event of HGT in a common ancestor of insects, Collembola, and ticks. Our findings thus highlight a more intricate history for *dae2* than initially described^22^, with the presence of several duplications of *dae2* following its horizontal transfer, and possible secondary transfers to other arthropods.

Our study revealed, however, well supported cases of transfer from viruses and bacteria. For example, we uncovered several annotated tick genes corresponding to endogenous viral elements (EVEs), many consistent with elements previously uncovered by virome studies^35^. Because our pipeline relied on annotated protein sequences, the actual number of EVEs we found is likely largely underestimated. It has been shown that EVEs can provide immune protection against related exogenous viruses via RNA interference (RNAi) pathways^36,37^. We observed a cluster of EVEs from different viral groups in the genome of *I. ricinus*, suggestive of roles in piRNA production, a phenomenon consistent with observations in mosquitoes^38^ and shrimps^39^. Most EVEs identified here appear degraded; however, the virus-derived *I. ricinus* gene IricT00012932 retains an intact open reading frame (ORF) and shows substantial expression, suggesting a possible function. Future studies examining complete tick genome assemblies, beyond coding annotations, will be essential to fully characterize EVEs and their evolutionary roles^40^.

For bacterial candidates, to avoid spurious cases, especially potential contaminants, we applied a stringent filter requiring presence in multiple tick genome assemblies, reducing the candidate list significantly, ultimately down to two robust cases. We thereby identified two previously unrecognized HGT events in ticks: a GNAT-family acetyltransferase transferred to the ancestor of ticks and Mesostigmata, and an *anmK* gene transferred to the last common ancestor of all ticks. Both genes satisfy multiple conservative criteria for HGT: high AI scores, presence in all tick genomes examined, integration into host gene structure (including introns and splicing variation), and robust phylogenetic placement within bacteria rather than animals. Codon usage patterns further support their long-term domestication. Genes of the GNAT-family are known for their propensity for horizontal transfer^41^, with documented events into fungi and soil nematodes. Our study demonstrates here a *Burkholderia*-like donor for its origin in an ancestor of ticks and Mesostigmata. By contrast, *anmK* has not previously been reported as horizontally transferred into eukaryotes, to our knowledge. Our phylogenetic analysis places the donor within Chitinophagaceae. Members of this bacterial group have been reported from the midgut of *I. ricinus* ^42^ and on vertebrate skin ^43,44^, the major point of contact during tick feeding. These observations suggest two plausible scenarios, including either a direct transfer from transient gut microbiota, or a transfer facilitated by the tick–host interface, possibly via bacteria associated with host skin. Because the transferred *anmK* sequence must have ultimately reached the germline, the latter possibility highlights how tick–host interactions may have provided opportunities for bacteria to invade reproductive tissues, although several barriers had to be overcome for HGT to occur. AnmK in bacteria participates in the PG recycling pathway, converting 1,6-anhydro-MurNAc into MurNAc-6-P, thereby contributing substantially to cell-wall turnover (representing a salvage pathway roughly as important as *de novo* PG biosynthesis) ^45^. Such transfer has thus intriguing parallels to horizontally transferred PG pathway genes observed in other symbiotic systems (e.g., mealybugs). The high expression of *anmK* in the ovaries during blood feeding, first shown by RNA-Seq data for *I. ricinus*, points to a functional role potentially tied to reproduction and symbiosis. We thus focused primarily on this gene and its potential role in tick biology.

RNA-Seq evidence and sequencing of both the cDNA and the genomic region indicated that *anmK* has acquired a eukaryotic gene structure, including an intron between the 5′ UTR and coding region, demonstrating its genomic and transcriptional assimilation. Phylogenetic analyses across multiple *Ixodes* species and Mesostigmata outgroups show *anmK* is present in all ticks but absent from non-tick Chelicerata, placing the transfer early in tick evolution (150–300 Mya). Conserved micro-synteny between tick species further supports a single, ancient introgression event followed by vertical inheritance. Lastly, an independent transfer of *anmK* to rotifers is also evident in our phylogenetic analysis, consistent with a high rate of HGT in this group. The biological function of this gene in rotifers remains to be determined.

Our comparative study of expression, using different tissues and time-points in the tick life-cycle, showed that *anmK* expression peaks in the ovaries of adult female ticks during and shortly after blood feeding, precisely when maternally inherited endosymbionts undergo rapid proliferation. This was observed both in *Ixodes* and non-*Ixodes* species of ticks. The conservation of this ovary-specific expression pattern across divergent tick lineages argues that *anmK* contributes to a fundamental and possibly symbiosis-dependent aspect of tick reproduction, possibly for the management of an adequate symbiont population or to control the vertical transfer of symbionts. Ovarian tissues represent a unique interface where host physiology, nutrient transfer, and bacterial activity converge, and the coincident upregulation of *anmK* during this window of symbiont expansion points to a functional link. One possibility is that *anmK* participates in modulating the PG pathways of endosymbionts, either by salvaging cell wall intermediates released during bacterial replication or by influencing the turnover of microbe-derived metabolites in the ovary. Alternatively, *anmK* may serve a host-centric role, buffering the ovary against microbial by-products at a stage when vitellogenin uptake and oocyte maturation are maximal. Our results point to different impacts of *anmK* silencing in different bacteria, since the bacterial activity (using 16S rRNA as a proxy) decreased in non-*Ixodes* species, but not in *I. ricinus*. This suggests there could be important differences across ticks regarding how this bacterial-derived enzyme shapes host–microbe interactions, which is not surprising considering that different tick species are associated with different species of symbiotic bacteria, differing for example by their degree of retention of genes of the PG pathway. Elucidating the role of tick *anmK* and its impact on bacterial symbionts will thus require investigating several tick-symbiont systems.

Biochemical assays show that tick AnmK retains detectable activity on 1,6-anhydro-MurNAc, but with substantially reduced catalytic efficiency compared to its bacterial homologs under the tested conditions. Of note, we found the activity of IrAnmK to be higher at acidic conditions, which contrasts with the properties of AnmK in *E. coli* and which could indicate a changed catalytical micro-environment. However, our observation of a strong reduction in IrAnmK activity rather prefigures that substrate specificity has changed. *In vivo*, IrAnmK does not colocalize with endosymbionts; instead, it accumulates in the oocyte cytosol, around the chorion and yolk granules. This anatomical distribution, combined with the reduced activity on classical MurNAc, suggests that tick AnmK may have evolved a modified or expanded substrate range, potentially acting on chitin-derived or other amino-sugar intermediates abundant in reproductive tissues. We determined that all the residues critical for catalysis are conserved within IrAnmK and the other known tick AnmK protein sequences, indicating the potential retention of the catalytic and binding properties in the domesticated tick enzymes. However in the AnmK clade 2 of Chitinophagaceae, several variations were observed. Further work is intended to elucidate the functional differences of the two AnmK versions of Chitinophagaceae, as well as the tick variants which emerged from clade 2. We discovered, however, a potentially important substitution in tick AnmKs, i.e. the N168C substitution (**Fig. S8**). Yet, an exchange of the cysteine back to asparagine did not dramatically affect activity, indicating that this substitution may not be critical for activity, but possibly for the binding of an alternative substrate. Also, preliminary 3D modelling indicates that the catalytic pocket of the tick enzyme appears “less open” than in bacteria, consistent with altered substrate accommodation. Further structural work is needed to confirm whether conformational changes underlie its reduced activity and potential neofunctionalization.

RNAi-mediated knockdown of *IranmK* impaired vitellogenin uptake, oocyte maturation, and offspring feeding success in the tick *I. ricinus*, demonstrating that the gene is now integral to tick reproductive physiology. In two species of Metastriata ticks hosting *Coxiella-like* endosymbionts (bacteria devoid of a coding *anmK*), we found that blood-feeding capacity (an inherent feature of all ticks supporting their large fecundity) was profoundly compromised after *anmK* silencing. These phenotypes highlight how bacterial genes can be domesticated to support essential host functions. PG is essential for most bacteria, but intracellular symbionts often reduce or modify their PG layer because they inhabit osmotically protected environments ^46^. In this context, host-encoded bacterial enzymes can compensate for missing bacterial encoded enzymes, thereby modulating PG turnover and limiting symbiont proliferation. While tick *anmK* could contribute to this process, as seen in mealybug symbioses, the absence of a strict association between tick *anmK* presence and endosymbiont *anmK* absence, alongside its reduced catalytic efficiency, suggests that its function in ticks extends beyond canonical cell-wall biochemistry. It is worth highlighting that *anmK* shows no co-expression with the horizontally transferred antibacterial amidase *dae2*, and the two genes display distinct tissue and temporal profiles, supporting independent evolutionary trajectories and functions.

All these observations prompt us to formulate alternative hypotheses about tick AnmK: (i) tick AnmK could exhibit molecular activity toward alternative substrates during chorion formation and vitellogenesis; (ii) tick AnmK could be involved in modulating the tick immune system, potentially interacting with peptidoglycan recognition protein pathways that respond to bacterial PG fragments; and (iii) this enzyme could play a more general role in managing microbe-derived sugar metabolites, preventing their accumulation during periods of intense symbiont replication. These hypotheses align with broader patterns observed across eukaryotes, where horizontally acquired bacterial genes, particularly those involved in cell-wall metabolism, are frequently co-opted into host immune or metabolic pathways (e.g., in rotifers ^47^).

## Conclusions

Through a combination of phylogenetics, gene structure, and conserved microsynteny, our work identified two previously unrecognized bacterial genes, a GNAT-family acetyltransferase and the PG recycling kinase *anmK*, as ancient horizontal acquisitions that have been fully domesticated into tick genomes. Functionally, we demonstrate that *anmK* is not a silent relic of bacterial metabolism but a conserved determinant of reproductive fitness. RNAi knockdown of *anmK* indeed causes defects in blood feeding and reproduction in different tick species, both in *I. ricinus* and two non-*Ixodes* hard tick species, with evidence of a cross interaction between *anmK* and bacterial activity in the non-*Ixodes* species. Overall, our results indicate that this domesticated enzyme underpins core reproductive functions across hard tick species, probably representing an ancestral evolutionary innovation that shaped the biology and global evolution of this group of arthropods.

Future work should determine whether tick *anmK* activity extends beyond canonical PG recycling and resolve how its activity is sensed and regulated within the ovary. Dissecting the biochemical and structural diversification of *anmK* after its transfer, and clarifying its interplay with other horizontally acquired genes, will shed light on how bacterial metabolic enzymes can be repurposed to coordinate symbiosis and reproduction in hematophagous vectors, and may expose lineage-spanning vulnerabilities that could be targeted for tick control.

## Methods details

### Tick maintenance and feeding

*I. ricinus* females were collected by flagging forested areas around Ceske Budejovice, Czech Republic, whereas *I. scapularis*, *R. sanguineus* and *H. longicornis* were obtained from the tick rearing facility of the Institute of Parasitology, Biology Centre, Czech Academy of Sciences. Ticks were fed on laboratory guinea pigs or laboratory mice (Animal facility of Institute of Parasitology). All experimental animals were handled in accordance with the laws of the Czech Republic (Act No. 246/1992 Coll., on the Protection of Animals against Cruelty; ethics approval No. 25/2020).

### Horizontal gene transfer detection

The detection of horizontal gene transfer was performed using the AvP software (Alienness vs Predictor, v1.0.2 ^48^) on the predicted protein sequences from *I. ricinus*, based on our recent study of new genome assemblies from this species^31^. Detection of HGT with AvP is based on a comparison of hits in BlatstP results, taking into account taxonomy and the expected order of similarity scores. In order to detect potential cases of horizontal gene transfer from distantly related groups of organisms (e.g. vertebrates, bacteria and viruses) to ticks, with a focus on HGT events occurring in a common ancestor of ticks or of ticks and other Acari, we defined Arthropoda as the “ingroup” and Acari as the “recipient group” (**<u>Fig. S1</u>**). With these parameters, all hits to Acari were masked, and the relative scores of hits to organisms expected to be phylogenetically closer (other Arthropoda) or more distant (e.g. vertebrates, bacteria, viruses) were compared. Hits with a higher score in the latter category, which is unexpected, were therefore HGT candidates. Although the monophyletic support for Acari is debated in the literature, our approach is simply conservative and allows to detect HGT from distant organisms that occurred both in the ancestry of any of the four *Ixodes* genomes screened, including common ancestors to all ticks and common ancestor to ticks and any other group belonging to Acari. First, tick protein sequences were aligned against the nr NCBI database (downloaded the 5th October 2023) using Diamond (v2.0.8) ^49^ with the following parameters: -k 500 (maximum number of target sequences per query) and --evalue 5e-2 (e-value threshold). We then ran AvP, choosing the alien index (AI) value to identify donor groups from the parsed BLASTP results. Two runs of AvP were performed with two different AI cut-off values (20 and 40) and default parameters (--percent_identity 100 –cutoffextend 20 –trimal false –min_num_hits 4 –percentage_similar_hits 0.7 –fastml: true –node_support 0 – complex_per_ingroup 20 –complex_per_donor: 80 –complex_per_node 90; mafft_options: ‘--anysymbol --auto’; trimal_options: ‘-automated1’; iqmodel: ‘-mset WAG,LG,JTT -AICc -mrate E,I,G,R’; ufbootstrap: 1000; iq_threads: 1). To limit the number of false positives (which can result from contaminant sequences or taxonomical mis-assignment), we kept only candidates with at least five sequences both in the recipient and in the donor groups based on the parsing of hit tables.

### Sequence alignment and phylogenies

Complementary searches of hits in organisms of interest were made through alignments against the NCBI databases (nt and transcriptome shotgun assemblies -TSA) using TBLASTN or against the nr database with BLASTP ^50^. For example, for bacterial HGT candidates, a blastp was performed against Bacteria only, retrieving the 500 first hits. To avoid analysing multiple very closely related sequences, we used CD-HIT with the parameter -c 0.7, and all the sequences retained were included in the subsequent analyses.

Candidate sequences and hit sequences were aligned using MAFFT (v7.505 ^51^) with the following parameters for bacterial HGTs: --leavegappyregion --maxiterate 1000 --retree 2 --genafpair; and with the following parameters for the viral HGTs: --maxiterate 1000 --genafpair. Alignments were then trimmed with TrimAl (v1.4.1 ^52^) using a gap threshold of 0.6. Maximum likelihood phylogenetic trees were built with IQ-TREE (v2.2.2.6 ^53^) with the following parameters: --alrt 1000 -B 1000 --seqtype AA -m TEST --merit BIC. Trees were finally visualized using the R packages treeio (v1.22.0 ^54^), ggtree (v3.6.2 ^55^) and ggplot2 (v3.4.2 ^56^). Alignment figures were generated using the R package ggmsa (v3.18 ^57^).

A complementary search of one candidate gene with a HGT origin, *anmK*, was done in the genomes of four groups of bacteria identified as tick endosymbionts. For this, we selected the most complete and continuous tick symbiont genomes available, our resulting list including two *Francisella*-like endosymbionts (GCA_001753795.1 and GCF_003069505.1), four *Coxiella*-like-endosymbionts (PRJNA301476, NZ_CP094378.1, NZ_CP094229.1, and CP011126.1) two *Midichloria* (CP002130.1 and CP094379.1) and one *Rickettsia*-like tick endosymbiont (GCF_000160735.1). A specific phylogenetic analysis including these sequences and those of the putative donor group was performed. Finally, we also searched in the same genomes other genes of the PG pathway, known to be involved in the de novo biosynthesis or recycling of PG (see Results).

Throughout the manuscript, gene names are written in italics (e.g. *anmK*, *IranmK*, *GNAT*), and protein names in regular font, including when a prefix indicates a species name (e.g. IrAnmK to designate the protein AnmK specifically in *I. ricinus*).

### Codon usage

Coding sequences of the genome of *I. ricinus* (assembly GCA_964199305.2) were analyzed through factorial correspondence analysis of their relative synonymous codon usage (RSCU) with CodonW (https://codonw.sourceforge.net/), in order to compare candidate genes identified as horizontally transferred by phylogenetic methods and the rest of the genome. Spurious candidates (e.g. contaminant sequences) or recent transfers are expected to be outliers, while ancient transfers are expected to have the same codon usage as the rest of the genome. We also distinguished ribosomal proteins, expected to display an optimal codon usage; indeed, because of their consistently high level of expression, ribosomal proteins are under stronger selection for translation efficiency than standard genes.

### Gene expression profile based on RNA-Seq data

Gene expression profiles of *I. ricinus* genes were retrieved from the expression atlas generated in ^31^ for which normalized counts of expression, measured in transcripts per million (TPMs) were produced for different conditions, tissues, and stages of the tick life-cycle.

### Amplification of *IranmK* and mRNA quantification

Genomic DNA (gDNA) of *I. ricinus* was extracted from a pool of adult females using the NucleoSpin Tissue Kit (Macherey-Nagel) according to the manufacturer’s instructions. Total mRNA was isolated from ovaries of fed adult females of *I. ricinus*, *H. longicornis*, *R. sanguineus* and *O. moubata* 4 days after detachment from a guinea pig, using the NucleoSpin RNA kit (Macherey-Nagel). The mRNA was transcribed into cDNA using random hexamers and the Transcriptor High-Fidelity cDNA Synthesis Kit (Roche) (0.5 μg RNA per 20 μl reaction) and then diluted 20-fold in sterile water. PCR was performed using Platinum SuperFi II DNA Polymerase (Invitrogen). The amplified fragments were cloned using the Zero Blunt TOPO PCR Cloning Kit. The gDNA-derived insert was sequenced using whole-plasmid sequencing (Eurofins genomics) and the cDNA-derived insert was sequenced using Sanger sequencing guided by M13 primers (M13F+M13R). For expression profiling, RNA was obtained from different developmental stages of the tick (larvae, nymphs, adults) as well as from the tissue of non-fed, semi-fed, and fully fed (4 days after feeding) adult females ^58^. Total RNA was isolated using the NucleoSpin RNA Kit (Macherey-Nagel); mRNAs were transcribed into cDNA using random hexamers and the Transcriptor High-Fidelity cDNA Synthesis Kit (Roche, 0.5 μg RNA per 20 μL reaction) and then diluted 20-fold in sterile water. All samples were prepared in triplicate for qPCR. Primers were designed in Primer3 (http://bioinfo.ut.ee/primer3-0.4.0/) and verified by PCR using the Fast Start Master mix (Roche). Expression levels of *anmK* were determined by qRT-PCR using the QuantStudio 6 Flex Real-Time PCR System (Applied Biosystems) and LightCycler 480 SYBR green I Master chemistry (Roche). Expression was normalized using tick elongation factor 1 (*ef1a*; GenBank ID: GU074769). For Sanger sequencing, the amplified gDNA/cDNA amplicons were cloned in the Zero Blunt TOPO PCR vector. For cDNA sets of multiple tick species, caeca of midgut were washed out of lumen contents, each ovary was split in half for RNA and DNA extraction and the remaining white tissue (salivary glands, fat body/trachea, Malpighian tubules and synganglia) were collected (denoted as rest) from females four days after full engorgement. Primers against species-specific *anmK* and *ef1a* are available in Supplementary Text. Responsiveness of *anmK* transcripts to microinjected model microbes was conducted as previously described ^59^.

### RNAi of anmK in I. ricinus, R. sanguineus, and H. longicornis

DNA fragments of the *anmK* gene, 418 bp from *I. ricinus* (IricT00013998), 418 bp from *R. sanguineus* (XM_037664325) and 418 bp from *H. longicornis* (JABSTR010000011), were synthesized as GeneArt Strings DNA (Invitrogen), see Supplementary Text for sequences of individual dsRNA. Each fragment was flanked by the sequences gcgaattgggtACc<u>gggccc</u> and gggccgcggtggcggccgctc<u>tctaga</u> containing ApaI and XbaI restriction sites (underlined) and overhangs compatible with Gibson Assembly. Fragments were cloned into an in-house-modified pll10 vector containing two T7 promoters in reverse orientation ^60^ using the NEBuilder HiFi DNA Assembly Kit (New England Biolabs). Double-stranded RNA (dsRNA) was synthesized using the MEGASCRIPT T7 Transcription Kit (Ambion) as previously described ^61^. As a control, GFP dsRNA was synthesized from the pLL6 plasmid under the same conditions. The dsRNA (3 μg/μL, adult tick = 280 nL) was injected into the hemocoel (through the coxa III) of the unfed tick using the microinjector (Drummond). After inoculation, the ticks rested in a humid chamber for three days and were fed on guinea pigs. After feeding, fully engorged females were re-injected the same amount of dsRNA. Success rate of blood feeding is defined as a number of ticks fully engorged and spontaneously detached from the host, out of the initial number of unfed ticks placed on the animal. Silencing of *anmK* by RNAi was verified by RT-qPCR as described above.

### Cloning and Expression of Recombinant AnmK

The complete sequence of the *anmK* gene of *I. ricinus* (GIXL01000730) was amplified by PCR from genomic DNA using the Platinum™ SuperFi II DNA Polymerase (Invitrogen) and the following primers (5’-->3’): Forward - CAGAGAACAGATTGGTGGTATGACCGGAGGCGCTCAAG, Reverse - TGCGCCGAATAAATACCTAAGCTTGTCTTCAGACCTTCCCGCACCAG. The complete sequence of the *anmK* gene from *E. coli* strain K-12 substrain MG1655 (NC_000913.3:c1719602-1718493) and *T. pollutisoli* (WP_276500879) were synthesized as GeneArt String DNA (Invitrogen) with CAGAGAACAGATTGGTGGT and AGACAAGCTTAGGTATTTATTCGGCGCA overhangs for Gibson cloning. The amplified DNA fragments were cloned into an in-house-modified pET SUMO plasmid (Invitrogen) with BsrGI-HF and HindIII restriction sites using the NEBuilder HiFi DNA Assembly Kit (New England Biolabs). Sequencing of the plasmids confirmed that the amino acid sequences of the cloned fragments were 100% identical to the respective templates.

The plasmid constructs were transformed into BL21(DE3) *E. coli* cells and grown overnight at 37°C in 10 ml LB medium with kanamycin and 1% glucose. The next day, 200 ml of LB medium with kanamycin was inoculated with the overnight culture and grown at 37°C with shaking at 200 rpm until the OD600 reached 0.5. The cells were then induced by addition of IPTG at a final concentration of 1 mM and grown for a further 4.5 hours. The cells were harvested by centrifugation and the resulting pellet was stored at −20°C until further use.

### Purification of Recombinant AnmK

The pellet of 200 ml of the bacterial culture was resuspended in a 20 ml buffer containing 50 mM TRIS-Cl (pH 9), 50 mM NaCl, 10 mM imidazole and 0.5% Triton X-100. The solution was then mixed with 5 mg lysozyme (Merck, 30,000 U/mg protein) and sonicated twice for 1 minute each time. The lysate was centrifuged several times at 17,000 g at 4°C and filtered through a 0.22 µm filter (TPP). The sample was then loaded onto a 5 mL HiTrap Ni2+-IMAC FF column (GE Healthcare) for purification via the His-tag located before the SUMO tag using ÄKTA purifier (GE Healthcare). Impurities were removed by washing the column with 20 mM and 40 mM imidazole. The target protein was eluted with a gradient of imidazole to a final concentration of 0.5 M. The eluted protein fraction was dialyzed overnight in 4 liters of 50 mM TRIS-Cl (pH 9.0) and 150 mM NaCl at 10°C. The next day, the sample was centrifuged at 17,000 g at 4°C and filtered through a 0.22 µm filter. SUMO tag cleavage was achieved by adding 1 µL of SUMO protease (Sigma, SAE0067, 25 U/µl) to 15 mL of the sample and incubating overnight at 10°C with gentle agitation. The sample was then centrifuged again at 17,000 g at 4°C and filtered through a 0.22 µm filter. To remove uncleaved protein, impurities, the cleaved SUMO tag and the SUMO protease, the sample was passed through another 5 mL HiTrap IMAC FF column charged with Ni^2+^ ions. The flow-through containing the cleaved protein was dialyzed overnight in 4 liters of 50 mM TRIS-Cl (pH 9.0) and 150 mM NaCl at 10°C. The protein solution was then concentrated to 2 mL using an Amicon Ultra-15 Centrifugal Filter 10 kDa MWCO (Merck, Sigma-Aldrich, UFC9010). The final protein product was mixed with glycerol (10% final), frozen in liquid nitrogen, and stored at −20°C.

### Generation, isolation, and labelling of anti-IrAnmK IgY antibodies

Recombinant IrAnmK (50 µg) was diluted with addition of 50 mM TRIS-Cl (pH 9) and 150 mM NaCl to a final volume of 500 µl. This protein preparation was mixed 1:1 with Complete Freund’s adjuvant and injected subcutaneously, and formulated with incomplete Freund’s adjuvant and injected in three subsequent doses (weeks 3, 5, and 7) as described previously ^62^. Two weeks after the last dose, 1 mL of hen’s blood was taken and serum obtained. To isolate IgY antibodies from hen’s immune sera, 2 mL of immune sera were combined with 4 mL of sodium acetate buffer (50 mM, pH 4.0), with gradual addition of caprylic acid (Serva, 15771) while stirring. The precipitated sample was centrifuged at 2500 × *g* for 10 minutes. The resulting supernatant was filtered using a 0.22 µm filter (Techno Plastic Products, 99722, Lot 20220354) and dialysed against 5 mM sodium phosphate (pH=7.2). The protein concentration of the final IgY preparation was determined using the Bradford to be 475 µg/mL. The IgY preparation was either used for western blotting or to conjugation with Alexa Fluor™ 647 NHS Ester (Succinimidyl Ester), according to the manufactureŕs protocol (Thermo Fisher Scientific, A37573).

### Western Blot and Immunolocalization

Fully engorged females were rested after detachment from the host, and their ovary was dissected. Proteins were extracted in 8M Urea, 500 mM NaCl, 50mM Tris-Cl pH 8 using a 29G syringe. Samples were then centrifuged 15,000 × g, for 10 min at 4°C. Electrophoretic samples for SDS-PAGE were prepared in reducing Laemmli buffer supplemented with β-mercaptoethanol. Identical volumes of the protein extract were applied per lane. Proteins were separated on gradient (4–15%) Criterion TGX Stain-Free Precast gels (BioRad, Hercules, CA) in Tris-Glycine-SDS running buffer (25 mM Tris, 192 mM glycine, 0.1% (w/vol) SDS, pH 8.3), visualized using TGX stain-free chemistry (BioRad), and transferred onto the PVDF membrane. For Western blot analyses, membranes were blocked in 3% (w/vol) non-fat skimmed milk in PBS with 0.05% Tween 20 (PBS-T), incubated in immune serum diluted in PBS-T (αIrAnmK-1:1000), and then in the rabbit anti-chicken IgG-peroxidase antibody (Sigma No A9046) diluted in PBS-T (1:50,000). Signals were detected using ClarityWestern ECL substrate (Number #1705061), visualized using a ChemiDoc MP imager, and analysed using Image Lab Software (BioRad).

Unfed females of *I. ricinus* were injected with dsRNA *IranmK* (280nl; 3µg/µl) or GFP (control), rested for 1 day and fed on guinea pigs. After detachment, the females were re-injected with dsRNA (280nl; 3µg/µl). Four days after detachment, ovaries were dissected in a phosphate buffer (PBS) and fixed in 4% paraformaldehyde (overnight, 4 °C). Then the ovaries were washed 3 × 10 minutes with PBS and transferred to ethanol series (70, 96, 100%; 2 × 30 minutes each) and chloroform. The samples were embedded in Paraplast overnight at 62 °C and sectioned to a thickness of 7 μm. The sections were deparaffinized, rehydrated and stained. For immunofluorescence detection after deparaffinization, sections were washed 2× for 5 minutes in PBS and then blocked in 1% bovine serum albumin (Sigma, No A9647) and 10% goat serum in 0.1% PBT-Tx (Triton X-100) for 1 hour at RT. Then these sections were incubated with primary chicken αIrAnmK antibody coupled with Alexa 647 dye (1:100 in PBS-Tx (0.1% Tx in PBS) overnight at 4°C. Slides were washed 2×10 min in PBS-Tx and nuclei were counterstained with DAPI (4,6-diamidino-2-phenylindole (Thermo Fisher Scientific, 62248) for 10 min and washed 2×10 min in PBS. The slides were then embedded in DABCO and examined with an Olympus FluoView FV3000 confocal microscope (Olympus, Tokyo, Japan).

### Investigation of the AnmK Reaction by HPLC-MS Analysis

An *in vitro* reaction was conducted by incubating AnmK from *E. coli* or *I. ricinus* (EcAnmK, IrAnmK; final concentration of protein 5 µM each) with ATP, MgCl_2_ and anhydroMurNAc (final concentrations 10 mM, 10 mM and 1 mM, respectively) in a Tris buffer, consisting of 100 mM Tris-HCl pH 9.0. A final volume of 50 µL was incubated at room-temperature for 20 h. Afterwards the reaction was stopped by incubation at 95°C for 5 min. To remove coagulated protein, the mixture was centrifuged (13400 rpm, 10 min, RT) and 40 µL of the supernatant were used for evaluation by LC-MS.

For LC-MS measurements a high-performance liquid chromatography (HPLC) system (UltiMate 3000, Dionex) is connected to an electrospray ionization-time of flight (ESI-TOF) mass spectrometer (micrOTOF focus II, Bruker), which is operated in negative-ion mode. 5 µL were injected into a Gemini C_18_ column (150 x 4.6 mm, 110 Å, 5µm; Phenomenex). For separation, a 45-min gradient was used with a flow rate of 0.2 ml/min, while the column compartment was kept at 37°C. The program begins with 5 min of washing with 100% buffer A (0.1% formic acid with 0.05% ammonium formate), followed by a linear gradient from 0% to 40% buffer B (100% Acetonitrile) over 30 min. The method is completed by a 5 min hold at 40% buffer B (60% buffer A) and 5 min of re-equilibration at 100% buffer A.

### Monitoring of the AnmK reaction by an ADP/NADH coupled assay

The AnmK reaction was continuously monitored by a coupled enzyme assay in 96-well plates. Reactions were conducted in a volume of 100 µL with 10 U µL^-1^ pyruvate kinase (PK, Merck), 15 U µL^-1^ L-lactate dehydrogenase (LDH, Merck), 1 mM anhydroMurNAc, 2 mM MgCl_2_, 2 mM ATP, 1 mM PEP, 0.5 mM NADH and 1 mM DTT, using either a Bicine buffer at pH 9.0 (50 mM Bicine, 50 mM NaCl, pH 9.0) or a Bis-Tris buffer at pH 6.0 (50 mM Bis-Tris, 50 mM NaCl, pH 6.0). To start the reaction an appropriate amount of enzyme (EcAnmK and IrAnmK at final concentrations of 0.5 µM and 5 µM, respectively) was added. The reaction was incubated in a spectrophotometer (BioTek Epoch2, Thermo Fisher Scientific) at 37°C for 180 min and the absorption at 340 nm was measured in 30 second intervals. Reaction progression can be continuously monitored with this assay due to the regeneration of ATP from ADP by PK. Thereby pyruvate is formed and subsequently reduced to lactate by LDH under the consumption of NADH. Consumption of NADH is followed by monitoring decline in absorption at 340 nm.

To calculate the reaction rate a background control (at 340 nm) with water instead of AnmK was measured. At each timepoint (x min) the decline in absorption in the water probe was calculated by subtracting the absorption at t = 0 min from the absorption at t = x min. The same was done for the respective AnmK probe at each timepoint. To calculate change in OD (ΔA340nm) the decline in absorption in the blank probe (water) at time point x is subtracted from the decline in absorption in the respective AnmK probe. The calculated values for ΔA340nm are transferred to GraphPad Prism 8.4.3., where the slope of the reaction progress is calculated by using the “straight line” option for nonlinear regression. For determination of the reaction rate, the slope was calculated for the linear portion of the curve.

## Declarations

### Data availability

All data and code reported in this paper will be shared by the corresponding authors upon reasonable request.

### Ethical statement

The laboratory animals were treated in accordance with 3Ŕs and the Czech law for the Protection of Animals against Cruelty No. 246/1992 Coll. together with decree No. 419/2012 Coll.. The experimental work on animals was approved by Ministry of Agriculture permit No. 50-2022-P.

### Authors contributions

Conceptualization, A.C.A.,C.M., J.P. and C.R.;

Methodology, A.C.A., C.M., J.P. and C.R.;

Investigation, A.C.A., O.H., V.U., T.Š., L.R., A.T., O.P., Y.S.,C.M., J.P. and C.R.;

Resources, A.T., C.M., J.P. and C.R.;

Writing - Original draft, A.C.A., Y.S., C.M., J.P. and C.R.;

Writing - Review & Editing, A.C.A., O.P., Y.S.,C.M., J.P. and C.R.;

Supervision: C.M., J.P. and C.R.

Funding acquisition, C.M., J.P. and C.R.

## Acknowledgements

Thanks to the Genotoul platform for access to its bioinformatics and calculation resources, to Pierrick Lucas for help with data visualization, and Libera Lo Presti for writing support. CM acknowledges infrastructural support from the Cluster of Excellence EXC 2124 ‘‘Controlling Microbes to Fight Infections’’ (390838134) of the German Research Foundation.

## Funding

The work was funded by the Czech Science Foundation grant 22-18424M (to JP), and also by a private partnership (grant TickTarget, funding ACA’s postdoc), the Czech Science Foundation 25-16064S (to OH), Czech Ministry of Education, Youth and Sports grant ERC CZ LL2503 (to JP), and the project “Biology of Hyaluronic Acid” (No. CZ.02.01.01/00/23_020/0008499) funded by the Ministry of Education, Youth, and Sport, Czech Republic and the European Regional Development Fund (to JP, OH).

## Notes

### Competing Interest Statement

The authors have declared no competing interest.

