## Supplemental Information for "The Bacterial Kinase AnmK Integrates into Tick Genome and Biology"

#### Supplementary Figures

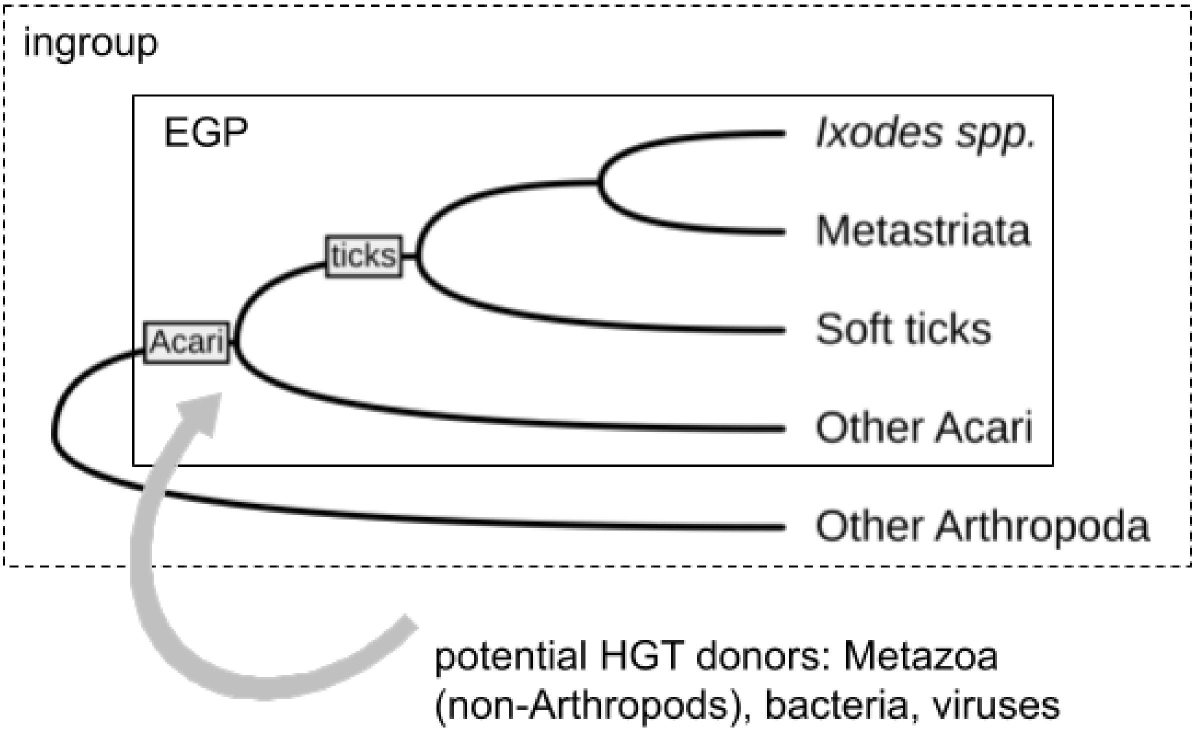


**Figure S1. Method to detect HGT events in ticks and other Acari.**

Phylogenetic relationships are illustrated by a cladogram, which includes ticks, represented by both soft ticks (e.g., *Ornithodoros* spp.) and hard ticks - the *Ixodes* genus and other genera (i.e., the Metastriata). The analysis was conducted with AvP: the starting data sets were protein sequences obtained from the complete genomes of the tick *I. ricinus*. HGT was determined based on taxonomic assignment of blastp hits and compared hit scores. An “Exclusion Group (EGP)” was defined to encompass ticks and other Acari, whereas the ingroup was defined as Arthropoda. These parameters allowed us to detect HGT from distant organisms (e.g. other Metazoa, bacteria, and viruses) into tick genomes, or into the common ancestor of ticks and other Acari.

**A.**

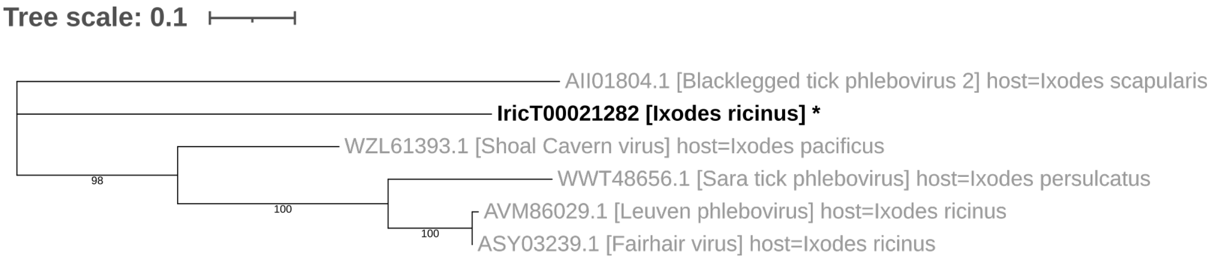
Tree scale: 0.1
AII01804.1 [Blacklegged tick phlebovirus 2] host=Ixodes scapularis
IricT00021282 [Ixodes ricinus] *
WZL61393.1 [Shoal Cavern virus] host=Ixodes pacificus
WWT48656.1 [Sara tick phlebovirus] host=Ixodes persulcatus
AVM86029.1 [Leuven phlebovirus] host=Ixodes ricinus
ASY03239.1 [Fairhair virus] host=Ixodes ricinus
98
100
100

**B.**

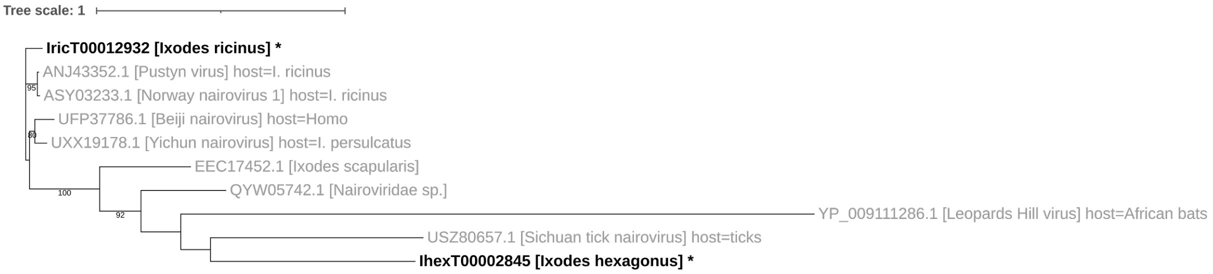
Tree scale: 1
IricT00012932 [Ixodes ricinus] *
ANJ43352.1 [Pustyn virus] host=I. ricinus
ASY03233.1 [Norway nairovirus 1] host=I. ricinus
UFP37786.1 [Beiji nairovirus] host=Homo
UXX19178.1 [Yichun nairovirus] host=I. persulcatus
EEC17452.1 [Ixodes scapularis]
QYW05742.1 [Nairoviridae sp.]
YP_009111286.1 [Leopards Hill virus] host=African bats
USZ80657.1 [Sichuan tick nairovirus] host=ticks
IhexT00002845 [Ixodes hexagonus] *
95
100
92

**C.**

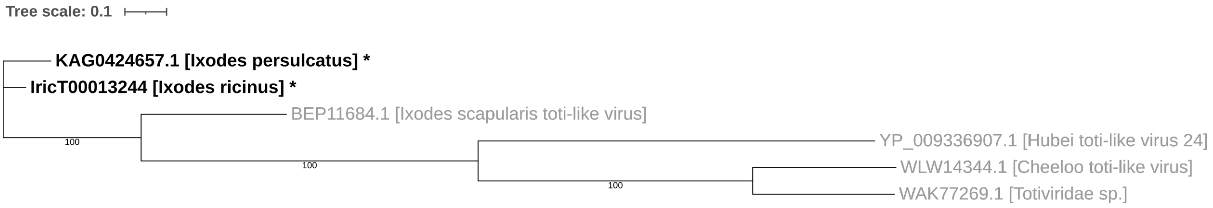
Tree scale: 0.1
KAG0424657.1 [Ixodes persulcatus] *
IricT00013244 [Ixodes ricinus] *
BEP11684.1 [Ixodes scapularis toti-like virus]
YP_009336907.1 [Hubei toti-like virus 24]
WLW14344.1 [Cheeloo toti-like virus]
WAK77269.1 [Totiviridae sp.]
100
100
100

**D.**

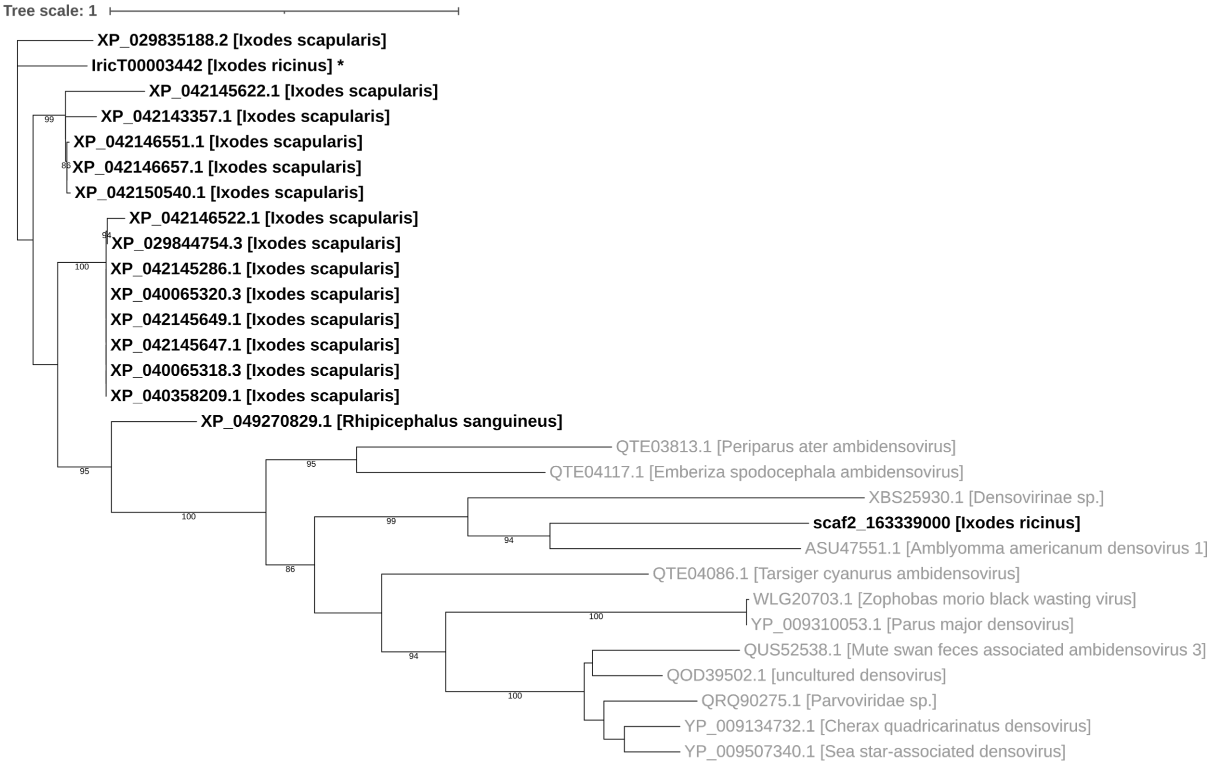
Tree scale: 1
XP_029835188.2 [Ixodes scapularis]
IricT00003442 [Ixodes ricinus] *
XP_042145622.1 [Ixodes scapularis]
XP_042143357.1 [Ixodes scapularis]
XP_042146551.1 [Ixodes scapularis]
XP_042146657.1 [Ixodes scapularis]
XP_042150540.1 [Ixodes scapularis]
XP_042146522.1 [Ixodes scapularis]
XP_029844754.3 [Ixodes scapularis]
XP_042145286.1 [Ixodes scapularis]
XP_040065320.3 [Ixodes scapularis]
XP_042145649.1 [Ixodes scapularis]
XP_042145647.1 [Ixodes scapularis]
XP_040065318.3 [Ixodes scapularis]
XP_040358209.1 [Ixodes scapularis]
XP_049270829.1 [Rhipicephalus sanguineus]
QTE03813.1 [Periparus ater ambidensovirus]
QTE04117.1 [Emberiza spodocephala ambidensovirus]
XBS25930.1 [Densovirinae sp.]
scaf2_163339000 [Ixodes ricinus]
ASU47551.1 [Amblyomma americanum densovirus 1]
QTE04086.1 [Tarsiger cyanurus ambidensovirus]
WLG20703.1 [Zophobas morio black wasting virus]
YP_009310053.1 [Parus major densovirus]
QUS52538.1 [Mute swan feces associated ambidensovirus 3]
QOD39502.1 [uncultured densovirus]
QRQ90275.1 [Parvoviridae sp.]
YP_009134732.1 [Cherax quadricarinatus densovirus]
YP_009507340.1 [Sea star-associated densovirus]
99
100
95
95
100
99
94
86
100
94
100

**E.**

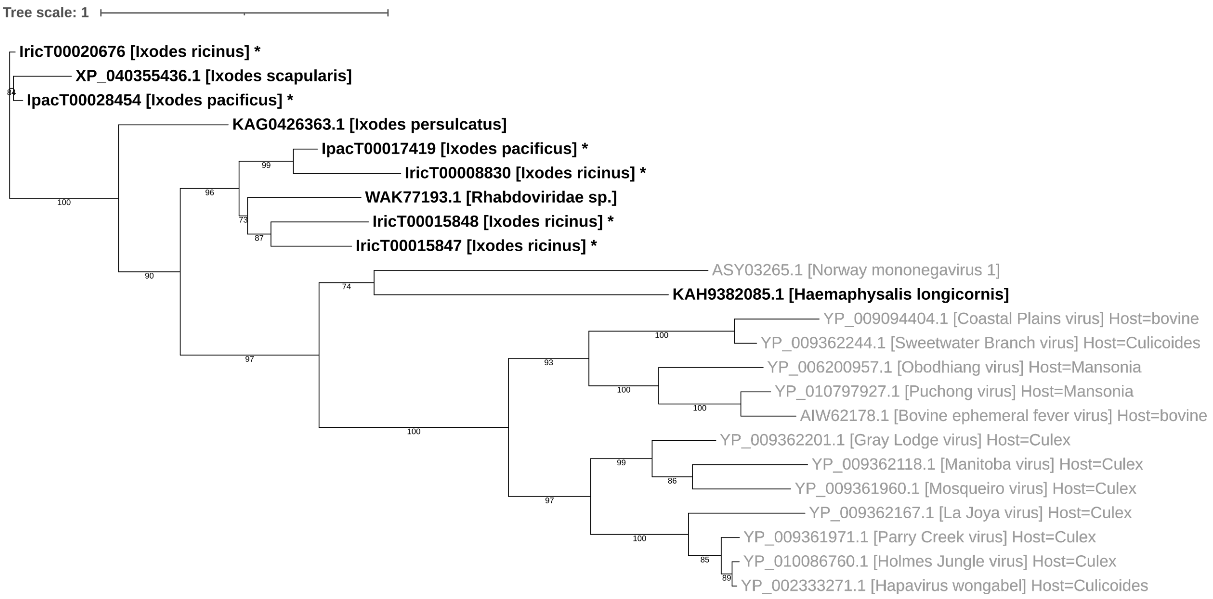


**Figure S2. Genes of viral origin in ticks identified as acquired through HGT.**

Phylogenetic trees include horizontally transferred viral sequences in the genomes of *I. ricinus* and homolog genes in other tick genomes. Shown in light grey are sequences described as tick-associated viruses (note that they may represent endogenous viral elements or EVEs). Bootstrap values are indicated at the nodes.

**A. Phlebovirus-like sequences (Bunyavirales; Phenuiviridae):** Five sequences from *I. ricinus* (IricT0001621, IricT00012740, IricT0015853, IricT00021282, IricT00021393) matched the nucleocapsid protein of phleboviruses. Only the longest sequence, IricT00021282, was retained in our phylogeny, but the other sequences were very similar. All five genes exhibit extremely low expression levels and likely represent degraded genetic elements.

**B. Nairovirus-like sequences (Bunyaviridales, Nairoviridae):** Three sequences (IricT00015852, IricT00008813, IricT00012932) from the *I. ricinus* genome closely matched the nucleocapsid protein sequences of Orthonairoviruses found in tick viromes [63] or vertebrate hosts, including humans [64]. For phylogenetic analysis, we retained only IricT00012932, as it was the longest and most complete among *I. ricinus* sequences. RNA-Seq profiling indicates that IricT00012932 is expressed, particularly in the eggs and in the synganglion. Whereas the other two partial sequences may represent degraded gene copies, the complete sequence with an intact ORF and substantial expression of IricT00012932 suggests functionality and potentially a new role in ticks.

**C. Totitivirus-like sequences (Ghabrivirales):** A sequence from the genomes of both *I. ricinus* (IricT00013244) and *I. persulcatus* (Jia et al. 2020) showed similarity to totiviruses associated with ticks[65]. The sequence from *I. ricinus* appears to have a complete open reading frame (ORF) but exhibits a very low level of expression.

**D. Densovirus-like sequences (Piccovirales):** A sequence from the *I. ricinus* genome (IricT00003442) was similar to non-structural proteins from Densovirinae (Parvoviridae). Multiple similar copies exist in the genome of *I. scapularis*. A second sequence, initially unannotated, was found in the genome of *I. ricinus* (scaf2_163339000): it is an apparently intact and complete ORF, but its sequence diverges significantly from IricT0003442, suggesting an independent origin and separate inclusion in the tick genome. Both *I. ricinus* sequences have negligible expression levels based on RNA-Seq data.

**E. Rhabdovirus-like sequences (Mononegavirales; Rhabdoviridae)** In total, four sequences from the *I. ricinus* genome (IricT00015847, IricT000015848 and IricT00020876, IricT00008830) and one from the *I. pacificus* genome (IpacT000017419) were identified as HGT from viral donors and assigned to rhabdoviruses. These sequences are similar to *I. scapularis* EVEs described in [15] and the Norway mononegavirus 1 described in [66]. The expression level of the *I. ricinus* sequences is negligible, except for IricT00020876 which is expressed at low levels, mostly in the synganglion and the ovary.

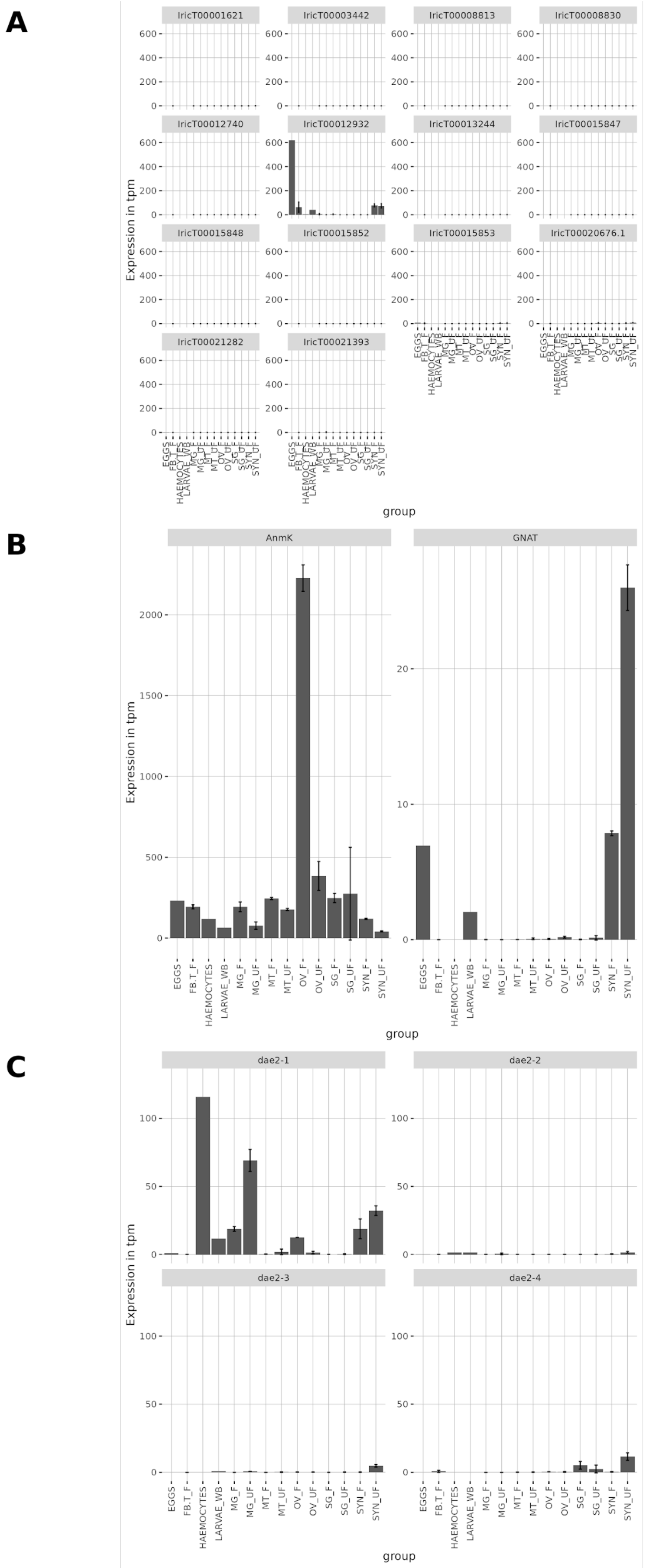
A
IricT00001621
IricT00003442
IricT00008813
IricT00008830
IricT00012740
IricT00012932
IricT00013244
IricT00015847
IricT00015848
IricT00015852
IricT00015853
IricT00020676.1
IricT00021282
IricT00021393
Expression in tpm
group
B
AnmK
GNAT
Expression in tpm
EGGS
FB.T_F
HAEMOCYTES
LARVAE_WB
MG_F
MG_UF
MT_F
MT_UF
OV_F
OV_UF
SG_F
SG_UF
SYN_F
SYN_UF
group
C
dae2-1
dae2-2
dae2-3
dae2-4
Expression in tpm
group

**Figure S3. Expression-levels in various stages and tissues of candidate HGT genes in *I. ricinus*.**

Expressions levels are normalized read counts taken from an atlas of expression based on RNA-Seq data[31]. Counts are in Transcripts per Million (TPM), for the following tissues/stages: eggs, Fat-body/trachea of fed ticks (FB-T_F), hemocytes, larvae (whole bodies, LARVAE_WB), midgut of unfed females (MG_UF) or of fed females (MG_F), malpighian tubules of fed females (MT_F) or unfed females (MT_UF), ovaries of fed females (OV_F) or unfed females (OV_UF), salivary glands of fed females (SG_F) or unfed females (SG_UF), synganglion of fed females (SYN_F) or unfed females (SYN_UF). **A:** Genes of viral origin - panels are at the same scale. **B:** Novel cases of genes of bacterial origin: *anmK (*left) and *GNAT* (right). **C:** *dae2-1* to *dae2-4* (other tick genes of bacterial origin) - panels are at the same scale. Gene Accessions in GenBank are given in (**Table S1**).

**A**

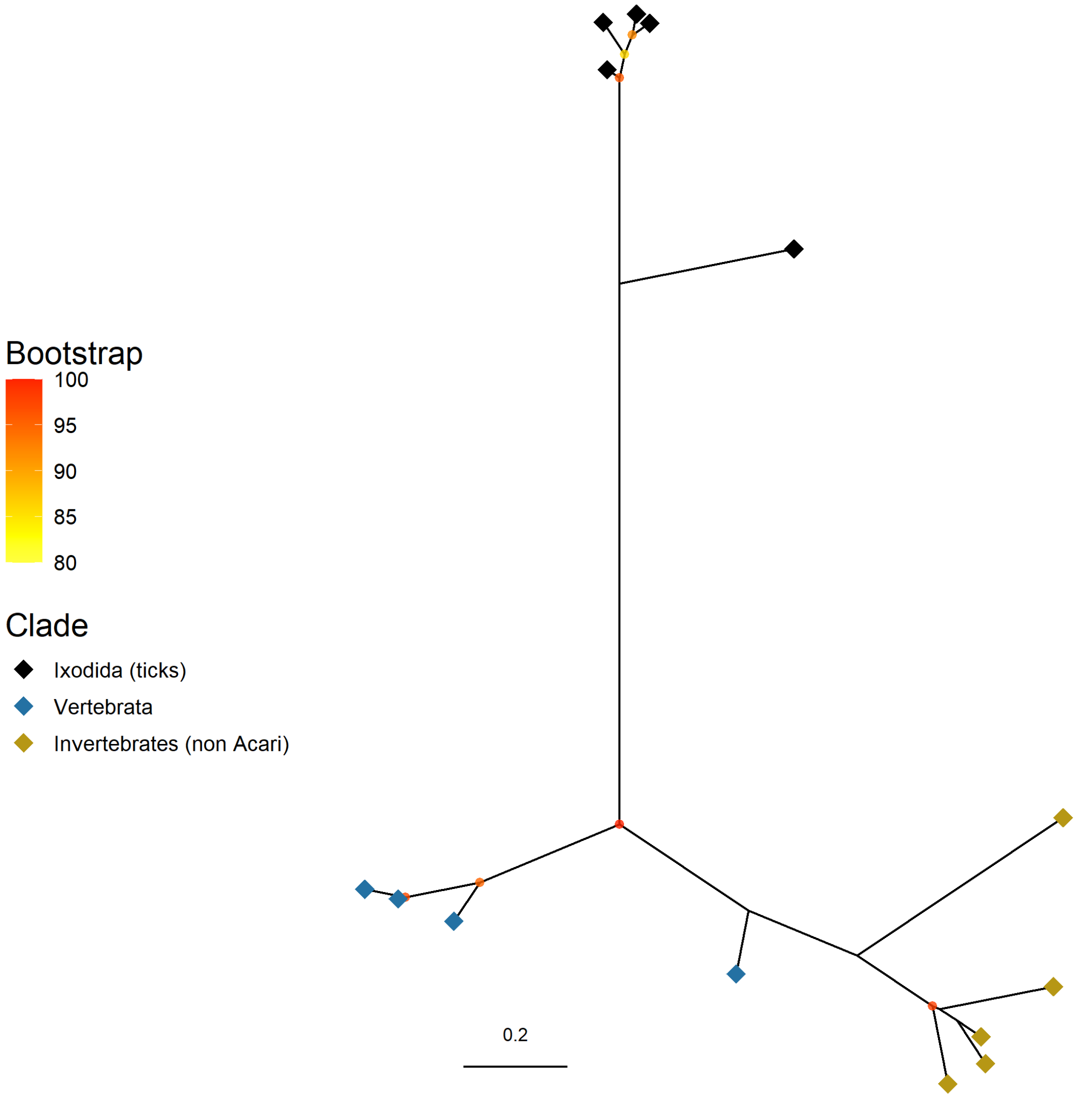

Bootstrap
100
95
90
85
80
Clade
Ixodida (ticks)
Vertebrata
Invertebrates (non Acari)
0.2

**B**

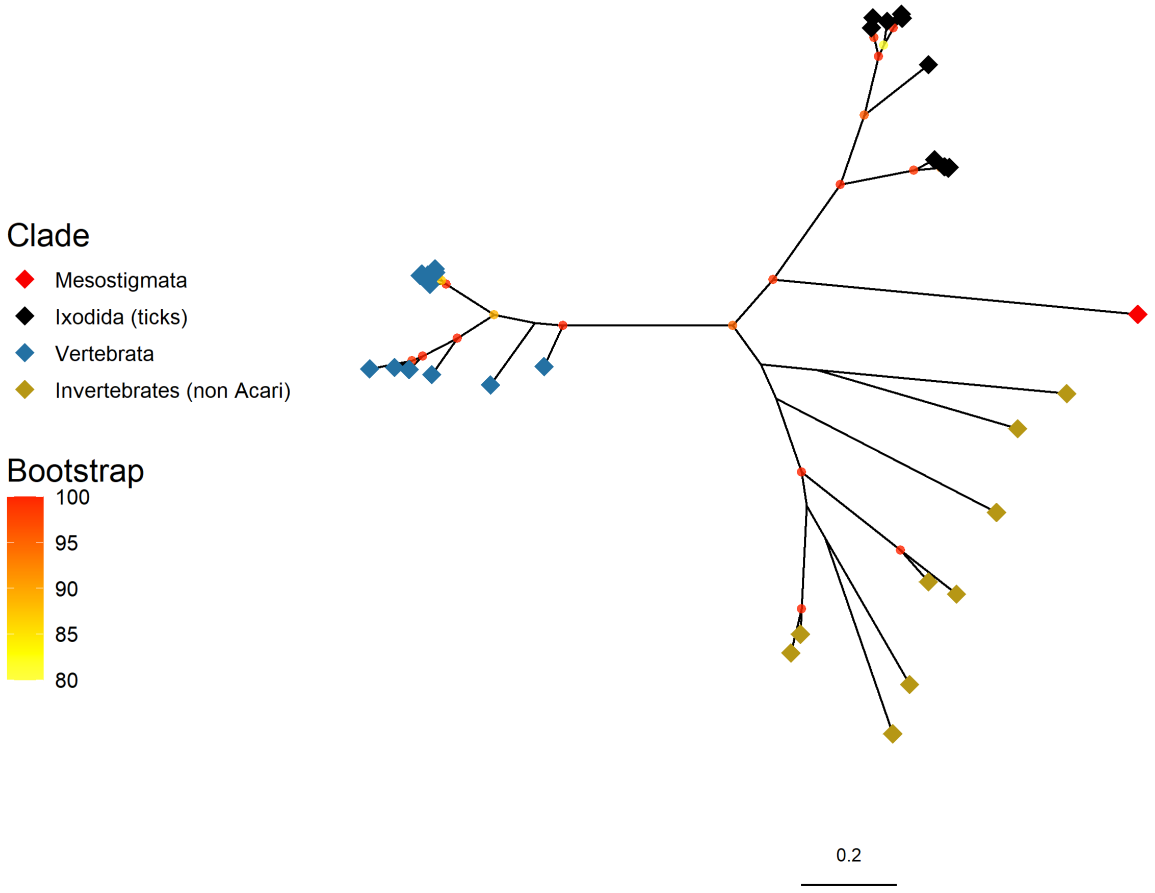
Clade
Mesostigmata
Ixodida (ticks)
Vertebrata
Invertebrates (non Acari)
Bootstrap
100
95
90
85
80
0.2

**C**

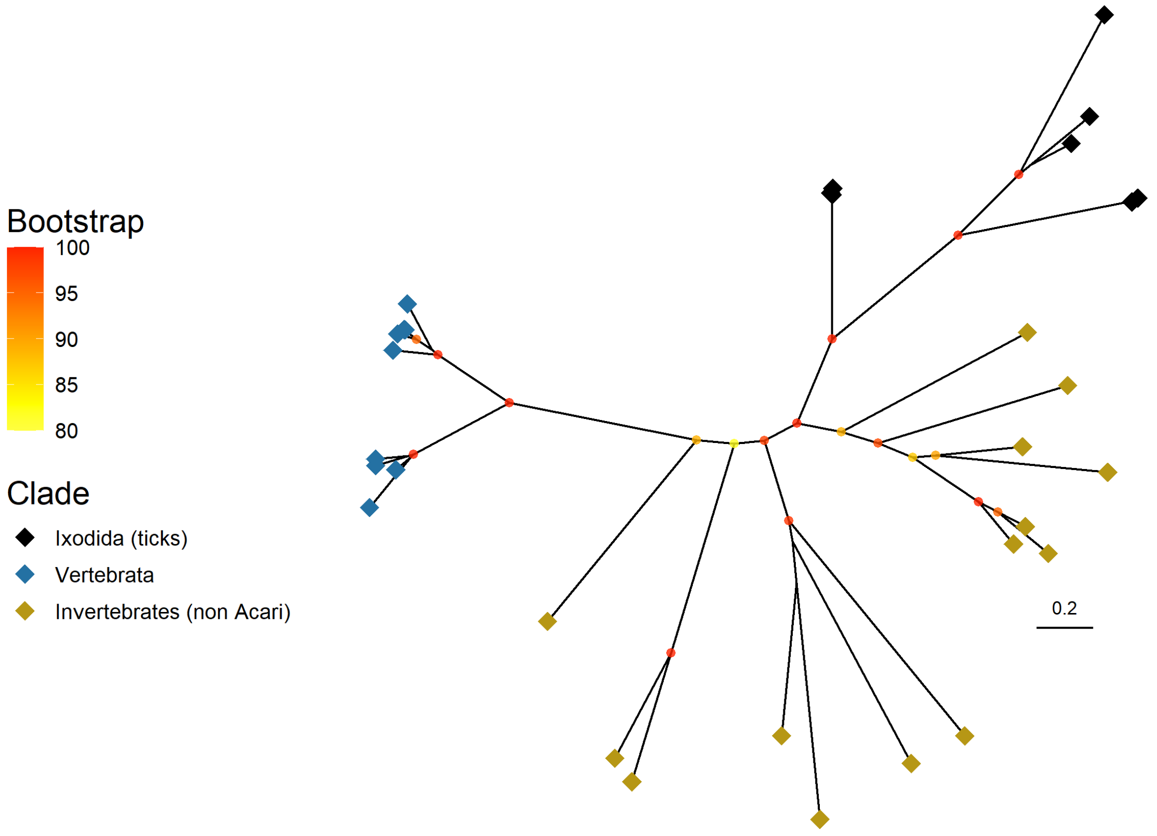


**Figure S4. Lack of phylogenetic support for horizontal transfers from vertebrates to ticks.**

For three candidates identified by the AvP software, we searched homologs in vertebrates, ticks, and other “Invertebrates” (Protostomes, excluding Acari) respectively. Best hits were recorded, gene collections were cleaned to suppress partial or redundant sequences, and a ML tree was built. Sequences are represented by a diamond colored according to their taxonomic group: vertebrates (blue), ticks (black), “invertebrates” (green). Panels correspond to three different HGT candidates identified by AvP in *I. ricinus* **A** IricT00015509 - the *I. ricinus* gene IricT00015509 encodes a Translocator protein which has homologs in ticks, vertebrates, and spiders. The tree shows that a tick clade was separated from all other sequences by a very long branch and did not group robustly with the vertebrate sequences clade. **B** IricT00013518. **C** IricT00019949 - because the gene model for IricT00019949 was incomplete, we rather used homologs of this sequence in *I. scapularis*; these homologs constitute a whole gene family (n=8 genes), annotated as “calcium activated chlorine channels” - For both **(B)** and **(C)**, the differences in BLASTP scores and identities between tick genes and vertebrate or non-tick arthropod hits were minimal, indicating very low conservation. Overall, these phylogenetic analyses do not support HGT.

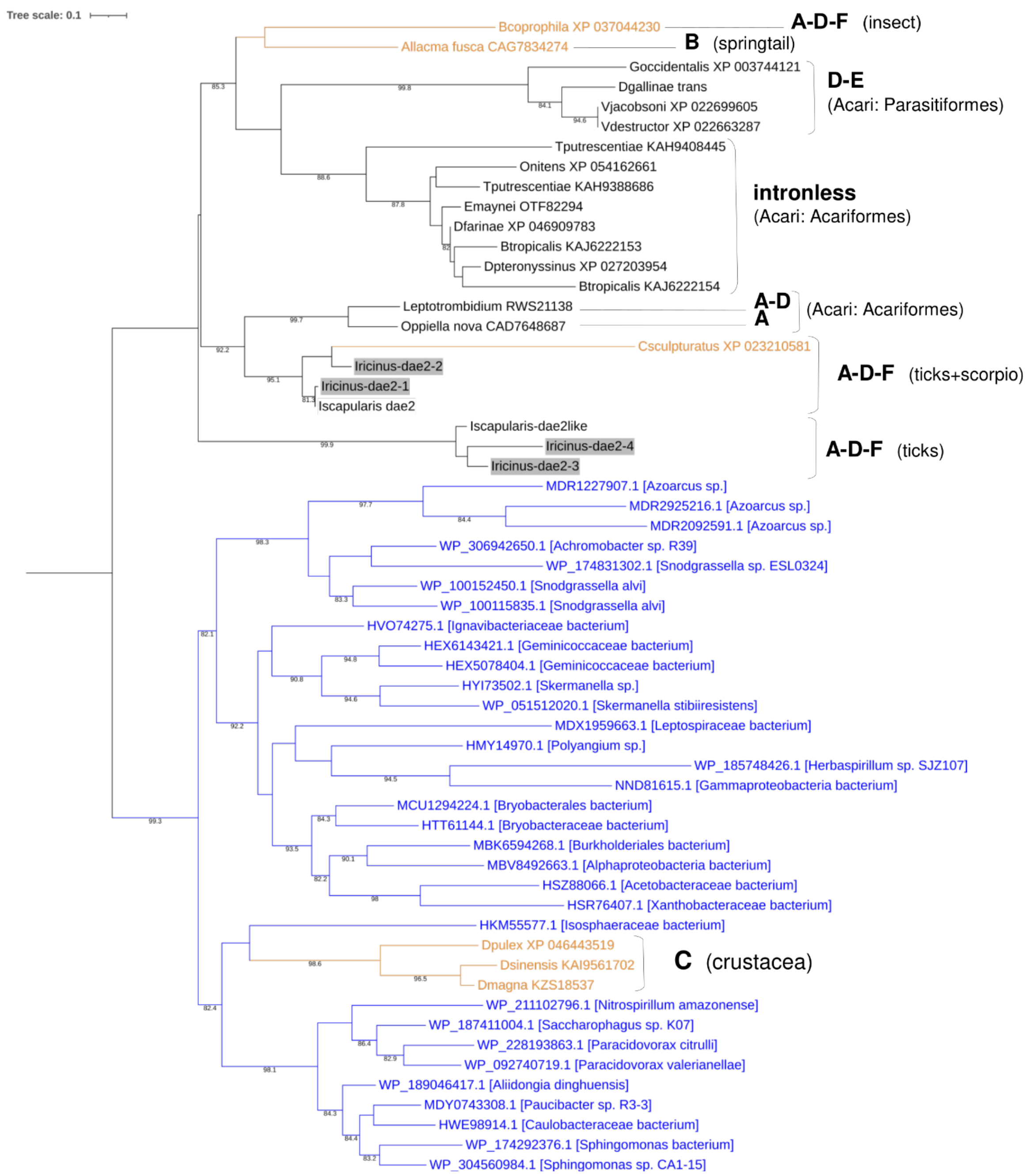
Tree scale: 0.1
Bcoprophila XP 037044230
A-D-F (insect)
Allacma fusca CAG7834274
B (springtail)
Goccidentalis XP 003744121
Dgallinae trans
Vjacobsoni XP 022699605
Vdestructor XP 022663287
D-E
(Acari: Parasitiformes)
Tputrescentiae KAH9408445
Onitens XP 054162661
Tputrescentiae KAH9388686
Emaynei OTF82294
Dfarinae XP 046909783
Btropicalis KAJ6222153
Dpteronyssinus XP 027203954
Btropicalis KAJ6222154
intronless
(Acari: Acariformes)
Leptotrombidium RWS21138
Oppiella nova CAD7648687
A-D
A
(Acari: Acariformes)
Csculpturatus XP 023210581
Iricinus-dae2-2
Iricinus-dae2-1
Iscapularis dae2
A-D-F (ticks+scorpio)
Iscapularis-dae2like
Iricinus-dae2-4
Iricinus-dae2-3
A-D-F (ticks)
MDR1227907.1 [Azoarcus sp.]
MDR2925216.1 [Azoarcus sp.]
MDR2092591.1 [Azoarcus sp.]
WP_306942650.1 [Achromobacter sp. R39]
WP_174831302.1 [Snodgrassella sp. ESL0324]
WP_100152450.1 [Snodgrassella alvi]
WP_100115835.1 [Snodgrassella alvi]
HVO74275.1 [Ignavibacteriaceae bacterium]
HEX6143421.1 [Geminicoccaceae bacterium]
HEX5078404.1 [Geminicoccaceae bacterium]
HYI73502.1 [Skermanella sp.]
WP_051512020.1 [Skermanella stibiiresistens]
MDX1959663.1 [Leptospiraceae bacterium]
HMY14970.1 [Polyangium sp.]
WP_185748426.1 [Herbaspirillum sp. SJZ107]
NND81615.1 [Gammaproteobacteria bacterium]
MCU1294224.1 [Bryobacterales bacterium]
HTT61144.1 [Bryobacteraceae bacterium]
MBK6594268.1 [Burkholderiales bacterium]
MBV8492663.1 [Alphaproteobacteria bacterium]
HSZ88066.1 [Acetobacteraceae bacterium]
HSR76407.1 [Xanthobacteraceae bacterium]
HKM55577.1 [Isosphaeraceae bacterium]
Dpulex XP 046443519
Dsinensis KAI9561702
Dmagna KZS18537
C (crustacea)
WP_211102796.1 [Nitrospirillum amazonense]
WP_187411004.1 [Saccharophagus sp. K07]
WP_228193863.1 [Paracidovorax citrulli]
WP_092740719.1 [Paracidovorax valerianellae]
WP_189046417.1 [Aliidongia dinghuensis]
MDY0743308.1 [Paucibacter sp. R3-3]
HWE98914.1 [Caulobacteraceae bacterium]
WP_174292376.1 [Sphingomonas bacterium]
WP_304560984.1 [Sphingomonas sp. CA1-15]

**Figure S5. Evolution of the tae2-like gene transferred from bacteria to ticks and other Acari.**

Maximum Likelihood phylogenetic tree of *tae2-dae2*. In their study, *Chou et al.* 2015 identified a gene of bacterial origin in ticks (*I. scapularis*) and other Acari, called *dae2*. Color labels indicate *dae2* homologs in different taxa as follow: bacteria (*tae2*, in blue), ticks and other Acari (black), other arthropods (orange). The tree indicates multiple copies similar to *dae2* in ticks (two in *I. scapularis*, four in *I. ricinus*), suggesting lineage-specific duplications. Next to labels, we report patterns of splice sites (sites A to F) based on each gene model and the combined alignment of sequences.

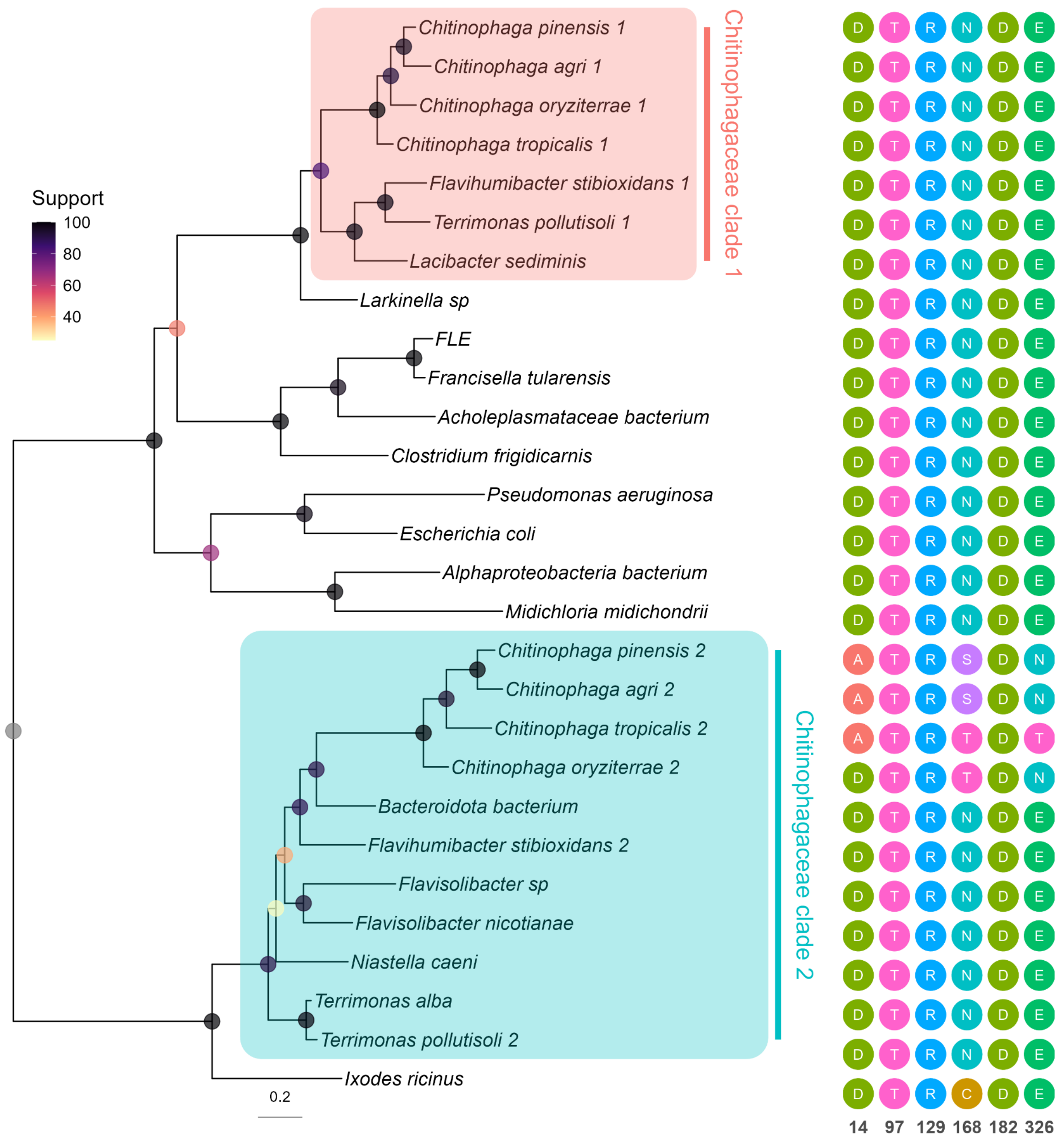


**Figure S6. Maximum-likelihood phylogenetic tree of anmK including sequences from several bacterial clades and from tick-associated endosymbionts.**
Tick endosymbionts retaining *anmK* and thus present in this tree include *Francisella*-like endosymbionts (FLE) and *M. mitochondrii*. Tick sequences (represented here by *Ixodes ricinus*) are closely related to sequences from the *Chitinophagaceae*, a bacterial genus which typically possesses two copies of this gene, forming two clades. Tick sequences cluster with clade 2, the more divergent of the two. Important functional sites reported in the literature are indicated next to each sequence, using the *P. aeruginosa* residue numbering (including the catalytic site D182, which is conserved across all sequences). Each amino acid is associated with a specific color, to highlight substitutions. Genbank accessions for each label are indicated between brackets: Terrimonas_pollutisoli_2 (WP_276500879), Bacteroidota_bacterium (MEP6750283.1), Niastella_caeni (WP_136575791), Flavisolibacter_nicotianae (WP_121352354.1), Flavihumibacter_stibioxidans_2 (WP_187255858), Chitinophaga_agri_2 (A0A6B9ZLU2), Chitinophaga_pinensis_2 (WP_146305114),

Chitinophaga_tropicalis_2 (A0A7K1TZK2_9BACT), Chitinophaga_oryziterrae_2 ( A0A6N8JEQ5_9BACT), Lacibacter_sediminis (WP_182805696), Flavihumibacter_stibioxidans_1 (WP_187256798), Terrimonas_pollutisoli_1 (WP_276500604.1), Chitinophaga_tropicalis_1 (WP_157308152.1), Terrimonas_alba (WP_385894194.1), Chitinophaga_agri_1 (WP_162332241.1), Chitinophaga_pinensis_1 (WP_146303413), Chitinophaga_oryziterrae_1 (WP_157298141.1), Flavisolibacter_sp. (HEU0065299.1), Larkinella_sp (WP_421828228), Escherichia_coli (WP_065226992.1), Pseudomonas_aeruginosa (WP_034069018), FLE (WP_143385200.1), Francisella_tularensis (WP_003034536.1), Alphaproteobacteria_bacterium (MCE9508715.1), Midichloria_midichondrii (AEI89037), Ixodes_ricinus (CAN7959379.1), Clostridium_frigidicarnis (WP_207647677.1), Acholeplasmataceae_bacterium (HBY64991.1).

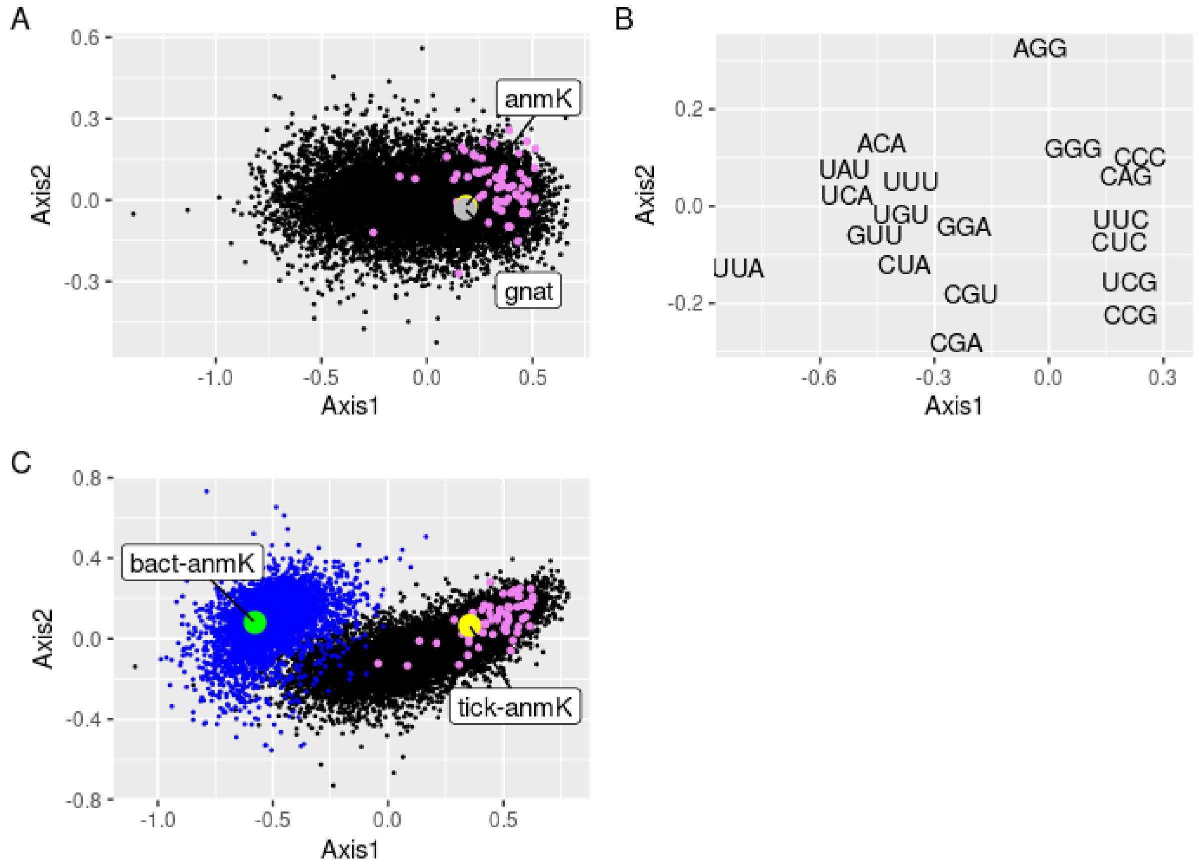


**Figure S7. Codon usage analysis of tick genes indicates codon adaptation of tick *anmK***

Factorial correspondence analysis on Relative Synonymous Codon Usage (RSCU), performed with CodonW. **A.** Distribution of *I. ricinus* genes on the first two axes, for the coding sequences of the tick *I. ricinus*. Dots in purple correspond to ribosomal protein genes, expected to have an optimized codon usage. The position of two tick genes horizontally transferred from a bacterial donor, *GNAT* (grey) and *anmK* (yellow), is shown. Relative inertias of the first two axes are 0.272 (Axis 1) and 0.033 (Axis 2). **B.** Position of the variables (codons) in the *I. ricinus* analysis of RSCU. showing that optimal codon usage is associated with G- and C-ending codons. **C.** Codon usage analysis on RSCU, combining the *I. ricinus* genome (black, purple and yellow) and the bacterium *Terrimonas* spp. (GCF_003075435.1; blue and green), representing the donor group of the horizontally transferred *anmK*. The positions of *anmK* genes from both genomes, tick and bacteria, are shown. Ribosomal proteins of *I. ricinus* are shown in purple Relative inertias of the first two axes are 0.365 (Axis 1) and 0.048 (Axis 2).

**Consensus**

1. CfAnmK WP_207647677.1
2. AbAnmK HBY64991.1
3. FeAnmK WP_143385200.1
4. FtAnmK WP_003034536.1
5. ApAnmK MCE9508715.1
6. MmAnmK AEI89037
7. PaAnmK AAG04055.1
8. EcAnmK AAC74712.1
9. LxAnmK WP_421828228.1
10. LsAnmK WP_182805696.1
11. FsAnmK WP_187256798.1
12. Tp_AnmK1 WP_276500604.1
13. CtAnmK1 WP_157308152.1
14. CoAnmK1 WP_157298141.1
15. CpAnmK2 ACU58695.1
16. CaAnmK2 WP_162333750.1
17. FbAnmK HEU0065299.1
18. FnAnmK WP_121352354.1
19. NcAnmK WP_136575791.1
20. TaAnmK WP_385894194.1
21. TpAnmK2 WP_276500879.1
22. FsAnmK2 WP_187255858.1
23. BbAnmK MEP6750283.1
24. CoAnmK2 WP_157302509.1
25. CtAnmK2 WP_157304936.1
26. CpAnmK1 WP_146303413.1
27. CaAnmK1 WP_162332241.1
28. OmAnmK MBZ3971232.1
29. RsAnmK XP_037520253.1
30. IrAnmK CAN7959379.1

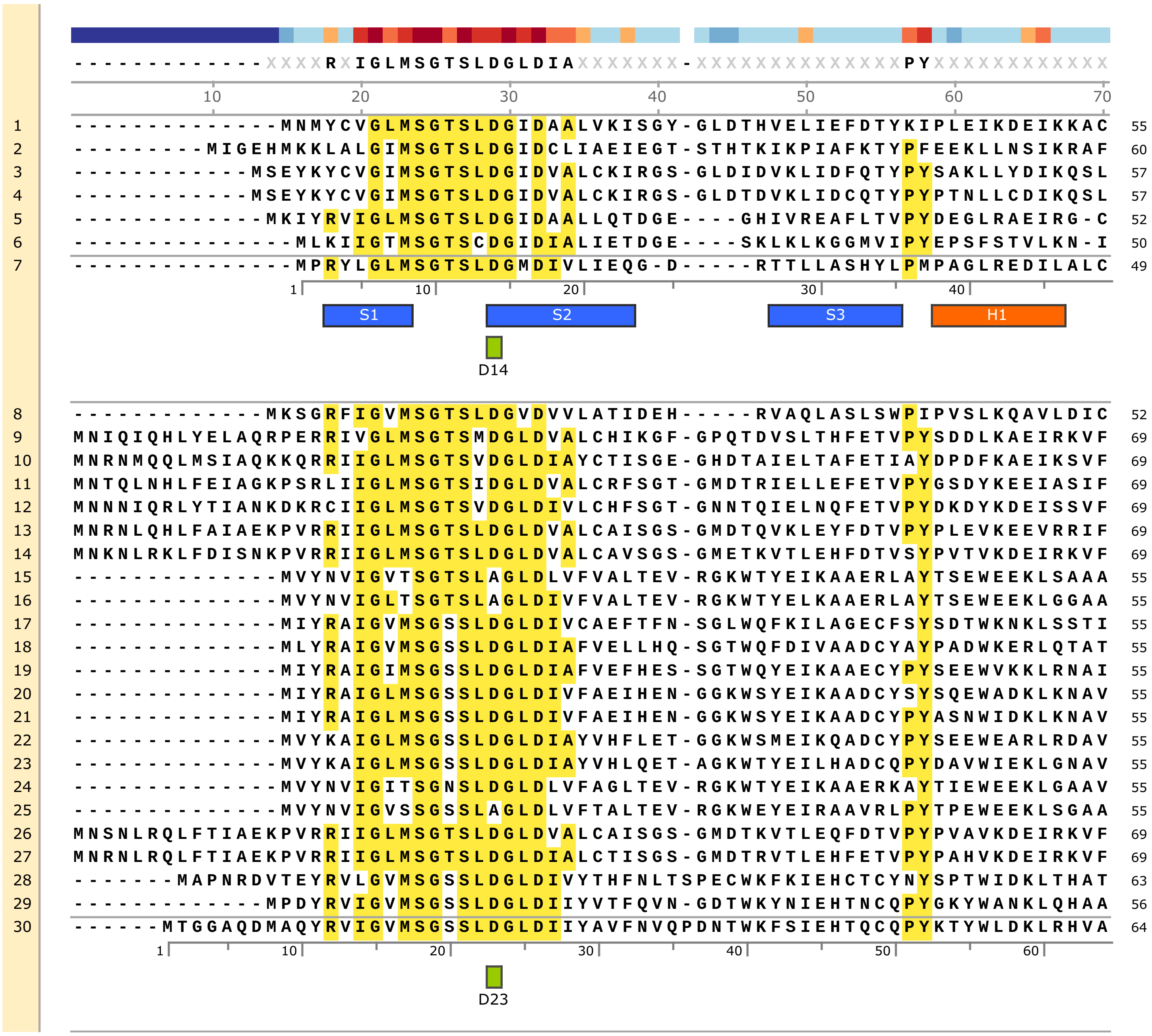

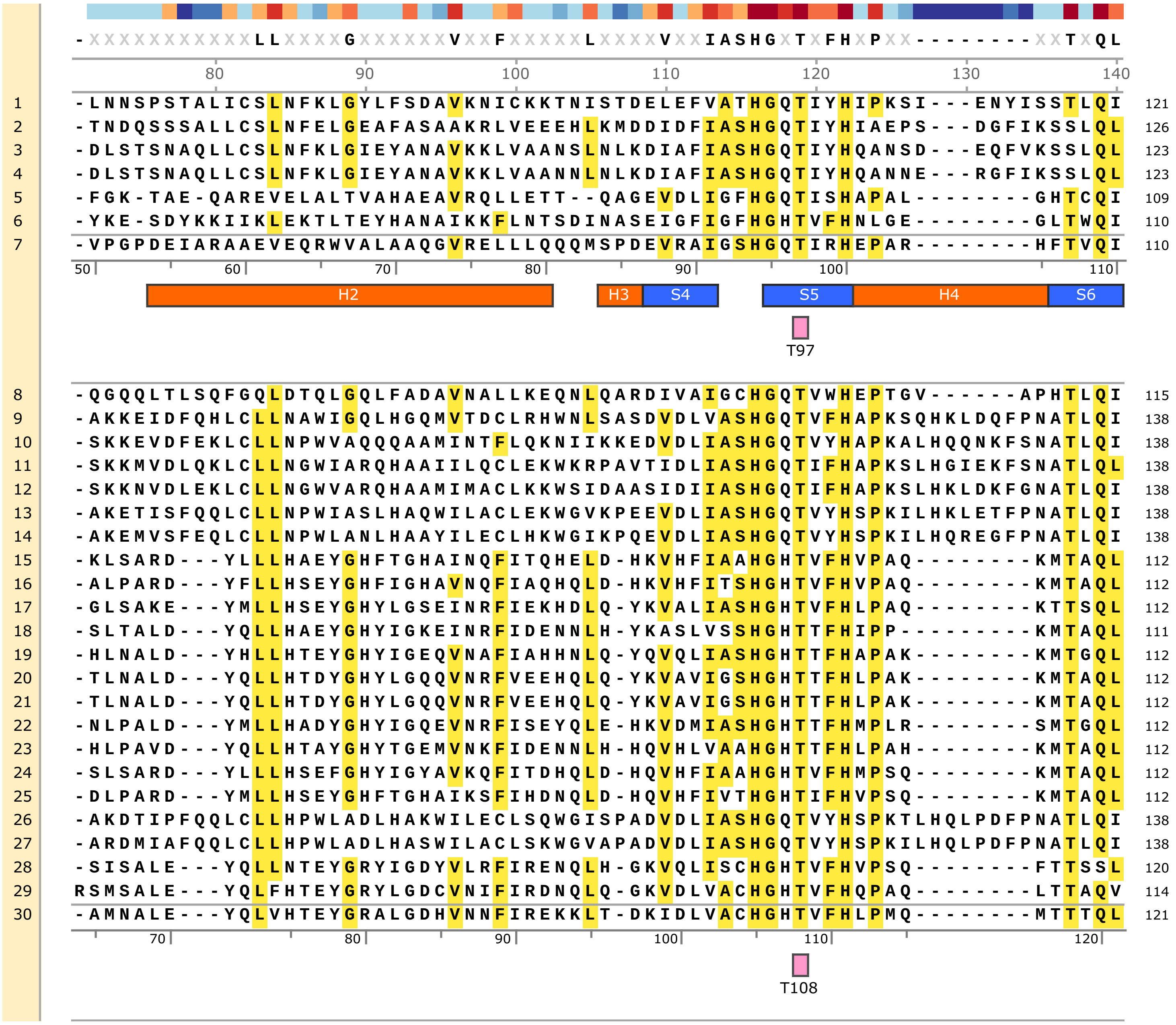

-XXXXXXXXXXXLLXXXXGXXXXXXVXXFXXXXXLXXXXVXXIASHGXTXFHXPXX--------XXTXQL
80 90 100 110 120 130 140
1 -LNNSPSTALICSLNFKLGYLFSDAVKNICKKTNISTDELEFVATHGQTIYHIPKSI---ENYISSTLQI 121
2 -TNDQSSSALLCSLNFELGEAFASAAKRLVEEEHLKMDDIDFIASHGQTIYHIAEPS---DGFIKSSLQL 126
3 -DLSTSNAQLLCSLNFKLGIEYANAVKKLVAANSLNLKDIAFIASHGQTIYHQANSD---EQFVKSSLQL 123
4 -DLSTSNAQLLCSLNFKLGIEYANAVKKLVAANNLNLKDIAFIASHGQTIYHQANNE---RGFIKSSLQL 123
5 -FGK-TAE-QAREVELALTVAHAEAVRQLLETT--QAGEVDLIGFHGQTISHAPAL--------GHTCQI 109
6 -YKE-SDYKKIIKLEKTLTEYHANAIKKFLNTSDINASEIGFIGFHGHTVFHNLGE--------GLTWQI 110
7 -VPGPDEIARAAEVEQRWVALAAQGVRELLLQQQMSPDEVRAIGSHGQTIRHEPAR--------HFTVQI 110
50 60 70 80 90 100 110
H2
H3
S4
S5
H4
S6
T97
8 -QGQQLTLSQFGQLDTQLGQLFADAVNALLKEQNLQARDIVAIGCHGQTVWHEPTGV------APHTLQI 115
9 -AKKEIDFQHLCLLNAWIGQLHGQMVTDCLRHWNLSASDVDLVASHGQTVFHAPKSQHKLDQFPNATLQI 138
10 -SKKEVDFEKLCLLNPWVAQQQAAMINTFLQKNIIKKEDVDLIASHGQTVYHAPKALHQQNKFSNATLQI 138
11 -SKKMVDLQKLCLLNGWIARQHAAIILQCLEKWKRPAVTIDLIASHGQTIFHAPKSLHGIEKFSNATLQL 138
12 -SKKNVDLEKLCLLNGWVARQHAAMIMACLKKWSIDAASIDIIASHGQTIFHAPKSLHKLDKFGNATLQI 138
13 -AKETISFQQLCLLNPWIASLHAQWILACLEKWGVKPEEVDLIASHGQTVYHSPKILHKLETFPNATLQI 138
14 -AKEMVSFEQLCLLNPWLANLHAAYILECLHKWGIKPQEVDLIASHGQTVYHSPKILHQREGFPNATLQI 138
15 -KLSARD---YLLLHAEYGHFTGHAINQFITQHELD-HKVHFIAAHGHTVFHVPAQ--------KMTAQL 112
16 -ALPARD---YFLLHSEYGHFIGHAVNQFIAQHQLD-HKVHFITSHGHTVFHVPAQ--------KMTAQL 112
17 -GLSAKE---YMLLHSEYGHYLGSEINRFIEKHDLQ-YKVALIASHGHTVFHLPAQ--------KTTSQL 112
18 -SLTALD---YQLLHAEYGHYIGKEINRFIDENNLH-YKASLVSSHGHTTFHIPP---------KMTAQL 111
19 -HLNALD---YHLLHTEYGHYIGEQVNAFIAHHNLQ-YQVQLIASHGHTTFHAPAK--------KMTGQL 112
20 -TLNALD---YQLLHTDYGHYLGQQVNRFVEEHQLQ-YKVAVIGSHGHTTFHLPAK--------KMTAQL 112
21 -TLNALD---YQLLHTDYGHYLGQQVNRFVEEHQLQ-YKVAVIGSHGHTTFHLPAK--------KMTAQL 112
22 -NLPALD---YMLLHADYGHYIGQEVNRFISEYQLE-HKVDMIASHGHTTFHMPLR--------SMTGQL 112
23 -HLPAVD---YQLLHTAYGHYTGEMVNKFIDENNLH-HQVHLVAAHGHTTFHLPAH--------KMTAQL 112
24 -SLSARD---YLLLHSEFGHYIGYAVKQFITDHQLD-HQVHFIAAHGHTVFHMPSQ--------KMTAQL 112
25 -DLPARD---YMLLHSEYGHFTGHAIKSFIHDNQLD-HQVHFIVTHGHTIFHVPSQ--------KMTAQL 112
26 -AKDTIPFQQLCLLHPWLADLHAKWILECLSQWGISPADVDLIASHGQTVYHSPKTLHQLPDFPNATLQI 138
27 -ARDMIAFQQLCLLHPWLADLHASWILACLSKWGVAPADVDLIASHGQTVYHSPKILHQLPDFPNATLQI 138
28 -SISALE---YQLLNTEYGRYIGDYVLRFIRENQLH-GKVQLISCHGHTVFHVPSQ--------FTTSSL 120
29 RSMSALE---YQLFHTEYGRYLGDCVNIFIRDNQLQ-GKVDLVACHGHTVFHQPAQ--------LTTAQV 114
30 -AMNALE---YQLVHTEYGRALGDHVNNFIREKKLT-DKIDLVACHGHTVFHLPMQ--------MTTTQL 121
70 80 90 100 110 120
T108

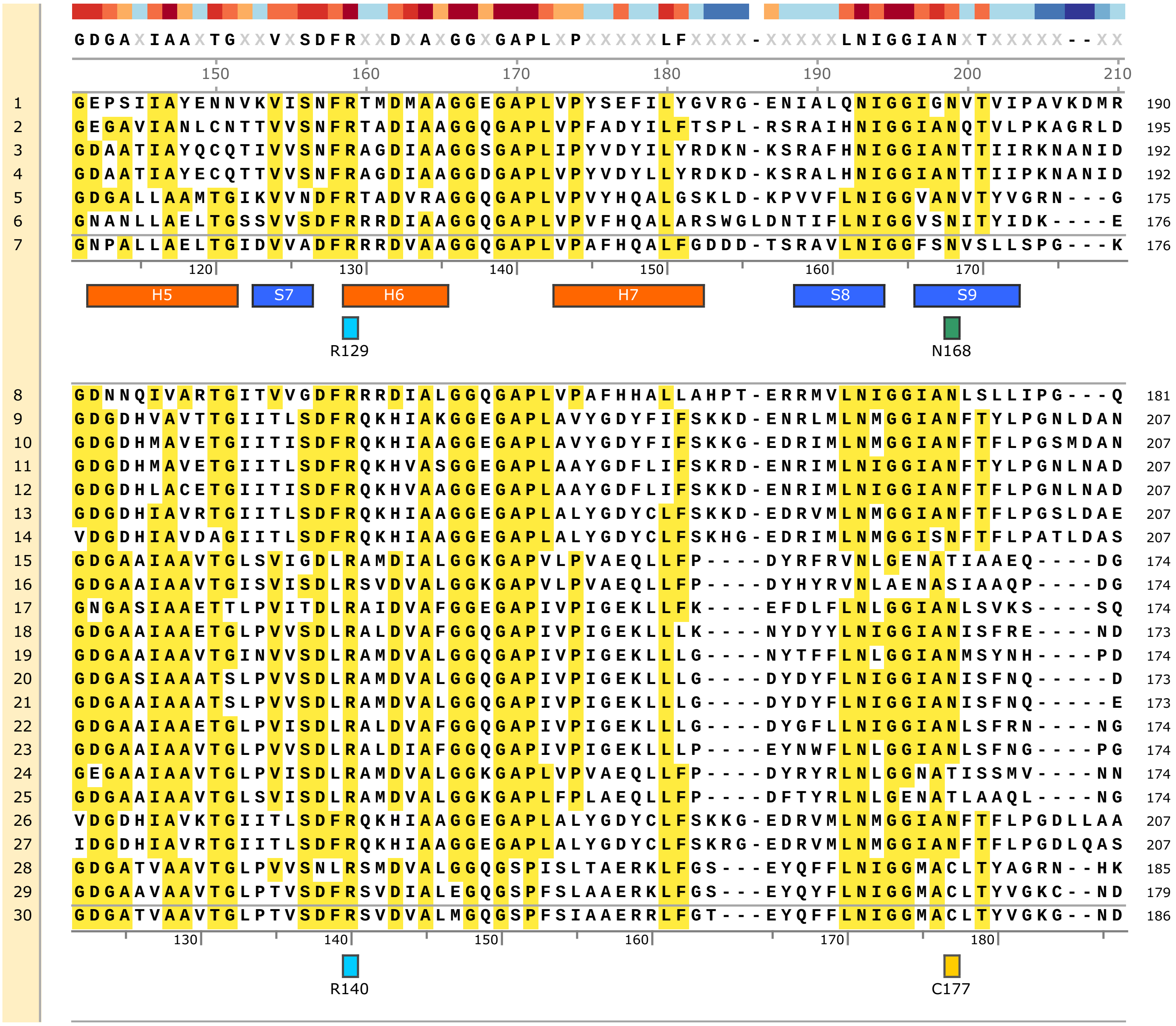
GDGAXIAAXTGXXVXSDFRXXDXAXGGXGAPLXPXXXXXLFXXXX-XXXXXLNIGGIANXTXXXXX--XX
150 160 170 180 190 200 210
1 GEPSIIAYENNVKVISNFRTMDMAAGGEGAPLVPYSEFILYGVRG-ENIALQNIGGIGNVTVIPAVKDMR 190
2 GEGAVIANLCNTTVVSNFRTADIAAGGQGAPLVPFADYILFTSPL-RSRAIHNIGGIANQTVLPKAGRLD 195
3 GDAATIAYQCQTIVVSNFRAGDIAAGGSGAPLIPYVDYILYRDKN-KSRAFHNIGGIANTTIIRKNANID 192
4 GDAATIAYECQTTVVSNFRAGDIAAGGDGAPLVPYVDYLLYRDKD-KSRALHNIGGIANTTIIPKNANID 192
5 GDGALLAAMTGIKVVNDFRTADVRAGGQGAPLVPVYHQALGSKLD-KPVVFLNIGGVANVTYVGRN---G 175
6 GNANLLAELTGSSVVSDFRRRDIAAGGQGAPLVPVFHQALARSWGLDNTIFLNIGGVSNITYIDK----E 176
7 GNPALLAELTGIDVVADFRRRDVAAGGQGAPLVPAFHQALFGDDD-TSRAVLNIGGFSNVSLLSPG---K 176
120 130 140 150 160 170
H5
S7
H6
H7
S8
S9
R129
N168
8 GDNNQIVARTGITVVGDFRRRDIALGGQGAPLVPAFHHALLAHPT-ERRMVLNIGGIANLSLLIPG---Q 181
9 GDGDHVAVTTGIITLSDFRQKHIAKGGEGAPLAVYGDYFIFSKKD-ENRLMLNMGGIANFTYLPGNLDAN 207
10 GDGDHMAVETGIITISDFRQKHIAAGGEGAPLAVYGDYFIFSKKG-EDRIMLNMGGIANFTFLPGSMDAN 207
11 GDGDHMAVETGIITLSDFRQKHVASGGEGAPLAAYGDFLIFSKRD-ENRIMLNIGGIANFTYLPGNLNAD 207
12 GDGDHLACETGIITISDFRQKHVAAGGEGAPLAAYGDFLIFSKKD-ENRIMLNIGGIANFTFLPGNLNAD 207
13 GDGDHIAVRTGIITLSDFRQKHIAAGGEGAPLALYGDYCLFSKKD-EDRVMLNMGGIANFTFLPGSLDAE 207
14 VDGDHIAVDAGIITLSDFRQKHIAAGGEGAPLALYGDYCLFSKHG-EDRIMLNMGGISNFTFLPATLDAS 207
15 GDGAAIAAVTGLSVIGDLRAMDIALGGKGAPVLPVAEQLLFP----DYRFRVNLGENATIAAEQ----DG 174
16 GDGAAIAAVTGISVISDLRSVDVALGGKGAPVLPVAEQLLFP----DYHYRVNLAENASIAAQP----DG 174
17 GNGASIAAETTLPVITDLRAIDVAFGGEGAPIVPIGEKLLFK----EFDLFLNLGGIANLSVKS----SQ 174
18 GDGAAIAAETGLPVVSDLRALDVAFGGQGAPIVPIGEKLLLK----NYDYYLNIGGIANISFRE----ND 173
19 GDGAAIAAVTGINVVSDLRAMDVALGGQGAPIVPIGEKLLLG----NYTFFLNLGGIANMSYNH----PD 174
20 GDGASIAAATSLPVVSDLRAMDVALGGQGAPIVPIGEKLLLG----DYDYFLNIGGIANISFNQ-----D 173
21 GDGAAIAAATSLPVVSDLRAMDVALGGQGAPIVPIGEKLLLG----DYDYFLNIGGIANISFNQ-----E 173
22 GDGAAIAAETGLPVISDLRALDVAFGGQGAPIVPIGEKLLLG----DYGFLLNIGGIANLSFRN----NG 174
23 GDGAAIAAVTGLPVVSDLRALDIAFGGQGAPIVPIGEKLLLP----EYNWFLNLGGIANLSFNG----PG 174
24 GEGAAIAAVTGLPVISDLRAMDVALGGKGAPLVPVAEQLLFP----DYRYRLNLGGNATISSMV----NN 174
25 GDGAAIAAVTGLSVISDLRAMDVALGGKGAPLFPLAEQLLFP----DFTYRLNLGENATLAAQL----NG 174
26 VDGDHIAVKTGIITLSDFRQKHIAAGGEGAPLALYGDYCLFSKKG-EDRVMLNMGGIANFTFLPGDLLAA 207
27 IDGDHIAVRTGIITLSDFRQKHIAAGGEGAPLALYGDYCLFSKRG-EDRVMLNMGGIANFTFLPGDLQAS 207
28 GDGATVAAVTGLPVVSNLRSMDVALGGQGSPISLTAERKLFGS---EYQFFLNIGGMACLTYAGRN--HK 185
29 GDGAAVAAVTGLPTVSDFRSVDIALEGQGSPFSLAAERKLFGS---EYQYFLNIGGMACLTYVGKC--ND 179
30 GDGATVAAVTGLPTVSDFRSVDVALMGQGSPFSIAAERRLFGT---EYQFFLNIGGMACLTYVGKG--ND 186
130 140 150 160 170 180
R140
C177

```
        XXXAFDXGPGNXXLDXXXXXXXGXX-YDXXGXXAXXGXVXXXLLXXLXXXXFXXXXXPKSXGXXXFGXXX

              220       230       240       250       260       270       280

1       EIYAFDTGPGNMMIDEACSKLFGVS-YDENGDIAKSGKLCETLMEELKSHPYIDLKPPKTTGREDFGEQF   259
2       DVFAFDSGPGNMMIDYAMKTLYGKP-FDQNGDIAKSGSIISSMMNELFNNPYFKKIPPKSTGRELFGDSY   264
3       DIYAFDTGPGNMMINRAMEVLFNRD-YDKDGDIAAAGIVIVDMLQELLENPYLKQKPPKSTGRELFGVEF   261
4       DIYAFDTGPGNMMINRAMEVLFNQD-YDKGGDTAAEGIVIVDMLQELLDNPYLKQKPPKSTGRELFGINY   261
5       ELIAFDTGPGNALIDDWMLKKIGHK-FDAGGKISLSGRVDEAVLGQLLGHPFFASPVPKSLDRNDFVSQA   244
6       EIIAFDAGPGNALIDDFLYKRVAIS-YDEGGKIGARGKVDFEVVRTFQKDVFYSGPIPKSLDRNHFNYID   245
7       PVRGFDCGPGNVLMDAWIHHQRGEH-FDRDGAWAASGQVNHALLASLLADEFFAARGPKSTGRERFNLPW   245

          180       190       200       210       220       230       240
          S10       H8            H9          H10                        H11
          D182

8       PVGGYDTGPGNMLMDAWIWRQAGKP-YDKDAEWARAGKVILPLLQNMLSDPYFSQPAPKSTGREYFNYGW   250
9       EVFTTDTGPGNTLIDAFTRQLFNKP-YDADGRIAASGTVNQDLVQMLKKDSFFKAPFPKTTGPEVFNPDY   276
10      KIFSTDTGTGNTLMDALMQQHFLGKYFDEDAAVAKKGTINEALLNALKDHSFFQQSFPKTTGPELFNLHY   277
11      EVFCSDTGTGNTLMDALTRKHFPGSYFDEGAAIAGRGVIHAGLLAALKDHPFFHQPFPKTTGPELFNLAY   277
12      EVFCTDTGPGNTLMDAFVQKHYPGKYFDENASLASLGVINEKLLALIDHSFFHQSFPKTTGPELFNLHY   277
13      GVFVTDTGPGNTLLDAFARKYFQAE-YDNDAAFAKQGTVSEALLAALKDNSFFKEPFPRTTGPELFSIRY   276
14      AVYVTDTGPGNTLLDAFARVYFNEE-YDKDALFAKQGKVSEALLAALKDHPFFKEPFPRTTGPELFNKTY   276
15      QLIAWDVCPANYVLDTLAGM-LGRA-FDEEGKLAAGGVTDQKLLDALNGLAFYNEKYPKTLAA-KFGTGT   241
16      QLIAFDVCPCNYVLDTLAGL-LGRP-FDEEGKLAAGGVTDQGLLDALNGLAFYSEQYPRTLTS-KFGTGT   241
17      RFIAFDVCPANRVLNMITET-VGKE-YDDNGNMASSGMVNEDLLNNLNAFEYYRKPYPKSLAN-DFGTDM   241
18      AYTAFDVCPANRVLNMLVAR-EGKE-YDAGGEMASGGRINEALLQQLAGLPYYSQPYPKSLAN-EFGTSE   240
19      KYIAFDVCPANRVLNMLAHD-AGKP-FDDAGQMAASGTLHPNLLKMLDELEYYELPYPKSLAN-DFGTDV   241
20      NYLAFDICAANRVLNMLTAK-EGKE-YDEGGKMAALGNVDNALLEKLNGLDFYKAAYPKSLAN-DFGTNI   240
21      SFLAFDICAANRVLNMLAAR-EGKE-YDEGGKMAASGNVDNALLAKLNGLDFYKEAYPKSLAN-DFGTNI   240
22      QYYAYDVCPANRVLNMLAAK-EKLA-FDEGGKLASSGKLYEPLLADLGRLDYYSMDYPKSLAN-DFGTDL   241
23      KYIAFDVCAANRVLNLLIKE-TGKA-YDDEGKMAATGIVQEALLAQLNKLDYYQQSFPKSLAN-DFGTGV   241
24      NFIAFDICPCNYILDALAET-LGKP-FDEDGKLAAGGVTDNALLESLNGLSYYGQDHPKTLAG-KFGTGT   241
25      AFAAFDICPCNYILDSLAAT-LGRA-YDEDGQLAAGGVTDAPLLESLNGLAFYRESYPRTIAS-KFGTGT   241
26      GVFVTDTGPGNTLLDAFARTHFQVE-YDKDALFAKQGTVHEILLSALKDHPFFKEPFPRTTGPELFSKAY   276
27      GIFVTDTGPGNTLLDAFARTYFQVE-YDQDAVLARQGRVNEVLLAALKDDSFFKAPFPRTTGPELFSKIY   276
28      DSFAVDVCPANQLLNLLANR-EGRA-FDRSGDMAKSGKVNTRLLKDLDDFDYYKKPFPKSLEM-DFGPEI   252
29      DSFAVDVCPANQLLNLLAEH-EGQT-FDNEGKIAAAGTVNSRLLKDLDGFTYYMKPLPKSLGV-DFGPEI   246
30      DSFAADVCPANQLLNLLAQR-ESKN-YDNEGEMASSGSINVRLLQDLDGFDYYKASFPKSLGV-DFGPEI   253

          190       200       210       220       230       240       250
          D192
```

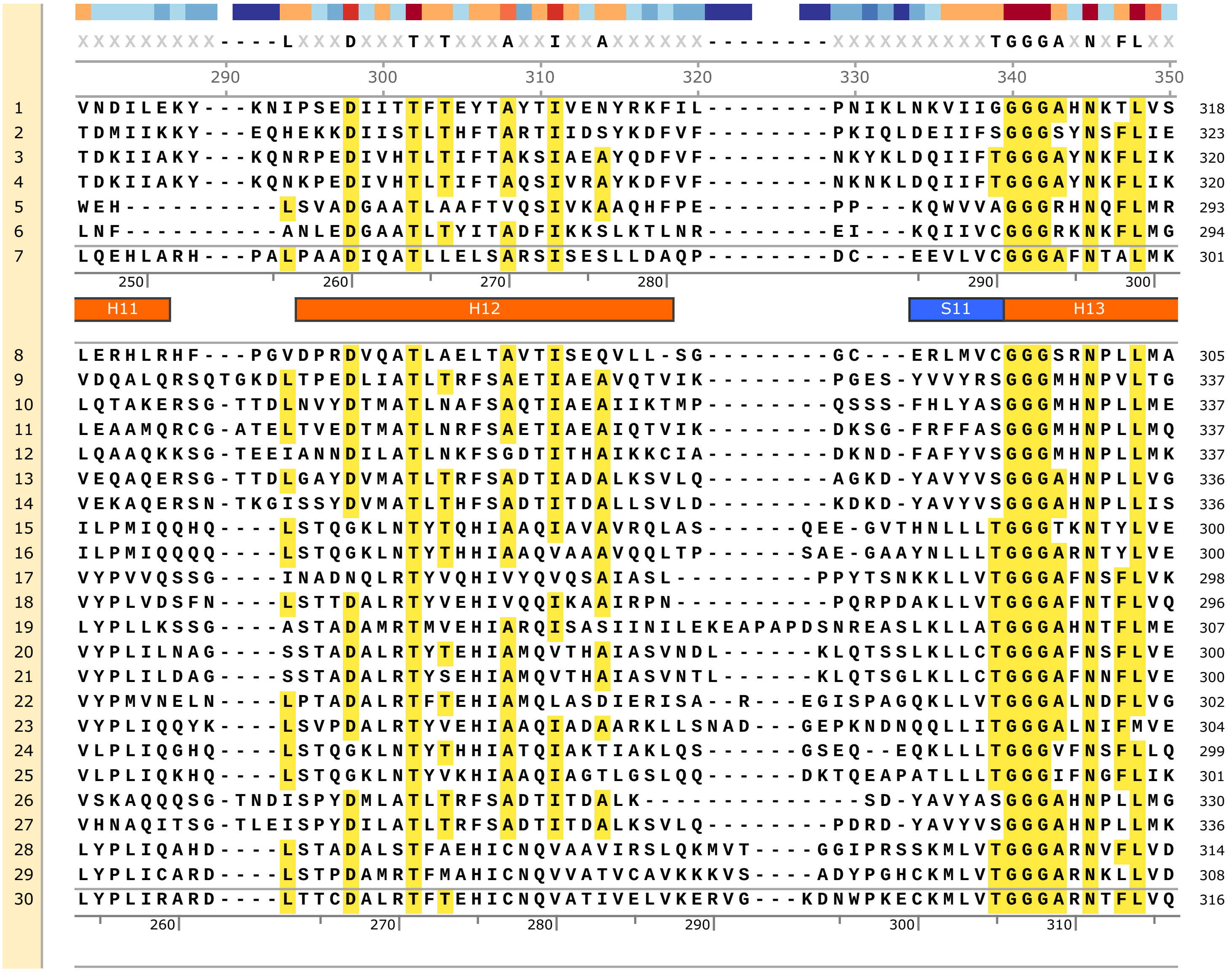
XXXXXXXXX----LXXXDXXXTXTXXXAXXIXXAXXXXXX--------XXXXXXXXXXTGGGAXNXFLXX
290 300 310 320 330 340 350
1 VNDILEKY---KNIPSEDIITTFTEYTAYTIVENYRKFIL--------PNIKLNKVIIGGGGAHNKTLVS 318
2 TDMIIKKY---EQHEKKDIISTLTHFTARTIIDSYKDFVF--------PKIQLDEIIFSGGGSYNSFLIE 323
3 TDKIIAKY---KQNRPEDIVHTLTIFTAKSIAEAYQDFVF--------NKYKLDQIIFTGGGAYNKFLIK 320
4 TDKIIAKY---KQNKPEDIVHTLTIFTAQSIVRAYKDFVF--------NKNKLDQIIFTGGGAYNKFLIK 320
5 WEH----------LSVADGAATLAAFTVQSIVKAAQHFPE--------PP---KQWVVAGGGRHNQFLMR 293
6 LNF----------ANLEDGAATLTYITADFIKKSLKTLNR--------EI---KQIIVCGGGRKNKFLMG 294
7 LQEHLARH--PALPAADIQATLLELSARSISESLLDAQP--------DC---EEVLVCGGGAFNTALMK 301
250 260 270 280 290 300
H11 H12 S11 H13
8 LERHLRHF---PGVDPRDVQATLAELTAVTISEQVLL-SG--------GC---ERLMVCGGGSRNPLLMA 305
9 VDQALQRSQTGKDLTPEDLIATLTRFSAETIAEAVQTVIK--------PGES-YVVYRSGGGMHNPVLTG 337
10 LQTAKERSG-TTDLNVYDTMATLNAFSAQTIAEAIIKTMP--------QSSS-FHLYASGGGMHNPLLME 337
11 LEAAMQRCG-ATELTVEDTMATLNRFSAETIAEAIQTVIK--------DKSG-FRFFASGGGMHNPLLMQ 337
12 LQAAQKKSG-TEEIANNDILATLNKFSGDTITHAIKKCIA--------DKND-FAFYVSGGGMHNPLLMK 337
13 VEQAQERSG-TTDLGAYDVMATLTRFSADTIADALKSVLQ--------AGKD-YAVYVSGGGAHNPLLVG 336
14 VEKAQERSN-TKGISSYDVMATLTHFSADTITDALLSVLD--------KDKD-YAVYVSGGGAHNPLLIS 336
15 ILPMIQQHQ----LSTQGKLNTYTQHIAAQIAVAVRQLAS------QEE-GVTHNLLLTGGGTKNTYLVE 300
16 ILPMIQQQQ----LSTQGKLNTYTHHIAAQVAAAVQQLTP------SAE-GAAYNLLLTGGGARNTYLVE 300
17 VYPVVQSSG---INADNQLRTYVQHIVYQVQSAIASL---------PPYTSNKKLLVTGGGAFNSFLVK 298
18 VYPLVDSFN----LSTTDALRTYVEHIVQQIKAAIRPN----------PQRPDAKLLVTGGGAFNTFLVQ 296
19 LYPLLKSSG----ASTADAMRTMVEHIARQISASIINILEKEAPAPDSNREASLKLLATGGGAHNTFLME 307
20 VYPLILNAG----SSTADALRTYTEHIAMQVTHAIASVNDL------KLQTSSLKLLCTGGGAFNSFLVE 300
21 VYPLILDAG----SSTADALRTYSEHIAMQVTHAIASVNTL------KLQTSGLKLLCTGGGAFNNFLVE 300
22 VYPMVNELN----LPTADALRTFTEHIAMQLASDIERISA--R---EGISPAGQKLLVTGGGALNDFLVG 302
23 VYPLIQQYK----LSVPDALRTYVEHIAAQIADAARKLLSNAD---GEPKNDNQQLLITGGGALNIFMVE 304
24 VLPLIQGHQ----LSTQGKLNTYTHHIATQIAKTIAKLQS------GSEQ--EQKLLLTGGGVFNSFLLQ 299
25 VLPLIQKHQ----LSTQGKLNTYVKHIAAQIAGTLGSLQQ------DKTQEAPATLLLTGGGIFNGFLIK 301
26 VSKAQQQSG-TNDISPYDMLATLTRFSADTITDALK--------------SD-YAVYASGGGAHNPLLMG 330
27 VHNAQITSG-TLEISPYDILATLTRFSADTITDALKSVLQ--------PDRD-YAVYVSGGGAHNPLLMK 336
28 LYPLIQAHD----LSTADALSTFAEHICNQVAAVIRSLQKMVT----GGIPRSSKMLVTGGGARNVFLVD 314
29 LYPLICARD----LSTPDAMRTFMAHICNQVVATVCAVKKKVS----ADYPGHCKMLVTGGGARNKLLVD 308
30 LYPLIRARD----LTTCDALRTFTEHICNQVATIVELVKERVG---KDNWPKECKMLVTGGGARNTFLVQ 316
260 270 280 290 300 310

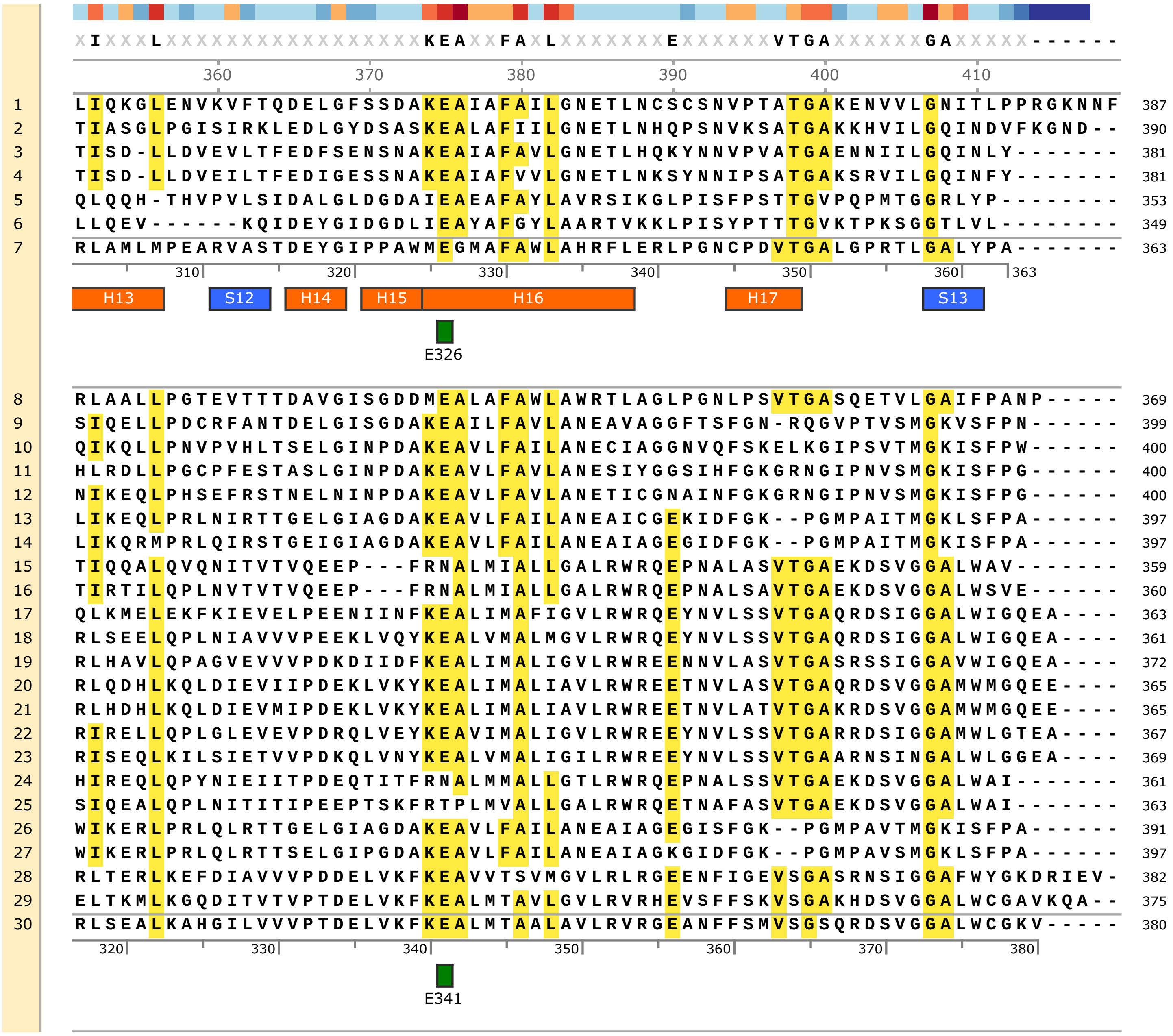
XIXXXLXXXXXXXXXXXXXXXXXKEAXXFAXLXXXXXXXXEXXXXXXVTGAXXXXXXXGAXXXXXX------
360 370 380 390 400 410
1 LIQKGLENVKVFTQDELGFSSDAKEAIAFAILGNETLNCSCSNVPTATGAKENVVLGNITLPPRGKNNF 387
2 TIASGLPGISIRKLEDLGYDSASKEALAFIILGNETLNHQPSNVKSATGAKKHVILGQINDVFKGND-- 390
3 TISD-LLDVEVLTFEDFSENSNAKEAIAFAVLGNETLHQKYNNVPVATGAENNIILGQINLY------- 381
4 TISD-LLDVEILTFEDIGESSNAKEAIAFVVLGNETLNKSYNNIPSATGAKSRVILGQINFY------- 381
5 QLQQH-THVPVLSIDALGLDGDAIEAEAFAYLAVRSIKGLPISFPSTTGVPQPMTGGRLYP-------- 353
6 LLQEV------KQIDEYGIDGDLIEAYAFGYLAARTVKKLPISYPTTTGVKTPKSGGTLVL-------- 349
7 RLAMLMPEARVASTDEYGIPPAWMEGMAFAWLAHRFLERLPGNCPDVTGALGPRTLGALYPA------- 363
310 320 330 340 350 360 363
H13 S12 H14 H15 H16 H17 S13
E326
8 RLAALLPGTEVTTTDAVGISGDDMEALAFAWLAWRTLAGLPGNLPSVTGASQETVLGAIFPANP----- 369
9 SIQELLPDCRFANTDELGISGDAKEAILFAVLANEAVAGGFTSFGN-RQGVPTVSMGKVSFPN------ 399
10 QIKQLLPNVPVHLTSELGINPDAKEAVLFAILANECIAGGNVQFSKELKGIPSVTMGKISFPW------ 400
11 HLRDLLPGCPFESTASLGINPDAKEAVLFAVLANESIYGGSIHFGKGRNGIPNVSMGKISFPG------ 400
12 NIKEQLPHSEFRSTNELNINPDAKEAVLFAVLANETICGNAINFGKGRNGIPNVSMGKISFPG------ 400
13 LIKEQLPRLNIRTTGELGIAGDAKEAVLFAILANEAICGEKIDFGK--PGMPAITMGKLSFPA------ 397
14 LIKQRMPRLQIRSTGEIGIAGDAKEAVLFAILANEAIAGEGIDFGK--PGMPAITMGKISFPA------ 397
15 TIQQALQVQNITVTVQEEP---FRNALMIALLGALRWRQEPNALASVTGAEKDSVGGALWAV------- 359
16 TIRTILQPLNVTVTVQEEP---FRNALMIALLGALRWRQEPNALSAVTGAEKDSVGGALWSVE------ 360
17 QLKMELEKFKIEVELPEENIINFKEALIMAFIGVLRWRQEYNVLSSVTGAQRDSIGGALWIGQEA---- 363
18 RLSEELQPLNIAVVVPEEKLVQYKEALVMALMGVLRWRQEYNVLSSVTGAQRDSIGGALWIGQEA---- 361
19 RLHAVLQPAGVEVVVPDKDIIDFKEALIMALIGVLRWREENNVLASVTGASRSSIGGAVWIGQEA---- 372
20 RLQDHLKQLDIEVIIPDEKLVKYKEALIMALIAVLRWREETNVLASVTGAQRDSVGGAMWMGQEE---- 365
21 RLHDHLKQLDIEVMIPDEKLVKYKEALIMALIAVLRWREETNVLATVTGAKRDSVGGAMWMGQEE---- 365
22 RIRELLQPLGLEVEVPDRQLVEYKEAVIMALIGVLRWREEYNVLSSVTGARRDSIGGAMWLGTEA---- 367
23 RISEQLKILSIETVVPDKQLVNYKEALVMALIGILRWREEYNVLSSVTGAARNSINGALWLGGEA---- 369
24 HIREQLQPYNIEIITPDEQTITFRNALMMALLGTLRWRQEPNALSSVTGAEKDSVGGALWAI------- 361
25 SIQEALQPLNITITIPEEPTSKFRTPLMVALLGALRWRQETNAFASVTGAEKDSVGGALWAI------- 363
26 WIKERLPRLQLRTTGELGIAGDAKEAVLFAILANEAIAGEGISFGK--PGMPAVTMGKISFPA------ 391
27 WIKERLPRLQLRTTSELGIPGDAKEAVLFAILANEAIAGKGIDFGK--PGMPAVSMGKLSFPA------ 397
28 RLTERLKEFDIAVVVPDDELVKFKEAVVTSVMGVLRLRGEENFIGEVSGASRNSIGGAFWYGKDRIEV- 382
29 ELTKMLKGQDITVTVPTDELVKFKEALMTAVLGVLRVRHEVSFFSKVSGAKHDSVGGALWCGAVKQA-- 375
30 RLSEALKAHGILVVVPTDELVKFKEALMTAALAVLRVRGEANFFSMVSGSQRDSVGGALWCGKV----- 380
320 330 340 350 360 370 380
E341

**Figure S8. Alignment of selected protein sequences of AnmK**

Alignment of representative AnmK protein sequences, including three tick sequences (*O. moubata*, *R. sanguineus* and *I. ricinus* - IrAnmK). The alignment corresponds to the sequences of the phylogenetic tree displayed in Fig. S6, comprising bacterial sequences from the expected donor group (Chitinophagaceae - clade 2), and other sequences from Chitinophagaceae - clade 1, tick endosymbionts, and free-living bacteria. Above, conservation of residues across the whole set of sequences, shown as a heat-map. Below the reference sequence 7 (from *Pseudomonas aruginosa*), are shown structural predictions : S for sheet and H for Helix. For both this sequence and sequence 30 (IrAnmK), are also shown residues determined as functionally important (e.g. the catalytic site, at D182 in *P. aruginosa* numbering), as in Fig. S6.

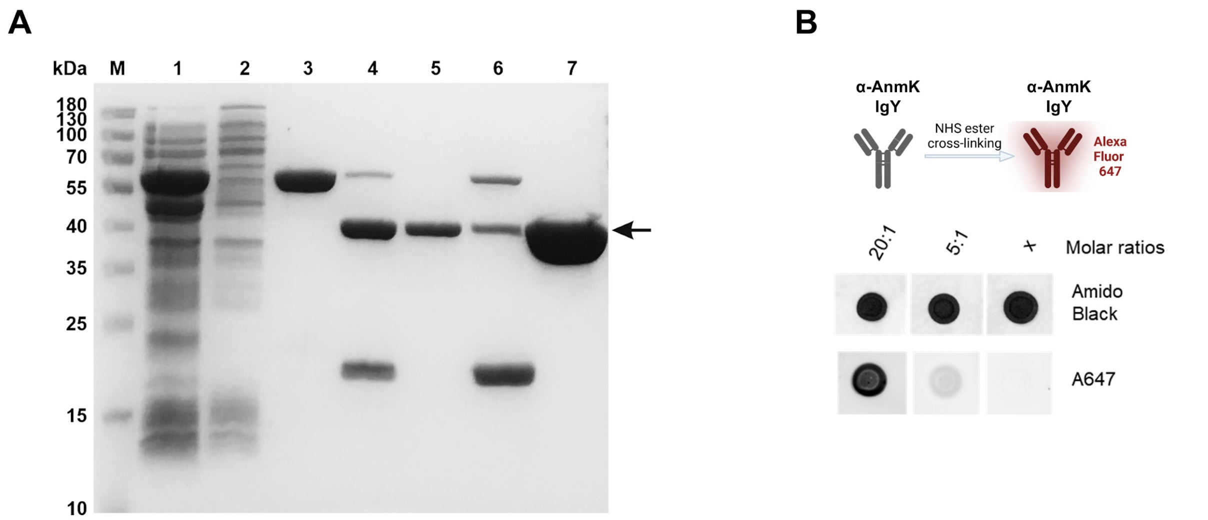


**Figure S9. Recombinant expression and purification of tick AnmK and fluorophore coupling to antibodies.**

**A** Purification of recombinant AnmK proteins (pET SUMO) from the soluble extract of BL-21(DE3) *E. coli* cells. Reducing SDS-PAGE showing *I. ricinus* AnmK at the different steps of purification. Lanes: M - Protein molecular weight marker, 1 - Soluble fraction of bacterial lysate, 2 - The flow-through fraction of the bacterial lysate from the nickel IMAC column, 3 - Elution fraction from the column by 0.5M imidazole, 4 - Elution fraction cleaved with SUMO protease and loaded onto the nickel IMAC column, 5 - Flow-through (cleaved protein), 6 - Control non-used elution with 0.5M imidazole, 7 - Final preparation of the protein (flow-through, lane 5) after concentration with Amicon centrifugal filter unit (10K MWCO) concentration. **B** A schematic of production of anti *I. ricinus* Anmk antibodies of IgY class in hens. IgY antibodies were isolated from chicken sera and conjugated to Alexa Fluor 647 using varying molar ratios. **C** Dot blot analysis was performed to evaluate the coupling efficiency of the fluorescent labeling.

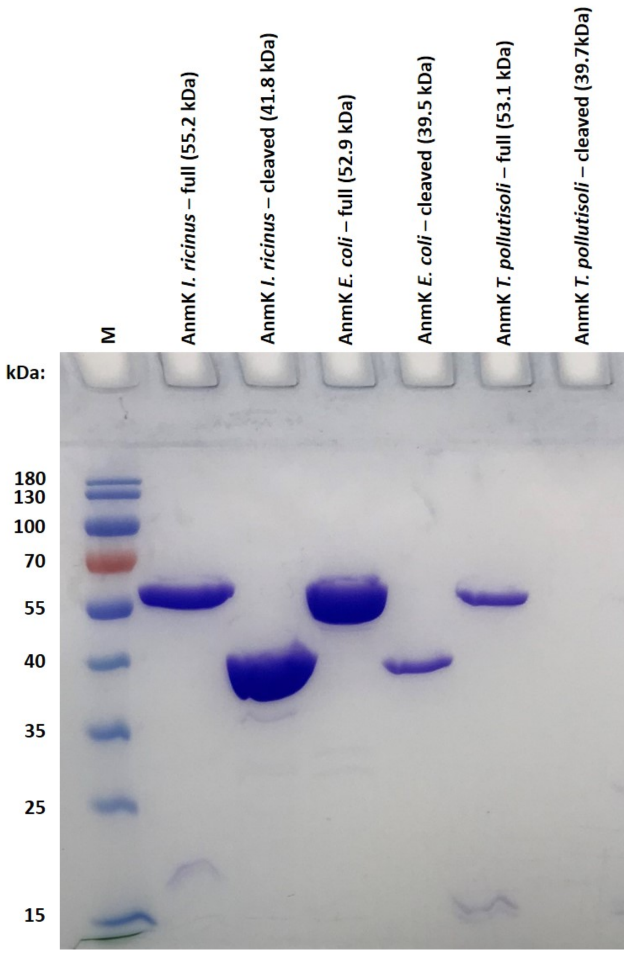


**Figure S10. Recombinant expression of AnmK homologs from *I. ricinus, E. coli,* and *T. pollutisoli*.**
*T. pollutisoli* represents the likely donor group of *anmK* to ticks. A reducing SDS-PAGE of purified AnmK recombinant proteins expressed in *E. coli* as fused proteins with SUMO tag. M - Protein molecular weight marker. Proteins are shown as purified from flow-through after SUMO cleavage using $Ni^{2+}$-IMAC affinity chromatography.

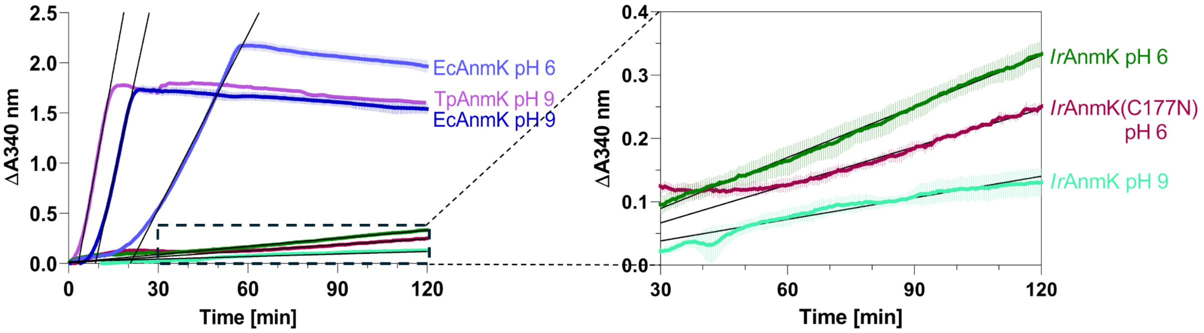


**Figure S11. Coupled enzyme analysis of anhydro-N-acetylmuramic acid kinase activity.**

AnmK activity was assessed by coupling ADP production to the conversion of PEP to pyruvate and then lactate by the sequential activity of pyruvate kinase (PK) and lactate dehydrogenase (LDH) –which simultaneously oxidizes NADH to NAD+ resulting in a decrease of absorbance at 340 nm. Shown are the reactions of AnmKs from *E. coli* (EcAnmK), *I. ricinus* (IrAnmK) and *T. pollutisoli* (TpAnmK) at pH 6 and 9 as well as the IrAnmK variant (C177N) at pH 6 used to calculate the reaction rate constants of linear regression equation (slope) as shown in the time courses.

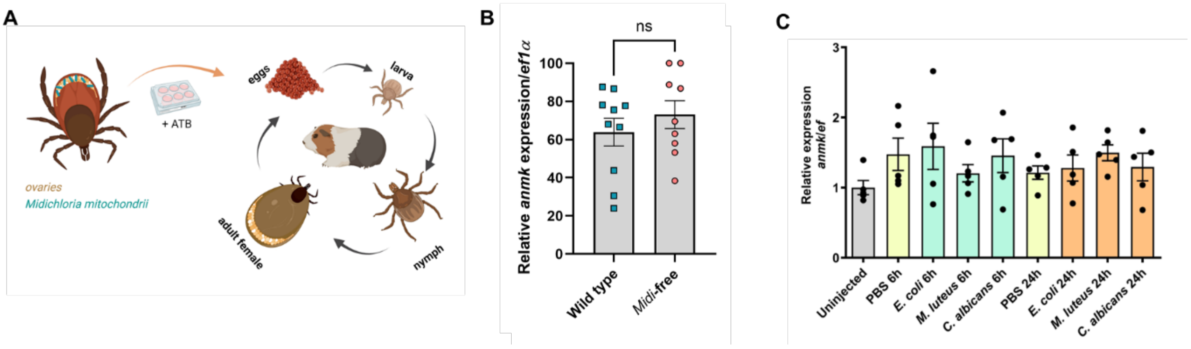


**Figure S12. Expression of *IranmK* is not altered in symbiont-free *I. ricinus* ticks or after injection of model microbes in the hemolymph .**

**A** Schematic overview of the steps leading to the generation of a *Midichloria*-free strain of *I. ricinus* ticks. **B** Relative expression (qRT-PCR) of *anmK* in *I. ricinus* females ovaries of wild-type ticks (control) and the symbiont-free line[67], denoted *Midi*-free. All bar graphs show mean and SEM, each dot representing a single female data. **C** Relative expression (qRT-PCR) of *anmK* in unfed nymphs 6 and 24 hours after injection of microbes. Each dot represents a pool of ten nymphs.

### Supplementary Tables

**Table S1: Horizontal Gene Transfer candidates in the genome of *I. ricinus*.**

Columns specify Detection method, Gene ID (GenBank accession between brackets), AvP parameters and results. Detection method: the search of candidates was primarily performed with the tool AvP, while a separate gene search was conducted for genes similar to *dae2* in *I. scapularis*. AvP parameters: “ingroup”, “recipient group” or “EGP” (as described in **Fig. S1**), Alien Index (AI) is indicated - stringency threshold-, identified donor group, description of the gene function, and localization (scaffold) in the genome of *I. ricinus*. Gene IDs followed by an asterisk correspond to genes for which a phylogenetic study did not support horizontal transfer (this was the case for all candidates with a vertebrate donor). Of note, due to a gap in the genome assembly of *I. ricinus* in the coding region of *dae2-1*, this gene model was not complete; therefore, for this gene, we indicated an accession corresponding to a transcriptome contig and comprising the full coding sequence.

| **Detection** | ***I. ricinus* gene ID (nr accession)** | **Ingroup** | **EGP** | **AI threshold** | **Donor group** | **Description** | **Scaffold** |
|---|---|---|---|---|---|---|---|
| AvP | IricT00015853 (CAN7961235.1) | Arthropoda | Acari | 40 | Phlebovirus (Bunyavirales) | Nucleocapsid protein N | Scaffold_5 |
| | IricT0000162 (CAN7953880.1) | Arthropoda | Acari | 40 | Phlebovirus (Bunyavirales) | Nucleocapsid protein N | Scaffold_1 |
| | IricT00021393 (CAN7966687.1) | Arthropoda | Acari | 40 | Phlebovirus (Bunyavirales) | Nucleocapsid protein N | Scaffold_9 |
| | IricT00021282 (CAN7966573.1) | Arthropoda | Acari | 40 | Phlebovirus (Bunyavirales) | Nucleocapsid protein N | Scaffold_9 |
| | IricT00012740 (CAN7958109.1) | Arthropoda | Acari | 40 | Phlebovirus (Bunyavirales) | Nucleocapsid protein N | Scaffold_4 |
| | IricT00015852 (CAN7961234.1) | Arthropoda | Acari | 40 | Nairovirus (Bunyavirales) | Nucleocapsid protein N | Scaffold_5 |
| | IricT00008813 (CAN7948355.1) | Arthropoda | Acari | 40 | Nairovirus (Bunyavirales) | Nucleocapsid protein N | Scaffold_16427 |
| | IricT00012932 (CAN7958306.1) | Arthropoda | Acari | 40 | Nairovirus (Bunyavirales) | Nucleocapsid protein N | Scaffold_4 |
| | IricT0001324 | Arthropoda | Acari | 40 | Totivirus | Hypothetical protein | Scaffold_ |

| | | | | | | | |
|---|---|---|---|---|---|---|---|
| | 4 (CAN7958619.1) | da | | | (Ghabrivirales) | | 4 |
| | IricT00003442 (CAN7969251.1) | Arthropoda | Acari | 40 | Densovirus (Piccovirales) | Non structural protein 1 (NS1) | Scaffold_10 |
| | IricT00020676 (CAN7965990.1) | Arthropoda | Acari | 40 | Rhabdovirus (Mononegavirales) | Nucleocapsid protein N | Scaffold_8 |
| | IricT00015847 (CAN7961229.1) | Arthropoda | Acari | 20 | Rhabdovirus (Mononegavirales) | Nucleocapsid protein N | Scaffold_5 |
| | IricT00015848 (CAN7961230.1) | Arthropoda | Acari | 20 | Rhabdovirus (Mononegavirales) | Nucleocapsid protein N | Scaffold_5 |
| | IricT00008830 (CAN7947248.1) | Arthropoda | Acari | 20 | Rhabdovirus (Mononegavirales) | Nucleocapsid protein N | Scaffold_1793 |
| | IricT00015509* (CAN7960882.1) | Arthropoda | Acari | 40 | Vertebrates | Translocator protein | Scaffold_5 |
| | IricT00013518* (CAN7958897.1) | Arthropoda | Acari | 40 | Vertebrates | 3-oxoacyl[acyl carrier protein] synthase | Scaffold_4 |
| | IricT00019949* (CAN7965248.1) | Arthropoda | Acari | 20 | Vertebrates | Calcium activated chloride channel regulator 1 | Scaffold_8 |
| | IricT00013998 (CAN7959379.1) | Arthropoda | Acari | 40 | Bacteria | anmK (Anhydro-N-acetylmuramic acid kinase) | Scaffold_4 |
| | IricT00008184 (CAN7973976.1) | Arthropoda | Acari | 20 | Bacteria | GNAT N acetyltransferase | Scaffold_14 |
| Other | IricT00022977 (GIDG01008499.1) | - | - | - | Bacteria | dae2-1 (similar to tae2 bacterial toxin) | Scaffold 7 |
| | IricT00001882 (CAN7964150.1) | - | - | - | Bacteria | dae2-2 (similar to tae2 bacterial toxin) | Scaffold 7 |
| | IricT00021525 (CAN7966821.1) | - | - | - | Bacteria | dae2-3 (similar to tae2 bacterial toxin) | Scaffold 9 |
| | IricT00020836 (CAN7966150.1) | - | - | - | Bacteria | dae2-4 (similar to tae2 bacterial toxin) | Scaffold 8 |

### Supplementary text

i) Sequences of dsRNA used in this study:

*Ixodes ricinus*

>AnmK_transcript [IricT00013998]

GCAGATGACCACCACACAGCTGGGAGACGGTGCGACCGTTGCAGCTGTTACCGG
TCTTCCGACGGTCAGCGACTTCCGTTCCGTAGACGTTGCACTGATGGGTCAGGGC
TCGCCCTTCTCCATAGCTGCTGAGCGCAGGCTCTTTGGCACAGAGTACCAGTTCT
TCCTCAACATCGGTGGGATGGCTTGCCTCACCTACGTCGGCAAAGGGAACGATGA
CAGCTTCGCGGCAGACGTATGCCCTGCGAATCAACTGCTCAACCTCCTAGCCCAA
CGGGAATCCAAGAACTATGATAACGAGGGCGAGATGGCATCTTCGGGATCCATCA
ACGTTCGTCTGCTTCAGGACCTGGATGGTTTTGACTACTACAAGGCCTCGTTTCC
GAAGTCGTTGGGCGTGGACTTTGGACCCGAGATC

*Haemaphysalis longicornis*

>GJUD01006560.1 TSA: Haemaphysalis longicornis HaelonEVm009155t1, transcribed RNA sequence

ACAGATGACAACAACGCAAGTCGGAGATGGTGCTGCCGTTGCTGCCGTAACTGGA
CTTCCCACAGTCAGTGACTTCAGATCAGTCGACGTCGCCCTGCAAGGTCAGGGAT
CTCCCTTCTCCTTAGCAGCAGAGAGGAAGCTGTTTGGCAGTGAATACCAATACTT
CCTCAACATAGGTGGCATGGCCTGCCTGACATACGTCGGCAAGAACGACGAAGAT
AGCTTCGCCTTGGACGTCTGCCCCGCCAACCAGCTTTTAAACCTCCTTGCTAAGC
AAGAAGGGTGCCATTACGACGACGATGGAAAGATAGCTGCGTCGGGGAAAGTCG
TTCCCAGGCTCCTCAAAGACCTAGACGGATTTGCCTACTACCAGAAGGCTTTGCC
GAAATCCTTGGGCGTGGACTTCGGCCCCGAAATCC

*Rhipicephalus sanguineus*

>XM_037664325.1:177-1304 PREDICTED: Rhipicephalus sanguineus anhydro-N-acetylmuramic acid kinase (LOC119396938), mRNA

TCAGCTGACAACTGCACAAGTCGGGGATGGCGCTGCTGTTGCTGCTGTAACCGGA
CTGCCCACTGTCAGCGACTTCAGATCGGTGGACATTGCTCTGGAAGGACAAGGCT
CCCCATTCTCGCTGGCTGCCGAGAGGAAGCTCTTTGGCAGCGAATACCAGTACTT
CCTGAACATAGGAGGCATGGCATGCCTAACGTACGTAGGCAAATGCAACGACGAC
AGCTTTGCTGTGGACGTTTGTCCTGCCAACCAGCTTTTGAACCTCCTCGCCGAGC
ACGAGGGTCAAACTTTCGACAATGAAGGAAAGATAGCAGCTGCGGGAACAGTTAA
CAGCCGACTACTGAAGGACCTCGATGGTTTCACCTACTACATGAAACCTCTGCCC
AAATCTCTCGGCGTAGACTTTGGCCCAGAAATC

2) Sequences of oligo DNA (PCR primers) used in this study:

Iric_AnmK_F: TGTCCAACGTCTGTCCGAAG

Iric_AnmK_R: TGAACTTGACCAGCTCGTCC

Iscap_AnmK_F: TGGCCCAAGGAATGCAAGAT

Iscap_AnmK_R: GACAGACGCTGGACAAGGAA

Hlong_AnmK_F: CACTGCAACATGCTGGTCAC

Hlong_AnmK_R: ATGTCTTGCTCTCGGAGTGC

Rsang_AnmK_F: TACTGTTACCGTGCCGACAG

Rsang_AnmK_R: AAGAAGCTGACCTCGTGCC

Omoub_AnmK_F: AAGATGCTCGTCACTGGAGG

Omoub_AnmK_R:CAAACTCTTTGAGCCGCTCC

F1(anmk_Iric): CGGTGATCGCCATCATCATTAC

R1(anmk_Iric): GTTCCTCAGCGACGAAGACG

F2(anmk_Iric): CGGCGCTTGTGGATCAACG

R2(anmk_Iric): CGTGCGACTGCTCGGATCTG

Iric_EF_F: TGTCCAACGTCTGTCCGAAG

Iric_EF_R: GGACGAGCTGGTCAAGTTCA

Iscap_EF_F: TGTCCAACGTCTGTCCGAAG

Iscap_EF_R: GGACGAGCTGGTCAAGTTCA

Hlong_EF_F: GTGACAACGTGGGCTTCAAC

Hlong_EF_R: TGGAGTCTCCACACACGTAC

Rsang_EF_F: CGTCGGCTTCAACGTCAAGA

Rsang_EF_R: CCTTCGAGTCTCCACACACA

Omoub_EF_F: TCGAGATGCATCACGAGGC

Omoub_EF_R: CGTTCTTGACGTTGAAGCCC

16S_F: TCCTACGGGAGGCAGCAGT

16S_R: GGACTACCAGGGTATCTAATCCTGTT

Midi_GyrB_F: CTTGAGAGCAGAACCACCTA

Midi_GyrB_R: CAAGCTCTGCCGAAATATCTT
